# *De novo* acyl carrier proteins display structure-independent modification and sequence novelty

**DOI:** 10.1101/2025.06.07.658270

**Authors:** Michael A. Herrera, Grace K. King, Zoe Ozols, Gioele A. Tiburtini, Nicoletta Schiavo, Francesca Spyrakis, Louise K. Charkoudian, Dominic J. Campopiano

## Abstract

Acyl carrier proteins (ACPs) are dynamic, structurally conserved α-helical proteins central to many primary and secondary metabolic processes. Whilst prior engineering efforts have focused on strategic mutagenesis and “helix swaps”, much of the ACP sequence design space remains underexplored. Here, we create diverse variants of the archetypal ACP subclass – AcpP – using a bespoke sequence-generating algorithm (ALGO-CP), which utilises a combined evolutionary and physicochemical design approach. Using ALGO-CP, we generated two soluble candidates – ALGO-055 and ALGO-059 – that can undergo full post-translational modification from *apo*→*holo*→acyl forms *in vitro*, using recombinant modifying enzymes. Building on these successful designs, we further adapted ALGO-CP to produce several chimeras, two of which – ^ch^ALGO-012 and ^ch^ALGO-024 – also exhibit full modifiability. We explore the structural plasticity of our ALGO variants via robust molecular dynamic simulations, and we further reveal by circular dichroism spectroscopy that ALGO-055 and ALGO-059 lack the canonical α-helical fold of an ACP, whilst remaining soluble and readily modifiable. Upon acylation of ALGO-055 and ALGO-059, we observe a marked increase in helicity indicative of partial restructuring. Of note, both ALGO-055 and ALGO-059 harbour several rare amino acid variations across their sequences, whilst preserving many important acidic “hotspots” involved in key protein-protein interactions. By testing the limits of the AcpP design space, our findings suggest that some key aspects of ACP behaviour (specifically post-translational modification) can be retained independently of the canonical structure. This work establishes a foundation for probing ACP sequence diversity through a hybrid computational-experimental approach. ALGO-CP is available under AGPL3.0 license: https://github.com/MAHerrera-94/ALGO_CP.

## Introduction

Acyl carrier proteins (ACPs), and their related peptidyl carrier proteins (PCPs), are small (8-10 kDa) and ubiquitous proteins that drive metabolic flux within cells.^1-4^ These highly dynamic proteins work by activating and shuttling metabolic substrates between enzymes within biosynthetic pathways/assemblies. Their function depends on the precise recognition of enzyme partners through transient and dynamic protein–protein interactions (PPIs), with each cognate partner evolving a complementary interface for ACP/PCP docking and substrate transfer. In addition to essential lipids and vitamins, countless polyketide and non-ribosomal peptide pharmaceuticals originate from ACP/PCP-driven biosynthesis including glycopeptide antibiotics,^5^ avermectins^6^ and statins.^7^ Without the precision and versatility of these proteins, such indispensable molecules are not possible in nature nor in the clinic.

ACPs are remarkably diverse at the sequence level, with many homologues sharing as little as ∼20– 30% identity. Despite this, ACPs generally conserve a compact helical fold, comprising four antiparallel helices (αI-IV) connected by flexible loops (Fig. 1A). For an ACP to be functionally competent, it must undergo post-translational modification (PTM) to its *holo-* form via the attachment of a 4′-phosphopantetheinyl (4’-PP) prosthetic group, derived from coenzyme A (CoASH, Fig. 1B). This PTM is catalysed by a 4’-phosphopantetheinyl transferase (4’-PPTase), which fixes the 4’-PP group to an invariant serine residue situated before helix αII. In this *holo-* form, substrates are covalently tethered to the terminal thiol group of the 4’-PP; bound lipophilic substrates can be sequestered within the hydrophobic cavity of the ACP, thereby protecting it from premature hydrolysis or undesired side-reactions (Fig. 1C).^8, 9^ The surface of the ACP mediates interactions with partner enzymes, with solvent-exposed residues along helix αII (often called the “universal recognition helix”) playing a major role in PPIs. The nature and precision of these PPIs is governed by the ACP surface charge distribution, hydropathy and topography, as evidenced by numerous structural and NMR analyses of ACP complexes.^10-17^ Like many ancient enzyme families, such as the short-chain dehydrogenase/reductase,^18^ it is widely held that ACPs are robust to sequence variation, provided these key structural, functional and biophysical features are retained.

**Fig. 1.**
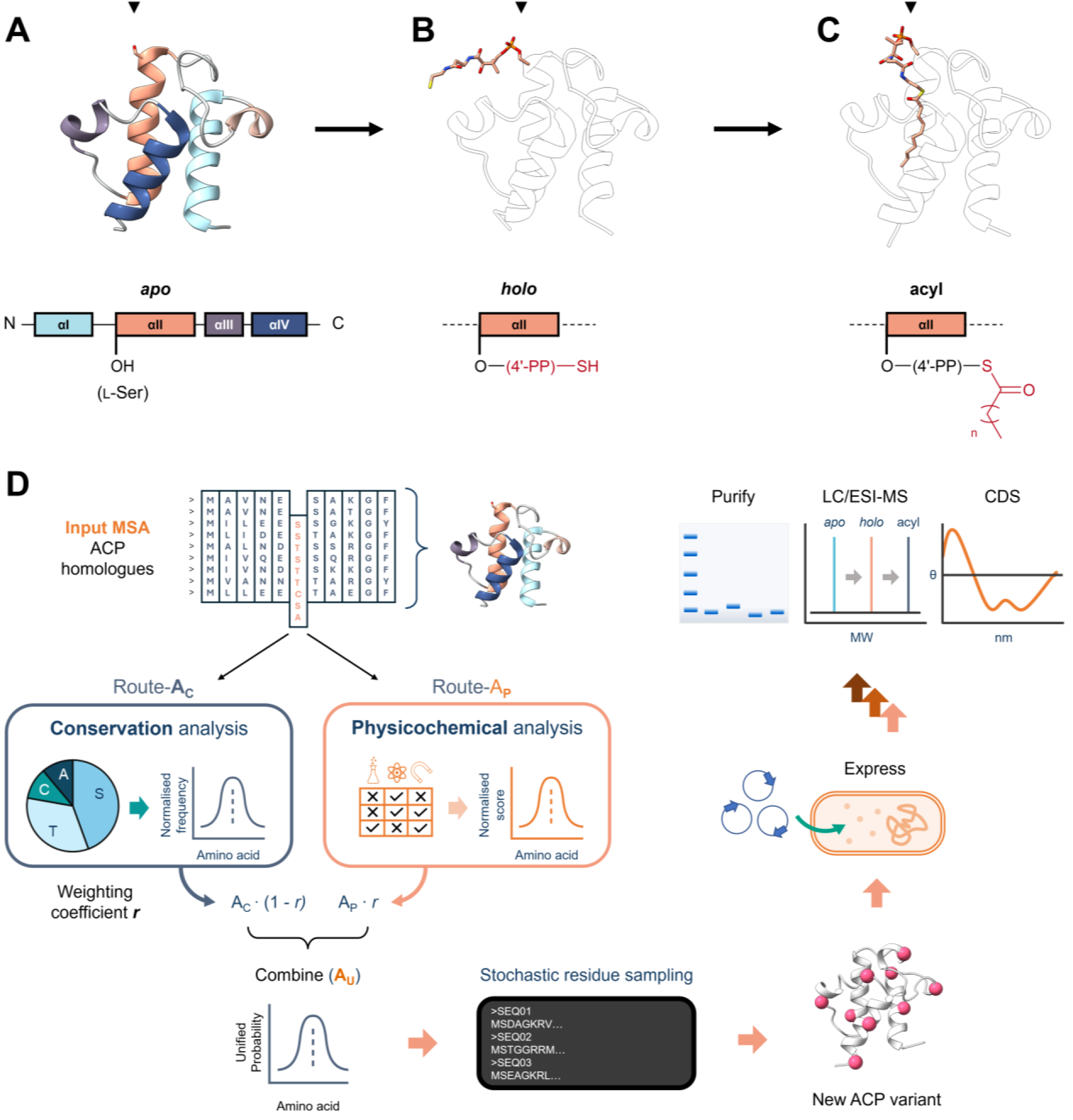
(A-C) Annotated schematic of an ACP, with colour-coded α-helices, in its *apo* (A), *holo* (B) and acyl (C) state. (PDB: 1T8K, 5H9H and 2FAE). Black triangles indicate the position of the invariant, essential serine. (D) Graphical summary of ALGO-CP and subsequent wet-lab analysis of *de novo* ACP designs, starting with a multiple sequence alignment (MSA) of ACP homologues. Using this MSA as a template, ALGO-CP utilises position-specific conservation and physicochemical analysis to generate new ACP-like sequences. The influence of evolutionary conservation vs. physicochemical analysis can be tuned via a weighting coefficient *r*, which ranges from 0 to 1. Randomly sampled ACP-like sequences are expressed, purified and examined for PTM from their inactive *apo*-form to functional *holo*- and acyl-forms. Successful candidates are then characterised by circular dichroism (CD) spectroscopy.

Efforts to engineer ACPs include rational mutagenesis^19, 20^ and the creation of chimeras,^21, 22^ in which helical segments are exchanged between natural homologues to introduce new functions and/or re-route biosynthetic pathways. Whilst such strategies have produced functional variants, they explore only a narrow corner of ACP design space, leaving many novel ACP designs – and functions – out of reach. A more rigorous exploration of this sequence space could deepen our understanding of ACP evolution and uncover new insights for engineering. Computational biology is well-placed to address this challenge, yet to the best of our knowledge, no such strategy has been applied to the design of new ACPs.

In this work, we developed a simple and tuneable sequence generation algorithm (ALGO-CP) designed to map the sequence space of ACPs. Rather than strictly relying on phylogenetic constraints, ALGO-CP takes a softened design approach that blends evolutionary conservation with local physicochemical information. In theory, this would permit the controlled sampling of amino acids that are rarely observed in natural homologues, thereby pushing the limits of natural sequence diversity. Using the prokaryotic housekeeping ACP (AcpP) subclass as a template, we applied ALGO-CP to generate diverse *de novo* ACP-like sequences, two of which – ALGO-055 and ALGO-059 – could undergo PTM to their *holo*- and acylated forms *in vitro*. By re-configuring ALGO-CP, we created two soluble chimeric variants of ALGO-055 and ALGO-059 – named ^ch^ALGO-012 and ^ch^ALGO-024 – that also undergo complete PTM. We further reveal by circular dichroism (CD) spectroscopy that ALGO-055 and ALGO-059 are modifiable despite lacking the canonical α-helical structure of an ACP, and we report evidence of structural recovery upon acylation. We propose that these unique properties arise from rare amino acid combinations not featured in natural AcpP homologues, alongside the preservation of key acidic regions necessary for productive PPIs.

## Results and Discussion

### ALGO-CP concept and summary

ALGO-CP is a tuneable sequence generation algorithm designed to explore the boundaries of the ACP sequence design space (Fig. 1D). It operates on a multiple sequence alignment (MSA) of constituent amino acids using two complementary sub-algorithms, route-**A**_**C**_ and route-**A**_**P**_, to capture position-specific conservation and physicochemical information, respectively. The physicochemical properties examined – isoelectric point (pI), Kyte–Doolittle hydropathy (H_KD_), and van der Waals volume (vdW, in Å^3^) – are important for ACP structure, solubility and interaction specificity. This information can be combined via a weighting coefficient *r*, ranging from 0.00 (purely **A**_**C**_-focused) to 1.00 (purely **A**_**P**_-focused), to generate a unified amino acid probability profile (**A**_**U**_) for each amino acid position, from which new residues are sampled stochastically to construct a new ACP-like sequence (see Experimental for full details). By design, ALGO-CP preserves highly conserved residues (such as the essential, modifiable serine), whilst allowing greater diversity at variable positions. Sequences that are generated exclusively via route-**A**_**C**_ are constrained to observable substitutions within the input MSA. In contrast, route-**A**_**P**_ promotes novelty by relaxing evolutionary constraints, instead evaluating the physicochemical profile of each alignment block against all 20 proteinogenic amino acids. In theory, this would enable the incorporation of many rare, yet physiochemically compatible, amino acid variations. We reasoned that a calibrated combination of these two design strategies would allow the exploration of ACP sequence space in a controlled and interpretable manner. To test this, we required a reliable and well-characterised template for the design of new ACPs.

### Designing AcpP variants using ALGO-CP

The archetypal ACP, known as AcpP, is essential in prokaryotes.^23^ With an estimated 350,000 copies per cell, *Escherichia coli* AcpP (*Ec*AcpP) can interact with >25 other proteins to synthesise several life-critical lipids (fatty acids, phospholipids, lipid A, lipoic acid, biotin),^10, 24-26^ mediate nutrient stress responses (via communication with SpoT)^27, 28^ and promote chromosome organisation (via interaction with the chromosome partition protein MukB).^29^ The multi-functional “pocket knife” versatility of AcpP, together with the sheer abundance of sequence homologues, makes it an ideal starting template for *de novo* ACP design using ALGO-CP.

To seed the design process, we first retrieved 2558 AcpP homologues from UniRef90 using 17 validated AcpP sequences as queries (Table S3). This dataset was cleaned to remove duplicate and incomplete sequences, bringing the new total to 2167. These homologues were aligned using MAFFT,^30^ and the resulting MSA was used as input for ALGO-CP. We configured ALGO-CP to generate sets of 5000 unique sequences, with each iteration raising the weighting coefficient *r* from 0.00 to 1.00 in increments of 0.05. For each set, we computed the mean positional values of H_KD_, pI, and vdW volume, and compared these to the input MSA via pairwise correlation (Fig. S1). As anticipated, the sequences generated by ALGO-CP became more divergent from the input MSA with increasing *r*, reflecting a shift towards sequence novelty permitted by the softer computational strategy of route-**A**_**P**_ (Fig. 2A).

**Fig. 2.**
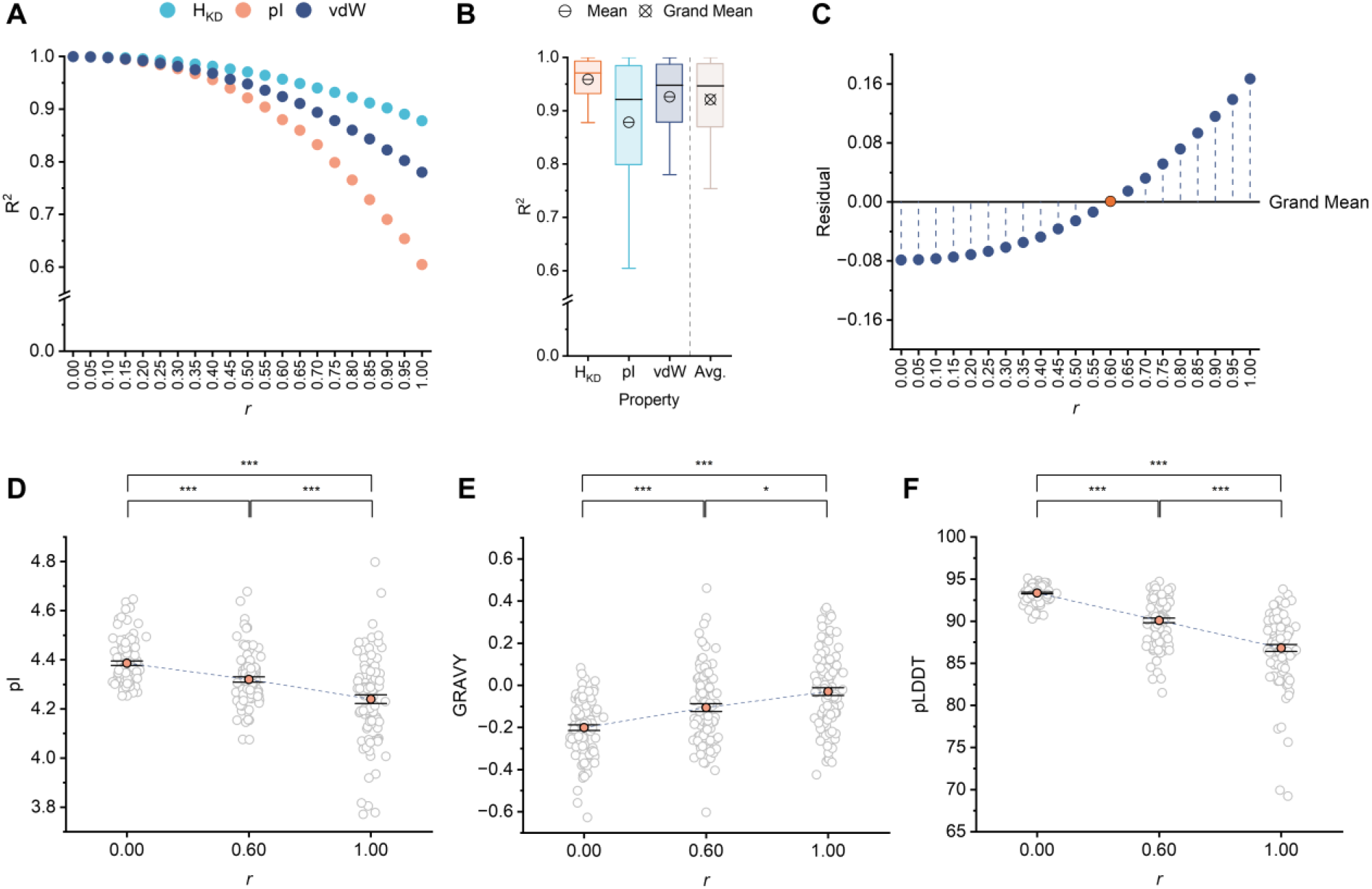
(A) Positional, pairwise property correlations (R^2^) between ALGO-CP sequences and the input MSA of natural AcpP homologues, with varying weighting coefficient *r*. Lower R^2^ indicates greater overall physicochemical deviation from natural AcpP homologues. (B) The spread and central tendency of individual (H_KD_, pI, vdW volume) and averaged (Avg.) pairwise property correlations across all ALGO-CP sequence sets (*r* = 0-1, n = 20). Median values are shown as black bands. Boxes represents the interquartile range (IQR). Whiskers extend to data points up to 1.5 x the IQR from the lower and upper quartiles. The crossed circle denotes the grand mean across all sets and properties. (C) Residual analysis of averaged pairwise property correlations relative to the grand mean. Negative residuals reflect sequences with more conservative physicochemical profiles, whereas positive residuals indicate more divergent profiles. Based on its proximity to the grand mean, sequences generated using *r* = 0.60 (orange circle) were taken to represent the ideal balance between sequence conservation vs. exploration. (D) Mean pIs of ALGO-designed sequences generated using *r* = 0.00, 0.60 or 1.00. Each datapoint (grey circle) represents a unique sequence. Error bars represent 95% confidence interval (n = 100). (***) = p ≤ 0.001 (Welch’s ANOVA with Games-Howell post-hoc test). (E) Mean Grand Average of Hydropathy (GRAVY) values of ALGO-designed sequences generated using *r* = 0.00, 0.60 or 1.00. Each datapoint (grey circle) represents a unique sequence. Error bars represent 95% confidence interval (n = 100). (*) = p ≤ 0.05; (***) = p ≤ 0.001 (Welch’s ANOVA with Games-Howell post-hoc test). (F) Mean Predicted Local Distance Difference Test (pLDDT) of ALGO-designed sequences, folded using AlphaFold3, generated at *r* = 0.00, 0.60 or 1.00. Each datapoint (grey circle) represents a unique sequence folded by AlphaFold3. Error bars represent 95% confidence interval (n = 100). (***) = p ≤ 0.001 (Welch’s ANOVA with Games-Howell post-hoc test).

To approximate the balance point between the two sub-algorithms, we averaged the pairwise property correlations within each sequence set and analysed their residual proximity to the grand property mean across all sequence sets (Fig. 2B-C). Negative residuals indicate relatively conservative sequences, whereas positive residuals indicate relatively divergent sequences. Based on this proximity analysis, we estimated that sequences generated using *r* = 0.60, which returned the smallest residual value (8.75 ×10^−4^), may best reflect this strategic balance between conservative vs explorative sequence design. A subset of 100 randomly-sampled, unique sequences generated at *r* = 0.60 were found to exhibit intermediate overall sequence properties relative to those sampled from pure **A**_**C**_-and **A**_**P**_-guided designs (Fig. 2D-E, Tables S4-5, S7-8, S10-11). Furthermore, this subset could be accurately folded using AlphaFold3^31^ (pLDDT = 90.1 ± 2.75), suggesting that they may adopt ACP-like folds if expressed in soluble form (Fig. 2F, Tables S6, S9, S12). However, since ALGO-CP does not explicitly predict solubility or biochemical function, we proceeded to characterise a sample of these sequences experimentally.

### Expression, purification and PTM of ALGO sequences

The ability to undergo functional PTM is an essential hallmark of all naturally occurring ACPs. For *in vitro* study, recombinant PTM enzymes can efficiently convert *apo*-ACPs to their functionally competent *holo*- and acyl-forms. *Bacillus subtilis* Sfp (*Bs*Sfp)^32^ or *E. coli holo*-ACP-synthase (*Ec*AcpS)^33, 34^ are the most commonly used 4’-PPTases for *apo*→*holo* PTM. *Holo*-ACPs can be further functionalised to their acyl form using a fatty acyl-ACP synthetase (AasS).^35^ AasS activates linear fatty acids as acyl adenylates, which are subsequently loaded onto the ACP via the terminal thiol group of the 4′-PP arm. Amongst this subclass of adenylating enzymes, the *Vibrio harveyi* B392 AasS (*Vh*AasS) is the most extensively characterised and versatile member,^36-39^ and it is a remarkably useful tool for *in vitro* ACP acylation. We reasoned that the successful PTM of our *de novo* sequences would serve as a foundational demonstration of ACP-like behaviour. Thus, we purified and deployed recombinant *Bs*Sfp, *Ec*AcpS and *Vh*AasS in our PTM pathway (Fig. 3A, Fig S4); *Ec*AcpP was used as a positive control throughout (Fig. S11-13).

**Fig. 3.**
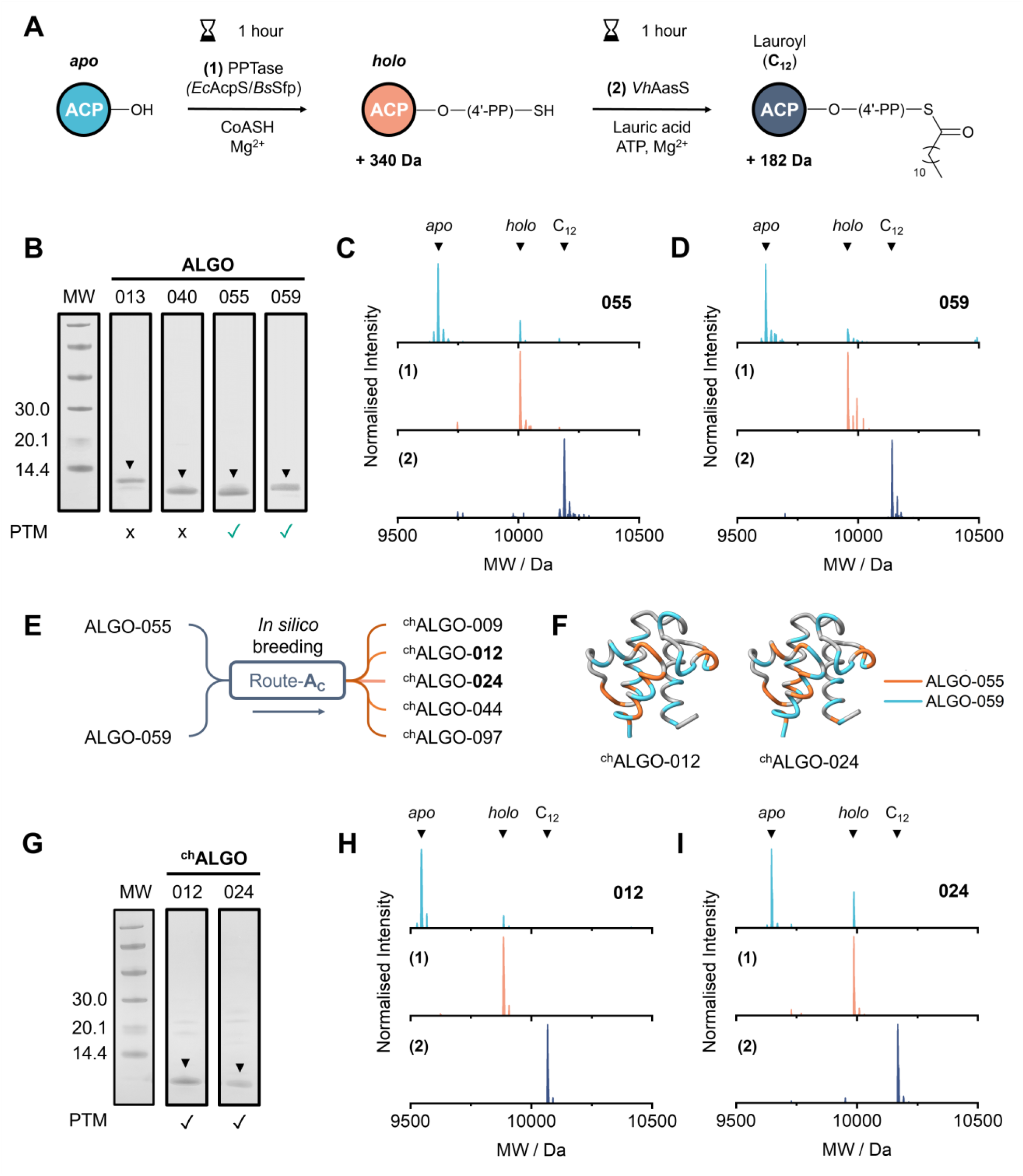
Expression, purification and PTM of ALGO-CP candidate sequences. (A) General scheme for the one-pot, two-step PTM of ACPs *in vitro*, via the sequential 4’-PPTase-catalysed attachment of 4’-PP **(1)** and AasS-catalysed acylation using lauric acid **(2)**. Anticipated MW increases are shown. (B) Annotated SDS-PAGE of purified ALGO-013, 040, 055 and 059. Candidates which undergo PTM are indicated with a green tickmark. An Amersham low molecular weight calibration kit was used as the protein ladder (lane MW). The figure is a composite of two 4-12% Bis-Tris gels that were run, stained and imaged under identical conditions (see also SI) (C) Deconvoluted mass spectra of ALGO-055, tracking PTM from *apo*- (top) to *holo*- (middle) to C_12_- (bottom) as previously outlined. *Ec*AcpS was used as the 4’-PPTase for reaction **(1)**. (D) Deconvoluted mass spectra of ALGO-059, tracking PTM from *apo*- (top) to *holo*- (middle) to C_12_- (bottom) as previously outlined. *Ec*AcpS was used as the 4’-PPTase for reaction **(1)**. (E) Generation of randomized chimeric variants ^ch^ALGO-009, 012, 024, 044 and 097 from parental candidates ALGO-055 and ALGO-059, by repurposing ALGO-CP route-**A**_**C**_. (F) AlphaFold3 models of ^ch^ALGO-012 (pLDDT = 94.48) and ^ch^ALGO-024 (pLDDT = 94.27). Positions/regions inherited from ALGO-055 or ALGO-059 are highlighted in orange or blue, respectively. Identical positions/regions are shown in grey. (G) Annotated SDS-PAGE of purified ^ch^ALGO-012 and ^ch^ALGO-024. Green tickmarks indicate successful PTM. An Amersham low molecular weight calibration kit was used as the protein ladder (lane MW). See also SI. (H) Deconvoluted mass spectra of ^ch^ALGO-012, tracking PTM from *apo*- (top) to *holo*- (middle) to C_12_- (bottom) as previously outlined. *Ec*AcpS was used as the 4’-PPTase for reaction **(1)**. (I) Deconvoluted mass spectra of ^ch^ALGO-024, tracking PTM from *apo*- (top) to *holo*- (middle) to C_12_- (bottom) as previously outlined. *Ec*AcpS was used as the 4’-PPTase for reaction **(1)**.

From the previous subset of *r* = 0.60 sequences sampled for property analysis, seven candidates – ALGO-(013, 023, 040, 044, 055, 057, 059) – were randomly selected for *in vitro* characterisation (Table S13, Fig. S2). The candidate sequences were codon-optimised for *E. coli*, synthesised and cloned into pET28a with a C-terminal His-tag for ease of purification. Following heat-shock transformation and antibiotic selection, overnight protein expression could be induced with 1 mM IPTG at low temperatures (18 °C). Of the seven candidates, ALGO-(023, 044, 057) were expressed as insoluble inclusion bodies, and were not taken further. The remaining candidates could be captured using Ni^2+^-affinity resin and further cleaned using a PES centrifugal filtration unit (Fig. 3B and Fig. S5-8).

Following dialysis, the identity and PTM status of each soluble ALGO candidate was confirmed by LC/ESI-MS (Table S14, Fig S18-20, S22), using purified *E. coli* AcpP as a positive control (Fig. S15-17). Notably, both ALGO-055 and ALGO-059 were purified partly in *holo*-form, as indicated by a molecular weight (MW) increase of 340 Da consistent with the covalent attachment of 4′-PP. Encouragingly, this implied some *in vivo* interaction with the endogenous *Ec*AcpS during protein expression. Both candidates could be readily converted to their respective *holo* forms *in vitro* using purified *Ec*AcpS, in the presence of Mg^2+^, CoASH and dithiothreitol (DTT, see Fig. 3C-D and Fig. S21, S23). Interestingly, both ALGO-055 and ALGO-059 could not be fully converted by *Ec*AcpS when immobilised on Ni^2+^-affinity resin, which is an established method for the reliable and complete PTM of His-tagged ACPs.^40^ This observation suggests that immobilisation could impose a significant conformational constraint on these ALGO sequences, hampering productive engagement with *Ec*AcpS (Fig. S24-25). Furthermore, only partial conversion was achieved using *Bs*Sfp, both *in vitro* (using purified recombinant enzyme) and *in vivo* (using the *sfp* knock-in strain *E. coli* BAP1),^41^ suggesting that these sequences struggle to establish productive PPIs with this 4’-PPTase (Fig. S26-27). Conversely, neither ALGO-013 nor ALGO-040 could undergo *apo*→*holo* PTM at all, and these candidates were not examined further (Fig. S28-29).

Using our standout candidates ALGO-055 and ALGO-059, we next attempted the covalent attachment of a lauroyl (C_12_) acyl chain *in vitro* using *Vh*AasS. The acylation of both *holo*-ALGO-055 and *holo*-ALGO-059 proceeded in the presence of Mg^2+^, ATP, DTT, lauric acid and 1% DMSO. Gratifyingly, we observed >99% acylation of *holo*-ALGO-055 and *holo*-ALGO-059 by LC/ESI-MS, as signified by a MW increase of 182 Da, after only 1 hour of room temperature incubation (Fig. 3C-D and Fig. S30-31). Thus, we identified two unique ACP-like sequences that can not only acquire 4’-PP, but can also fix acyl cargo.

Encouraged by these results, we configured ALGO-CP (via route-**A**_**C**_) to stochastically generate chimeric variants of ALGO-055 and ALGO-059, mimicking the process of molecular breeding to expand sequence diversity within the AcpP design space. A sequence alignment of both ALGO candidates was used as input. From 5000 generated sequences, five (^ch^ALGO-[009, 012, 024, 044, 097]) were sampled from a random subset (n = 100) for experimental testing (Fig. 3E, Table S13). The chimeras ^ch^ALGO-012 and ^ch^ALGO-024 (Fig. 3F) were expressed in soluble form and could be further purified by Ni^2+^-affinity chromatography (Fig. 3G, S9-10). As before, these proteins were also recovered partly in *holo*- form, signalling *in vivo* PTM by endogenous *Ec*AcpS. Like their parental sequences, both candidates could readily undergo complete PTM (*apo*→*holo*→acyl) *in vitro* using *Ec*AcpS and *Vh*AasS (Table S14, Fig. 3H-I, Fig. S32-37).

To the best of our knowledge, both ALGO-055 and ALGO-059, and their chimeric variants ^ch^ALGO-012 and ^ch^ALGO-024, are the first examples of expressible, soluble and modifiable ACPs designed entirely *de novo* by a computer algorithm. More broadly, this result showcases how ALGO-CP can be tuned to probe the sequence design space and to create further diversity from initial successful designs.

### *In silico* biophysical characterization

The canonical *Ec*AcpP is a highly acidic and conformationally dynamic α-helical bundle. This combined negative electrostatic potential and structural plasticity is crucial for substrate shuttling and PPIs. To assess whether our ALGO sequences share these characteristics with natural homologues, we applied a combination of complementary computational analyses.

Secondary structure prediction using PSIPRED^42^ indicates that both ALGO sequences are likely to be predominantly α-helical (Fig. S38-39), with complementary DISOPRED3^43^ analysis predicting minimal intrinsic disorder (Fig. S40-41). We further generated high-confidence structural models of our ALGO variants in their *apo*- form (created using AlphaFold3, pLDDT = 93.74 and 93.52 for ALGO-055 and ALGO-059, respectively, Fig. S42). These models display well-ordered α-helices typical of an ACP fold, with solvent-exposed surfaces that exhibit similar Coulombic electrostatic potential and molecular lipophilicity potential to the 1.1 Å crystal structure of *Ec*AcpP (PDB: 1T8K, Fig. S43-44).^44^

These predicted structural models were subjected to multiple 1 µs molecular dynamics (MD) simulations, with *Ec*AcpP used as the reference system. The structural stability of all proteins was assessed in *apo*-, *holo*- and C_12_-acyl states. In brief, the protein backbone of both *holo*-*Ec*AcpP and *holo*-ALGO-059 appears to be partly destabilised by the presence of the 4’-PP group, as reflected by larger RMSD fluctuations relative to their *apo*-forms. A partial recovery of conformational stability is observed upon acylation, especially ALGO-059, likely due to favourable interactions between the C_12_ alkyl chain and the hydrophobic residues lining the ACP core. In contrast, ALGO-055 equilibrated rapidly and remained comparatively consistent across *apo*-, *holo*- and acyl-forms, indicating intrinsic sequence-structure features that buffer 4’-PP-induced perturbations. To investigate the relationship between protein stability and 4’-PP dynamics, the RMSD of the Ser-4′-PP moiety was calculated relative to a pocket-bound reference conformation (Fig S45A). Indeed, replica 2 of *holo*-*Ec*AcpP (Fig. S45B), and replica 1 of *holo*-ALGO-059 (Fig S45C) display a concomitant increase in the RMSD of the 4’-PP moiety and the backbone RMSD of the protein, associated with the progressive displacement of the prosthetic group from the ACP core (∼230 ns and ∼250 ns for *holo*-*Ec*AcpP and *holo*-ALGO-059, respectively). In contrast, the movement of 4’-PP does not induce backbone destabilisation in *holo*-ALGO-055. This analysis reveals a correlation between the fluctuation of 4’-PP and protein RMSD, although it cannot definitively determine whether 4’-PP displacement precedes or follows protein destabilisation.

Further DSSP analysis revealed that both *holo*- and *acyl*- forms exhibit localised secondary structure rearrangements, primarily transient transitions from α-helical regions into turns and bends (Fig S46). These structural changes are more evident in *Ec*AcpP and ALGO-059, whilst ALGO-055 shows a more conserved secondary-structure pattern, consistent with its enhanced conformational stability. To further dissect helix-specific contributions to ACP stability, we analysed the temporal evolution of α-helical content for the four canonical helices (αI–αIV) across all proteins and states (Fig 5). This per-helix analysis extends the global DSSP profiles by isolating secondary structure dynamics within individual helical segments. ALGO-059 exhibits the most pronounced helix destabilisation, particularly in helices αI and αIII. Helix αII, which harbours the 4’-PP attachment site, shows a marked decrease in α-helical content across all proteins in their *holo*- forms. This is attributable to the dynamic motion of the 4’-PP prosthetic group, which repeatedly enters and exits the hydrophobic binding pocket, as confirmed by principal component analysis/clustering centroid analysis (Fig S47).

In general, whilst ALGO-055/059 may capture key global properties of natural AcpP homologues, our MD simulations point toward distinctive conformational behaviours. Relative to *Ec*AcpP, ALGO-055 demonstrates superior stability across all states, whilst ALGO-059 suffers pronounced plasticity and α-helical loss, occasionally approaching complete unfolding in select helices. These differences appear to be driven by the extent of coupling between 4’-PP dynamics and the protein backbone, with ALGO-059 more perturbed by 4’-PP motions. Collectively, these results suggest that subtle sequence-dependent effects give rise to markedly different dynamic responses.

### CD spectroscopy of ALGO variants

Though robust, it was important to consider that our MD simulations were initialised using pre-folded and energy-minimised models, and therefore may not capture the full range of conformations accessible in solution. Thus, we sought to experimentally validate our biophysical predictions using CD spectroscopy. Based on our *in silico* data, we anticipated subtle differences in helicity in response to different PTM states, using *Ec*AcpP as a comparator.

However, to our surprise, neither *apo*/*holo*-ALGO-055 nor *apo*/*holo*-ALGO-059 demonstrated the diagnostic stalls of an α-helix when studied by CD spectroscopy (Fig 6A-C). In stark contrast to our *in silico* predictions, the CD spectra of *apo/holo*-ALGO-055 and *apo/holo*-ALGO-059 deviate significantly from the characteristic double minima (208 nm and 222 nm), strong maxima (193 nm) and isodichroic point (201 nm) of a well-folded helical protein, as exemplified by *apo*/*holo*-*Ec*AcpP (Fig. 4D). Given that some ACPs, such as *V. harveyi* AcpP, are natively unfolded in the absence of Mg^2+^, we supplemented the buffer with 5 mM MgCl_2_ to evaluate whether the ALGO variants can recover their helicity with the assistance of divalent cations. Remarkably, this supplement only slightly perturbed the CD spectra and did not clearly restore helicity to either ALGO protein (Fig S48), indicating that they may well exist as minimally structured, predominantly non-helical ensembles in these forms.

**Fig. 4.**
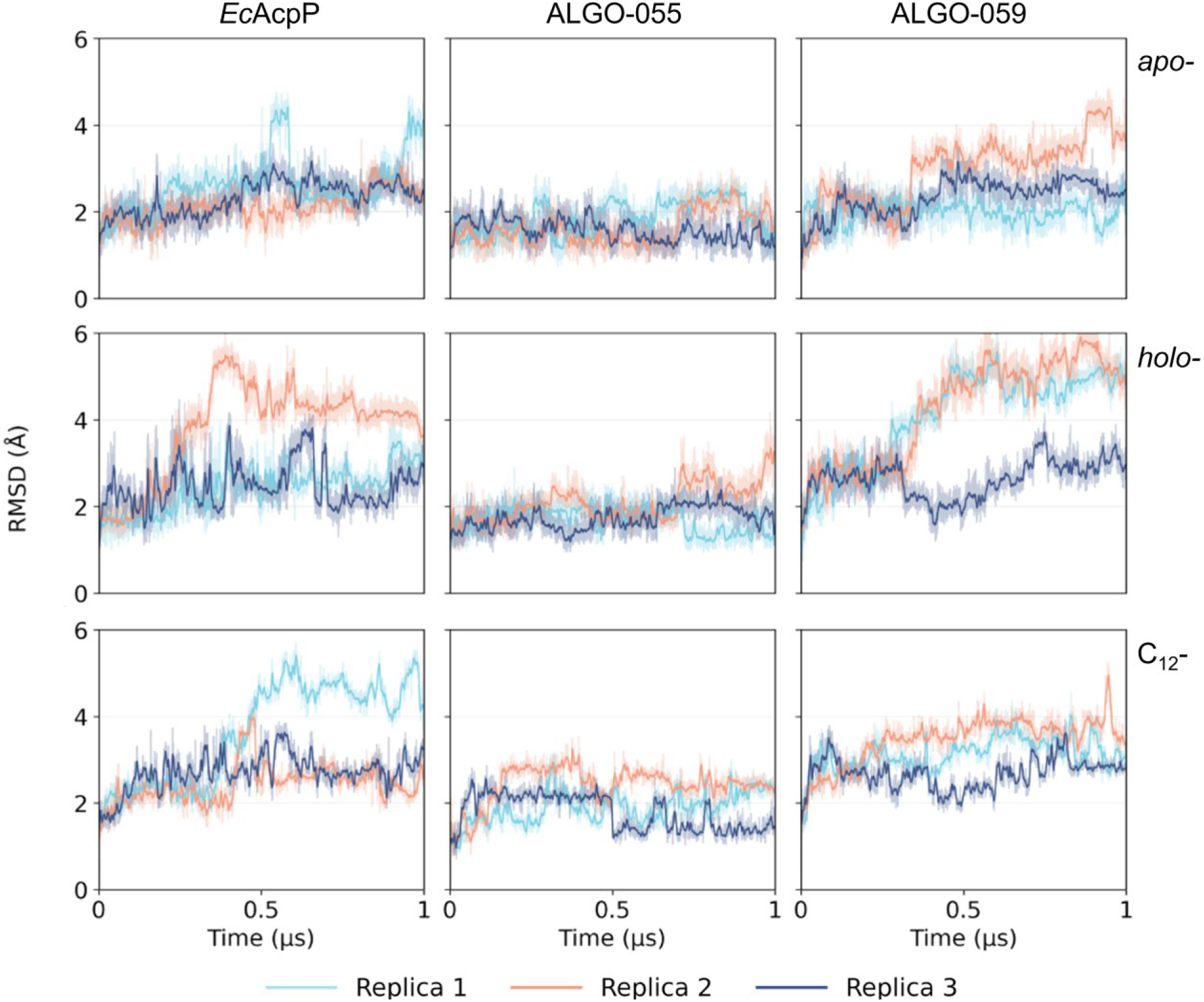
Backbone RMSD profiles of MD trajectories. MD trajectories of *Ec*AcpP, ALGO-055, and ALGO-059 models were analysed in *apo*- (top), *holo*- (middle) and C_12_-acylated (bottom) forms. Each MD simulation was performed in triplicate.

**Fig. 5.**
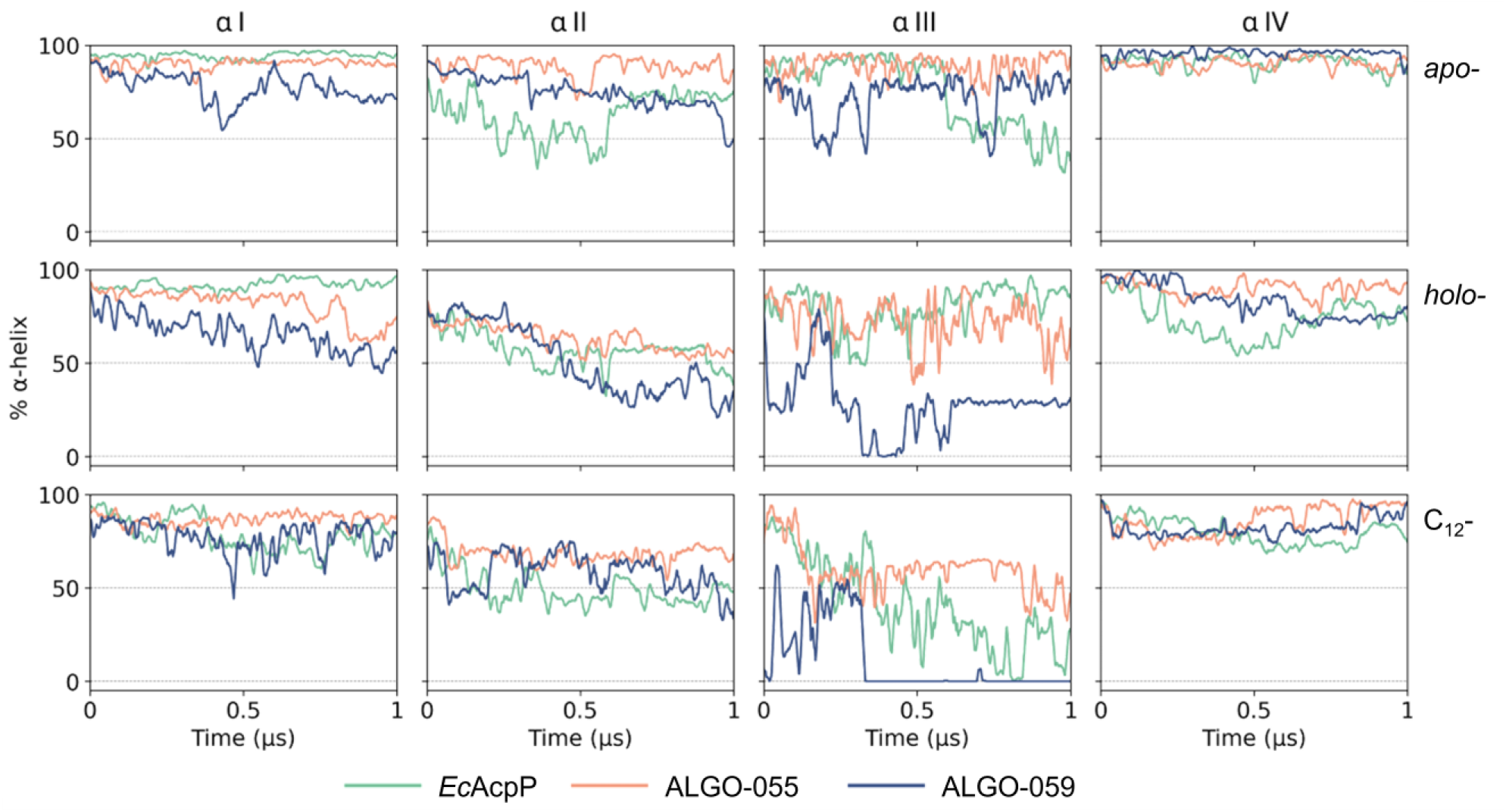
Per-helix α-helical content evolution. MD trajectories of *Ec*AcpP, ALGO-055, and ALGO-059 were examined in their *apo*- (top), *holo*- (middle), and C_12_-acylated (bottom) forms. Helices αI–IV are arranged from left to right.

**Fig. 6.**
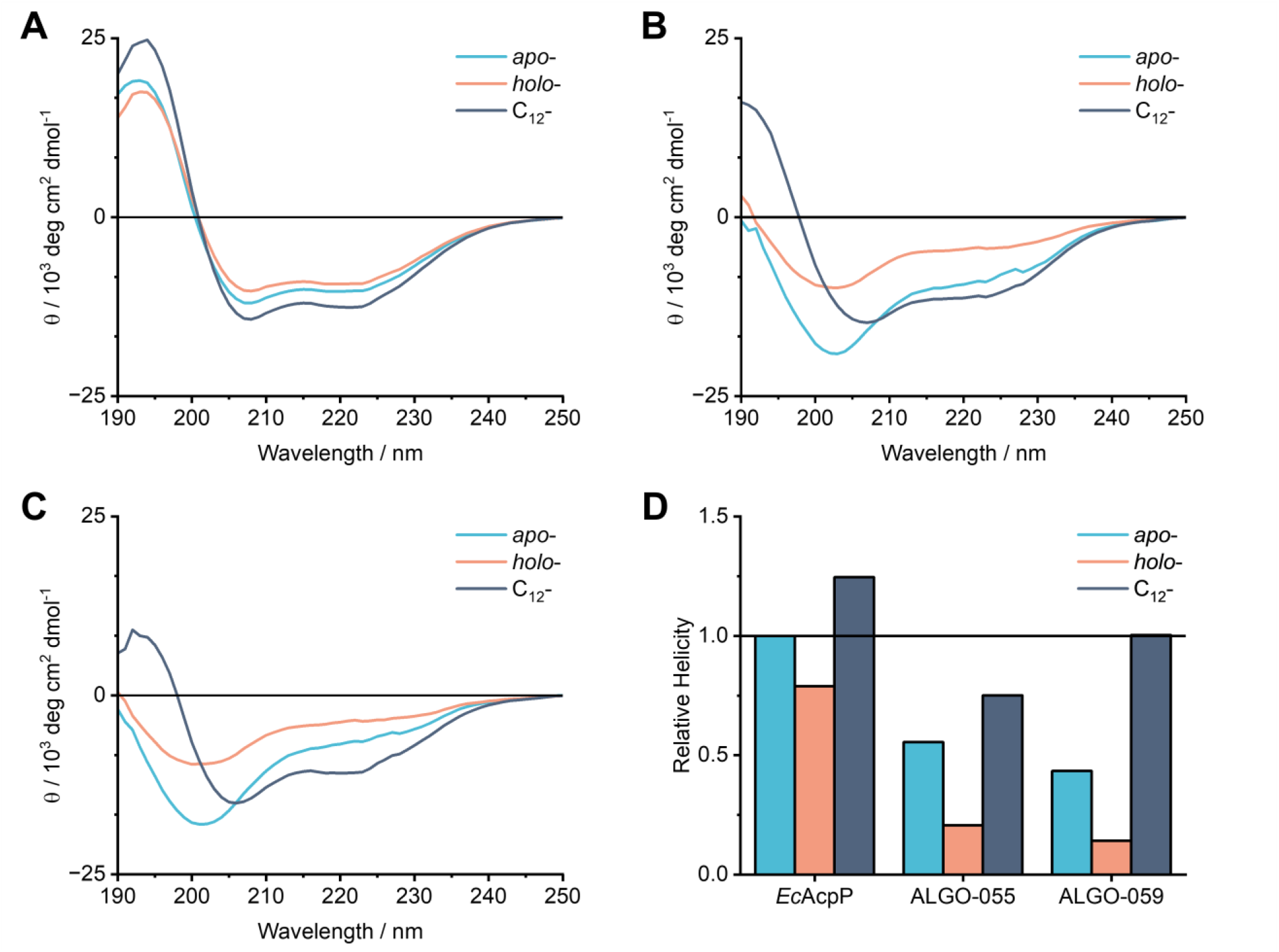
CD spectroscopy and α-helical content across PTM states. (A) CD analysis of *Ec*AcpP. (B) CD analysis of ALGO-055. (C) CD analysis of ALGO-059. (D) Estimated helical content computed from CD spectra, normalised relative to *apo*-*Ec*AcpP.

Prompted by this discrepancy, we then examined whether the acylation of our ALGO proteins could induce structural ordering, either in-full or in-part. Following *holo*→acyl- conversion with *Vh*AasS and C_12_-acid, we observed a marked increase in the helical content of both ALGO variants and *Ec*AcpP. We further purified C_12_-*Ec*AcpP and C_12_-ALGO-055/059 from the acylation mixture and confirmed that these proteins retain their helicity when isolated from other assay components (Fig 6A-C, S11-14), thus showing that this gain-of-structure is intrinsic to the C_12_-acylated state. Binary order-disorder classification,^45^ based on our CD data, predicts that both ALGO variants transition from disordered (*apo*/*holo*) to ordered states upon C_12_-acylation (Table S15).

A comparative analysis of the estimated helical content of *Ec*AcpP, ALGO-055 and ALGO-059 (Fig 6D and Table S16) reveals a consistent drop in helicity upon *apo*→*holo* conversion, in agreement with our MD simulations of pre-folded models. However, we provide further evidence that C_12_-acylation significantly promotes structural organisation across all proteins studied. Whilst this observation aligns with previous literature,^46, 47^ the extent of structural rehabilitation in our ALGO variants (especially ALGO-059) is noteworthy, albeit our analysis does not definitively conclude complete or canonical folding. Nevertheless, our combined CD and MD data supports that: i) acyl cargo bound to the 4’-PP arm can act as a structural chaperone, most likely by organising and burying hydrophobic residues within the protein core, and ii) 4’-PP modification enhances the structural plasticity of natural ACPs by partially destabilising α-helices, which may help to promote promiscuous PPIs.

The discrepancy between our *apo*/*holo* CD spectroscopy results and established findings on *V. harveyi* AcpP prompted a closer inspection of these ALGO sequences. It has been reported that a single A_76_H mutation in *V. harveyi* AcpP restores its helicity via π-stacking interactions with Y_72_, even in the absence of divalent cations.^48^ Whilst ALGO-055 and ALGO-059 retain the equivalent Y_72_ residue, ALGO-055 harbours the “destabilising” A_76_ variant, whereas ALGO-059 carries the “stabilising” H_76_. Taken together with our Mg^2+^ supplementation experiment, this suggests that ALGO-055 and ALGO-059 may be too divergent from natural AcpP homologues for this stabilisation mechanism to be effective. Alongside our PTM results, this points toward an underlying, hitherto undescribed sequence-structure-function relationship. Accordingly, we performed a deeper post-hoc analysis of our ALGO sequences to explore this relationship further.

### Primary sequence analysis

To assess the sequence divergence of our ALGO candidates, we quantified the rarity of all amino acid variations relative to our collection of 2167 AcpP homologues. We then evaluated their evolutionary likelihood and physicochemical dissimilarity relative to *Ec*AcpP using BLOSUM62^49^ and Grantham distance^50^ matrices, respectively.

Our analysis of ALGO-055 and ALGO-059 revealed that both proteins harbour at least 16 amino acid variations that are represented by <5% of the AcpP homologues used in the input MSA. Of these variations, ten (ALGO-055) and seven (ALGO-059) are present in <1% (Table S17-18, Fig. 7A). These variations occur across the sequence length of the proteins, with the regions typically associated with helix αII and αIII being the best-preserved by ALGO-CP. By BLASTp, the closest homologues of ALGO-055 and ALGO-059 belonged to *Providencia heimbachae* (69.23% identity) and *Lacimocrobium alkaliphilum* (65.38% identity), respectively. When compared to *Ec*AcpP, both proteins contain ≥30 amino acid variations, approximately half of which are non-conservative by BLOSUM62 analysis with median Grantham distance scores of 89 (ALGO-055) and 78 (ALGO-059) (Table S17-18, Fig. 7B-C). Remarkably, ALGO-055 and ALGO-059 share only 51.28% sequence identity with each other, with 17 non-conservative variations scoring a median Grantham distance of 91 (Table S19).

**Fig. 7.**
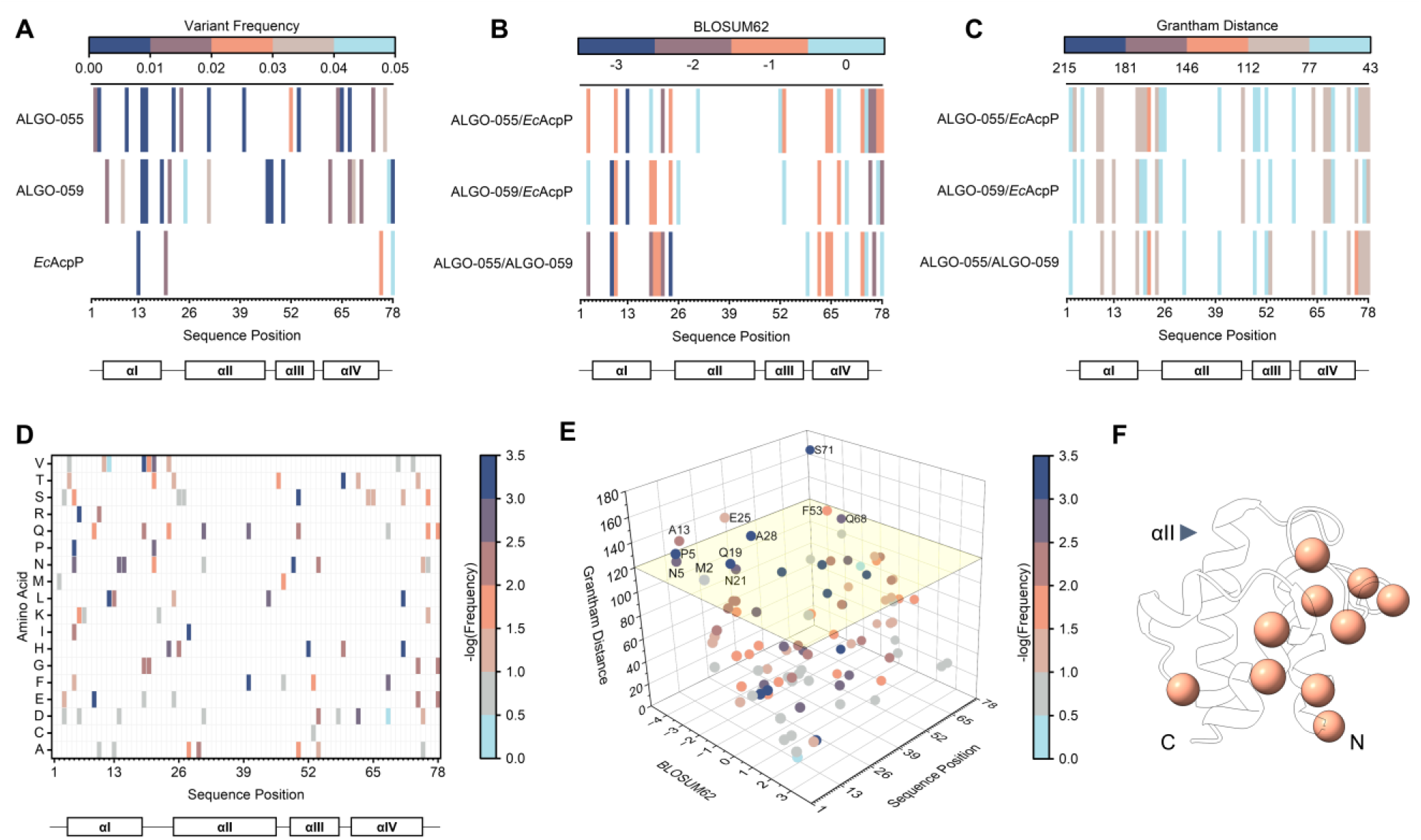
(A) Distribution of rare amino acid variations (≤5% occurrence) in ALGO-055 and ALGO-059, relative to the input multiple sequence alignment (MSA) of AcpP homologues (n = 2167) used for ALGO-CP. *Ec*AcpP was also studied for comparison. (B) BLOSUM62 analysis of non-conservative amino acid variations identified by sequence alignment. BLOSUM62 scores >0 are not coloured. (B) Grantham distance analysis of physicochemically-divergent amino acid variations observed by sequence alignment. Grantham distances <43 are not coloured. (D) Frequency and distribution of amino acid variations found exclusively in unsuccessful ALGO sequences. Neither tryptophan nor tyrosine were identified in this analysis and are therefore not featured in the heatmap. (E) Grantham distance and BLOSUM62 scores of amino acid variations found exclusively in unsuccessful ALGO sequences, determined by sequence alignment with *Ec*AcpP (see also Table S18). Datapoints are colour coded by -log_10_(frequency). Non-conservative variants with Grantham distances ≥120 (above yellow plane) are labelled. (F) Positions of non-conservative, physiochemically-divergent variations mapped onto *Ec*AcpP (PDB: 1T8K). The position of helix αII is indicated. N- and C- termini are labelled.

Overall, the Grantham distance scores of many of these variations fall within the lower 50^th^ percentile of all possible amino acid pairings (i.e., <96), suggesting moderate, rather than extreme, physicochemical divergence. Nevertheless, it remains possible that the accumulation of these rare and non-conservative amino acid variations contributes to the observed destabilisation of the canonical α-helical structure, though the precise variants/mechanisms responsible for this effect are difficult to predict. We note, however, that ALGO-055 and ALGO-059 preserve several acidic “hotspots”, clustered mostly downstream from the invariant serine, that are known to interface with the highly electropositive docking surfaces of *Ec*AcpS and *Vh*AasS (Fig. S49).^39, 51^ This electrostatic complementarity may still provide adequate priming/orientation for productive PPIs, even in the absence of an ordered structure.

Similarly, it is difficult to ascertain deleterious amino acid variations (or combinations of mutations) from such a limited set of *de novo* sequences. However, a meta-analysis of our initial ALGO candidates identified 95 total variations that are exclusively found in unsuccessful ALGO sequences (i.e., sequences that failed to express in soluble form or undergo PTM) versus *Ec*AcpP, ALGO-055 and ALGO-059 (Table S20, Fig. 7D). Of these, 69 are featured in <10% of all 2167 AcpP homologues studied, including 38 that are found in <1%. In total, we identified 11 non-conservative amino acid variations with Grantham distances >120 when compared to *Ec*AcpP; many of these are clustered around αI and αIV, with none occurring along the crucial recognition helix αII (Table S20, Fig. 1A-C, 7E-F). We propose that sequences enriched with these rare and/or unorthodox amino acid variants may be predisposed to poor solubility or functionality, which may explain why such variations were selected against during AcpP evolution.

Both ALGO-055 and ALGO-059 occupy a peculiar region of an otherwise fragile sequence design space, in which some biochemical function (specifically, essential PTM) is retained in the absence of a hallmark α-helical structure. In 2005, Christopher Walsh and team used phage display to identify a 11-mer PCP fragment (ybbR-13) that undergoes *apo*→*holo* PTM via interaction with *Bs*Sfp.^52^ Notably, even this highly truncated peptide adopts a clear α-helical conformation in solution, further highlighting the idiosyncrasy of these ALGO designs. The behaviour of these proteins may reflect a disordered “precursor-like” state to the canonical ACP fold, one in which an amorphous configuration of key contact residues is still sufficient for PTM. Indeed, it is widely appreciated that intrinsically disordered proteins can adopt three-dimensional folds via proximity to/engagement with a protein partner, and it is this structural malleability which enables promiscuous interactions.^53^ As biosynthetic assemblies and physiological requirements grew more complex, it is possible that the emergence of a defined α-helical fold may have conferred advantages in partner selectivity and/or kinetic efficiency through more nuanced PPIs, a hypothesis which could be explored via further phylogenetic/co-evolutionary analyses. Based on our current data, we postulate that ACP sequences with greater helical propensity – with or without the assistance of chemical chaperones - were gradually selected to meet the demands of biosynthetic precision and function, rather than out of necessity for PTM alone. Our findings also provide further evidence that the canonical ACP fold may have partially emerged as a consequence of cargo engagement, which in turn would facilitate substrate sequestration and protection. Assuming that the canonical helical structure of AcpP is essential for cell fitness, we propose that ALGO-055 and ALGO-059 could be evolved, under *in vivo* selective pressure, to determine whether and how this fold can be re-acquired to support complex physiological functions. To this end, the *E. coli* Δ*acpP* knockout strain CY1877 would make an excellent background for a live/death complementation screen.^22^

## Conclusion

Despite their small size, ACPs are deceptively complex and nuanced proteins. Their sheer versatility enables certain subclasses (chiefly AcpP) to operate at the heart of both cell metabolism, division and physiology. In this study, we developed a tuneable sequence-generating algorithm, ALGO-CP, to create novel AcpP-like sequences that both skirt and surpass the natural design constraints of this ancient protein family. Even from a limited sample of *de novo* sequences, we uncovered surprising insights that both facilitate and test our understanding of ACP engineering and evolution. We further observe the tendency for state-of-the-art structural prediction tools to over-predict α-helicity in our *de novo* sequences, perhaps reflecting training biases that make this approach less effective for our specific application. Furthermore, some discrepancies between our MD predictions and CD data may arise from the fact that simulations were initiated from pre-folded models, thereby biasing the system toward helical conformations. Consequently, MD simulations in this context probe the stability of an imposed fold rather than its spontaneous formation, potentially overestimating the intrinsic folding propensity of these *de novo* sequences. Nevertheless, for the first time, we have set the groundwork for computer-led ACP design supported by a simple, yet effective, screen for essential ACP-like behaviour. We propose that ALGO-CP could be adapted to study different ACP subclasses across different kingdoms of life, including those involved in fungal polyketide biosynthesis^54^ or mitochondrial function.^55, 56^ With further experimental feedback, we envision that ALGO-CP, especially when coupled with machine learning and high-throughput screening, could become a powerful tool for pioneering entirely new ACP lineages, with potential applications in metabolic engineering and beyond.

Future work could investigate further structural and biophysical details using ^1^H-^15^N HSQC correlation spectra and thermal denaturation experiments, as well as kinetic differences in PTM between our ALGO variants and natural ACPs. Substrate sequestration behaviour of our acyl-ALGO designs could also be investigated further using vibrational spectroscopy.^9, 57^ Finally, the systematic sampling of sequence designs across different *r* weighting coefficients may provide additional insights into sequence-structure-function relationships.

## Experimental

All materials, expression plasmids, media, reagent stock solutions and LC/ESI-MS methods are detailed in the SI.

### Retrieval of AcpP homologues

Sequence homologues were retrieved from UniRef90 via ConSurf.^58^ In brief, 17 AcpP homologues (table S3) were used as queries to retrieve the top 150 closest sequence homologues each (95-50% sequence identity). These homologous sequences were compiled into a single FASTA file, which was filtered for duplicate/incomplete entries and aligned via MAFFT v7^59^ using default parameters.

### Amino acid and sequence properties

For ALGO-CP, the requisite physicochemical data for each proteinogenic amino acid were retrieved from AAindex.^60^ Sequence GRAVY and MW were computed using Multiple Protein Profiler 1.00.^59^ Sequence pI was computed using Isoelectric Point Calculator 2.0.^61^

### Predictive structural modelling and molecular dynamics simulation

All structural predictions of were performed using AlphaFold3^31^ (via AlphaFold server). General static model viewing and analysis was performed using ChimeraX v1.8.^62^

GROMACS 2023^63^ package was used to run MD simulations of *apo*-, *holo*- and acylated systems. AMBER99sb was used as force field and TIP3P as water model. The solvated system was neutralized with counterions and a 0.1 M NaCl concentration was added. Energy minimization was performed using a maximum of 50,000 steps of steepest descent with grid neighbour searching, no constraints, and no thermostating. Long-range electrostatic interactions were treated using the PME method, while short-range Lennard-Jones and van der Waals interactions were calculated using a 12 Å distance cutoff. The system was equilibrated in four steps. Three consecutive NVT simulations were used to gradually heat the system from 0 K to 300 K in increments of 100 K, each lasting 0.1 ns, followed by a 1 ns NPT equilibration at 300 K. Harmonic positional restraints were applied to the protein and eventual prosthetic group’s heavy atoms throughout the equilibrations, using a force constant of 1000 kJ mol^−1^ nm^−2^. Velocities were propagated between equilibration steps. Exclusively for the *holo*-simulations of the ACP enzymes ALGO-059 and ALGO-055, three NPT equilibration phases were performed, during which harmonic positional restraints on the protein backbone had to be gradually reduced from 1000 to 100 kJ mol^−1^ nm^−2^.

Production runs were carried out in the NPT ensemble at 300 K without any restraints, using a 2 fs integration step. Each ACP was simulated in three independent replicas (1 μs). The temperature was set at 300 K using a velocity-rescale thermostat.^64^ Long-range electrostatics were treated with PME, bonds to hydrogens were constrained with LINCS,^65^ and van der Waals interactions used a Verlet cutoff with force-switching. Trajectories and energies were saved every 10 ps, and PBC were applied in all directions.

The covalently attached prosthetic group (4′-phosphopantetheine) and its acylated derivative (lauroyl-/C_12_) were parametrised prior to MD simulations. Their structures were extracted from the PDB entries, respectively 5H9H and 2FAE PDB IDs. Missing atoms were added, partial atomic charges were calculated using the DFT method, and bonded and non-bonded parameters were assigned using the GAFF2 force field.^66^ ACPYPE^67^ was employed to generate GROMACS-compatible topologies for the ligand. The ligands were incorporated into the target ACPs by extracting the 4′-phosphopantetheine and lauroyl groups from the donor X-ray structures using PyMOL, manually positioning them into the binding pockets, and building the covalent linkage to the invariant serine residue to model *holo*- and acyl states; positional restraints were applied to ensure stability during equilibration.

Trajectories were centred and fitted to the protein backbone prior to analysis. Secondary structure was analysed using the DSSP dictionary, integrated in GROMACS. Structural stability was evaluated through backbone RMSD. Trajectories were inspected using VMD.^68^

To further characterise the conformational space of the *acyl*- and *holo*-ACP systems, principal component analysis (PCA) and clustering were performed on all three independent replicas for each system using MDTraj^69^. Trajectories were merged and non-hydrogen atoms of the covalently attached prosthetic group or acyl chain were analysed. Cartesian coordinates were projected onto the first two principal components and clustered using a quality-threshold-based approach with a 4.5 Å cutoff for acyl- and 5.5 Å for *holo*-. Representative frames were selected as those closest to each cluster centroid, and cluster populations were computed as fractions of frames.

### ALGO-CP overview

ALGO-CP begins by extracting position-specific data from alignment blocks, including residue frequency and the average H_KD_, pI, and vdW. Heavily-gapped positions can be excluded from analysis; in this study, only alignment blocks containing ≥50 residues were considered. For any given aligned position *i*, route-**A**_**C**_ generates a probability distribution 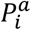 based solely on the frequency of each observed amino acid in the alignment block:

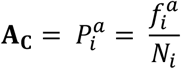

Where *f* is the frequency of a given amino acid, and *N* is the total number of aligned residues in the alignment block. For alignment blocks which exhibit ≥90% amino acid identity, the modal amino acid is selected automatically, thus ensuring the preservation of critical functional residues such as the catalytic serine. In contrast, route-**A**_**p**_ interprets physicochemical descriptions of alignment blocks, rather than explicit conservation information. For any given aligned position *i*, the mean values of the three physicochemical properties - H_KD_, pI and vdW - are computed across the alignment block:

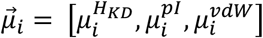

Each proteinogenic amino acid *a* is then scored based on the similarity of its physicochemical profile 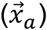 to 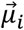, where:

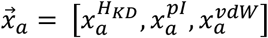

The similarity score 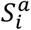 is calculated using the weighted reciprocal difference for each physicochemical property:

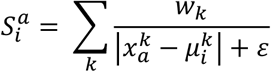

Where *k* is the physicochemical property, *w*_*k*_ is a user-defined weight assigned to the physicochemical property, and ε is a miniscule constant (10^−6^) to prevent division by zero. In this study, the weights for H^KD^, pI and vdW were arbitrarily set to 0.22, 0.44 and 0.33, respectively; we encourage users to experiment with these weights. These scores are subsequently normalised to give a distinct probability distribution based on this physicochemical data:

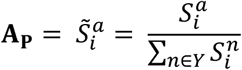

Where 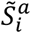 is the normalised similarity score for a given amino acid, and *Y* denotes the set of all 20 canonical (proteinogenic) amino acids. This scoring function favours amino acids whose physicochemical properties closely match the local average at each position, enabling route-**A**_**p**_ to explore substitutions not necessarily observed in natural sequences, but are compatible with the general physicochemical properties of the sequence position.

The distributions generated by route-**A**_**C**_ and route-**A**_**P**_ can be weighted and combined into a unified probability profile **A**_**U**_, allowing the user to bias amino acid selection towards conservation or physicochemical-guided design:

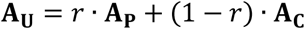

Where *r* is a user-defined weight ranging from 0-1. ALGO-CP then generates new sequences end-to-end by stochastically sampling amino acids according to the positional A_U_.

### General procedure for recombinant expression

For each expression construct used in this work, an exhaustive list of antibiotics, growth media, expression conditions can be found in table S2. The desired expression construct was used to transform chemically-competent *E. coli* BL21 (DE3) cells via the heat-shock method. Colonies were developed overnight on antibiotic VLB-agar plates. A single colony was propagated overnight in antibiotic VLB media (50 mL) by shaking incubation (37 °C). The cells were subcultured (OD_600_ = 0.1, 37 °C) in fresh antibiotic VLB or Autoinduction Media (500 mL) as required, until mid-log phase. Cultures requiring manual induction were cooled to room temperature, and protein expression was induced by the addition of IPTG. All protein expression proceeded with rigorous agitation at the requisite temperature and time. The biomass was harvested by centrifugation using a Fiberlite F14-6 × 250y fixed-angle rotor (7000 rpm, 5 minutes), consolidated into 2-5 g pellets using a Fiberlite F15-8 × 50cy fixed angle rotor (7000 rpm, 10 minutes) and stored at -20 °C.

### Purification of ALGO-(013, 040, 055, 059) and *Ec*AcpP

Potassium phosphate (100 mM, pH 7.6) with glycerol (10% v/v) was used as the buffer base throughout, with varying imidazole strengths as described. Cell pellets were defrosted and resuspended (10% w/v) in ice-cold resuspension buffer (10 mM imidazole). Benzamidine hydrochloride (1 mM) was added to the resuspension and the cells were lysed by sonication (10 second pulse, 10 second cooldown, 15 cycles) on ice. Cell debris was pelleted by high-speed centrifugation using a Fiberlite F15-8 × 50cy fixed angle rotor (13000 rpm, 45 minutes, 4 °C). The cell-free extract was clarified by filtration (Millex-HP 0.45 µm polyethersulfone, Merck). The recombinant protein was captured from cell-free extract using a Histrap HP (1 mL) column and washed (35 mM imidazole, 15-20 mL). Protein was eluted in fractions (300 mM imidazole, 1 mL) and pooled. ALGO proteins were passed through a centrifugal filtration unit (30-100 kDa MWCO) to remove protein contaminants, and the flowthrough was dialysed in imidazole-free buffer. *Ec*AcpP was polished by size exclusion chromatography using HiLoad 16/600 Superdex 75 pg (120 mL) column. For long-term storage, protein aliquots were flash frozen in liquid nitrogen and stored at -80 °C.

### Purification of *Bs*Sfp

HEPES (200 mM, pH 7.5) with glycerol (10% v/v) was used as the buffer base throughout, with varying imidazole strengths as described. Cell pellets were defrosted and resuspended (10% w/v) in ice-cold imidazole-free resuspension buffer. Benzamidine hydrochloride (1 mM) was added to the resuspension and the cells were lysed by sonication (10 second pulse, 10 second cooldown, 15 cycles) on ice. Cell debris was pelleted by high-speed centrifugation using a Fiberlite F15-8 × 50cy fixed angle rotor (13000 rpm, 45 minutes, 4 °C). The cell-free extract was clarified by filtration (Millex-HP 0.45 µm polyethersulfone, Merck). The recombinant protein was captured from cell-free extract using a HiTrap TALON Crude (1 mL) column and washed (10 mM imidazole, 15-20 mL). *Bs*Sfp was eluted in fractions (150 mM imidazole, 1 mL), pooled and dialysed in imidazole-free buffer. For long-term storage, protein aliquots were flash frozen in liquid nitrogen and stored at -80 °C.

#### *In vitro apo*→*holo*→acyl PTM of ALGO candidates and *Ec*AcpP

An *apo*→*holo* conversion mixture (100 µL) containing purified ACP (20 µM), *Bs*Sfp/*Ec*AcpS (2 µM), MgCl_2_ (5 mM), DTT (1 mM) and CoASH (100 µM) was prepared in potassium phosphate buffer (100 mM, pH 7.6) with glycerol (10% v/v). The conversion proceeded at room temperature for 1 hour, after which *Vh*AasS (3 µM), ATP (100 µM) and lauric acid (100 µM, from a 1000X stock prepared in DMSO, see SI Materials and Methods) were added directly to initiate *holo*→*acyl* conversion (104 µL total volume). *Holo*→*acyl* conversions proceeded at room temperature for an additional hour prior to LC/ESI-MS analysis.

#### Circular Dichroism Spectroscopy

CD spectra were collected using an Aviv Model 410A circular dichroism spectropolarimeter. Protein samples (10 µM) were prepared in sodium phosphate buffer (50 mM, pH 7.6, 300 µL). Samples were dispensed into a High Precision Quartz SUPRSIL cuvette with 0.1 cm pathlength (Hellma Analytics). The spectropolarimeter was purged with nitrogen for two hours. The UV lamp was warmed for 30 minutes, after which CD spectra were collected at 25 °C with a range of 190–250 nm using bandwidth of 1 nm, a 0.5 nm step size, averaging time of 3 seconds, and 5 scans. The resulting spectrum was converted to units of mean residue ellipticity (MRE) using the protein amino acid sequence and the sample concentration. Analysis of protein secondary structure characteristics was conducted by uploading normalized data in units of MRE to the web server BeStSel.^70, 71^

## Abbreviations

ATP: adenosine triphosphate
DSSP: dictionary of secondary structure of proteins
IPTG: isopropyl β-D-thiogalactopyranoside
RMSD: root mean square deviation

## Author contributions

M.A.H conceived of and lead the study with conceptual input from D.J.C and L.K.C. M.A.H designed the algorithm, performed wet and dry lab experiments and analysed the data. G.K and Z.O performed wet lab experiments and data analysis under supervision of L.K.C and M.A.H. G.A.T and N.S performed MD simulations and data analysis under supervision of F.S with conceptual guidance from M.A.H. M.AH, G.A.T, N.S., F.S, L.K.C and D.J.C wrote the manuscript. All authors have given approval to the final version of the manuscript.

## Data Availability

The data supporting this article have been included as part of the Supplementary Information. The code for ALGO-CP and related data analysis can be found at https://github.com/MAHerrera-94/ALGO_CP. The version of the code employed for this study is version 1.0.

## Acknowledgements

We thank the Biotechnology and Biological Sciences Research Council (BBSRC, grant number: BB/Y002210/1) and the National Institutes of Health (NIH, grant number: 2R15GM12704-03) for funding. MS data were acquired on instruments funded by the Engineering and Physical Sciences Research Council (EPSRC, EP/K039717/1). We would like to thank Dr. Christopher Wells-Wood and Dr. Antonia Mey for their invaluable advice and enthusiastic support of this work. We thank Dr. Gustavo Perez-Ortiz (University of Edinburgh) for the expression and purification of *Vh*AasS and *Ec*AcpS. We also thank Prof. Robert Fairman (Haverford College) for his assistance with CD spectroscopy.

## Footnotes

M.A.H dedicates this manuscript to his close friends and family.

