## Supplemental Information for "*De novo* acyl carrier proteins display structure-independent modification and sequence novelty"

#### Supplementary Information


| Heading | Contents | Page |
|---|---|---|
| Additional Materials and Methods | Buffers and reagents<br>Plasmids, strains and proteins<br>LC/ESI-MS instrumentation, hardware and parameters.<br>Size-exclusion chromatography following *holo*→C12-acyl conversion. | 2-3 |
| Supplementary Tables | S1-S20 | 4-20 |
| Supplementary Figures | Sequence analysis (Fig. S1-3) | 21-22 |
| | Purification data (Fig. S4-14) | 23-29 |
| | LC/ESI-MS (Fig. S15-37) | 30-41 |
| | Structural modelling and MDS (Fig. S38-S47) | 32-48 |
| | Additional CD data (Fig. S48) | 49 |
| | Additional sequence data (Fig. S49) | 50 |
| | Raw SDS-PAGE images | 51 |
| Plasmid Maps | pALGO and pCHALGO maps | 53-64 |

#### Additional Materials and Methods

**Buffers and reagents.** A summary of all reagent stocks and rich media can be found in Table S1. CoASH, lauric acid, chloramphenicol and kanamycin sulfate were purchased from Sigma-Aldrich. Imidazole, DTT and IPTG were purchased from Fluorochem. Yeast extract and tryptone were purchased from Merck Life Sciences, and SOC recovery medium (for transformations) was purchased from New England Biolabs. VLB-Millar broth, VLB-agar and Autoinduction media was purchased from Formedium. All other chemicals and solvents were purchased from Fisher Scientific. Deionised water ($dH_2O$) for buffers, media and reagent stocks was obtained using a Sartorius purifier. All media was sterilised by autoclave prior to use. All buffers (see table S1) for protein purification were filtered (0.22 µm) and de-gassed under vacuum. For antibiotic selection, the working concentrations of carbenicillin, kanamycin and chloramphenicol were 100, 50 and 30 µg $mL^{-1}$, respectively.

**Plasmids, strains and proteins.** A summary of all expression plasmids used in this study can be found in Table S2. Plasmid pACYC-*sfp* was kindly provided by Ingenza Ltd. Plasmid pSGL-006 was kindly provided by the Prof. Michael D. Burkart research group. Plasmids pALGO-(013, 027, 040, 044, 055, 057, 059) and pCHALGO-(009, 012, 024, 044, 097) were synthesised and cloned by Genscript. Plasmids were propagated using NEB 5-alpha (New England Biolabs) cells and purified using Promega Wizard Plus SV miniprep kit. DNA sequencing was performed by GeneWiz. All purification hardware (including ÄKTA Start and ÄKTA Go FPLC purifiers) were purchased from Cytiva. SDS-PAGE was performed using pre-cast NuPAGE 4-12% Bis-Tris gels and NuPAGE MES running buffer purchased from Invitrogen and used according to manufacturer protocol. Protein gels were stained using InstantBlue Coomassie stain (Abcam). *Vh*AasS and *Ec*AcpS were provided as pre-purified aliquots courtesy of Dr. Gustavo Perez-Ortiz. Protein concentrations were determined using the BCA assay.

**LC/ESI-MS instrumentation, hardware and parameters.** This study was a joint expedition between labs in the University of Edinburgh (UK) and Haverford College (USA). Each lab reproduced the expression, purification and PTM of *Ec*AcpP and ALGO candidates ALGO-055 and ALGO-059, albeit mass analysis was performed using distinctive LC/ESI-MS setups as described below.

*University of Edinburgh*: Proteins were diluted to ~2-5 µM using LC-MS grade water and analysed using a Waters Synapt G2-Si Quadrupole/Time of Flight (TOF) HDMS coupled to a Waters Acquity Class I Plus UPLC, equipped with a Phenomenex Aeris C4 column (200 Å, 3.6 µm, 2.1 mm x 50 mm). Samples were analysed in positive ion mode. LC-MS grade water + 0.1% formic acid was used as Solvent A. ACN + 0.1% formic acid was used as Solvent B. Analytes were resolved using a fixed flow rate (0.2 ml min-1) using the following method parameters: 95% A / 5% B (initial ratio, 1.5 min hold); 5% A / 95% B (3.5 min gradient, 6 min hold), 95% A / 5% B (0.1 min gradient, 0.9 min hold). Mass spectra were analysed and deconvoluted using MassLynx.

*Haverford College*: Proteins were diluted to ~10 μM using LCMS grade water and analysed using an Agilent Technologies InfinityLab G6125B LC/MS coupled with an Agilent 1260 Infinity II LC system, equipped with a Waters XBridge Protein BEH C4 reverse phase column (300 Å, 3.55 μm, 2.1 mm x 50 mm) heated to 45 °C. Samples were analysed via electrospray ionization mass spectrometry (ESI-MS) in positive ion mode. LC-MS grade water + 0.1% formic acid was used as Solvent A. ACN + 0.1% formic acid was used as Solvent B. Analytes were resolved using a fixed flow rate (0.2 ml min-1) using the following method parameters: 95% A / 5% B (initial ratio, 1 min hold); 5% A / 95% B (3.1 min gradient, 1.42 min hold), 95% A / 5% B (0.4 min gradient, 4.08 min hold). Mass spectra were deconvoluted using ESIprot online.

**Size-exclusion chromatography following *holo*→$C_{12}$-acyl conversion.** All buffers used in FPLC were filter sterilized (0.2 μm Nylon membrane filter) and degassed by vacuum, except for small buffer volumes used to wash syringe, needle, and injection port which were not degassed. FPLC was used to separate ACPs from other enzymes after *in vitro* reactions, in preparation for CD spectroscopy. FPLC was performed on an Akta Pure chromatography system (GE Healthcare) equipped with a Superdex 75 Increase 10/300 GL column (GE Healthcare). The column was equilibrated with 50 mM sodium phosphate buffer (pH 7.6) for ~ 90 minutes. A 500 μL sample loop was primed with 1 mL of 50 mM sodium phosphate buffer (pH 7.6), before ~500 μL of sample was injected into the loop. Buffer was run at a flow rate of 0.8 mL/min, with pressure maintained at 0.3-0.4 MPa, and UV absorbance monitored at 280 nm. Fractions were collected in 0.5 mL increments. Fractions containing significant UV absorbance (280 nm) were analyzed by LCMS, and pooled and concentrated by centrifugal filtration for further use. Samples were also confirmed by SDS-PAGE

#### Supplementary Tables (S1-20)

**Table S1** Rich media and reagent stock solutions

| Item | Contents |
|---|---|
| **VLB media** | Yeast extract (5 g $L^{-1}$), tryptone (10 g $L^{-1}$), NaCl (10 g $L^{-1}$) |
| **VLB-Agar** | Yeast extract (5 g $L^{-1}$), tryptone (10 g $L^{-1}$), NaCl (10 g $L^{-1}$), Agar (15 g $L^{-1}$) |
| **Autoinduction Media** | Yeast extract (5 g $L^{-1}$), tryptone (10 g $L^{-1}$), D-glucose (0.5 g $L^{-1}$), α-lactose (2 g $L^{-1}$), $(NH_4)_2SO_4$ (3.3 g $L^{-1}$), $KH_2PO_4$ (6.8 g $L^{-1}$), $Na_2HPO_4$ (7.1 g $L^{-1}$), $MgSO_4$ (0.15 g $L^{-1}$), trace elements (0.03 g $L^{-1}$) |
| **Kanamycin (1000X)** | Kanamycin sulfate (50 mg $mL^{-1}$) in $dH_2O$ |
| **Carbenicillin (1000X)** | Carbenicillin (100 mg $mL^{-1}$) in $dH_2O$ |
| **Chloramphenicol (1000X)** | Chloramphenicol (30 mg $mL^{-1}$) in EtOH |
| **1 M IPTG (1000X)** | IPTG (238 mg $mL^{-1}$) in $dH_2O$ |
| **DTT (100X)** | DTT (15.4 mg $mL^{-1}$) in $dH_2O$ |
| **Lauric acid (1000X)** | Lauric acid (20.0 mg $mL^{-1}$) in DMSO |
| **CoASH (1000X)** | Coenzyme A sodium salt (76.6 mg $mL^{-1}$) in $dH_2O$ |
| **ATP** | Adenosine triphosphate sodium salt (55.1 mg $mL^{-1}$) in $dH_2O$ |
| **$MgCl_2$** | Magnesium chloride anhydrous (47.6 mg $mL^{-1}$) in $dH_2O$ |

**Table S2** Plasmids and protein expression summary

| Internal Name | Product | Vector | Resistance | Expression |
|---|---|---|---|---|
| pACYC-sfp | *Bs*Sfp | pACYC-Duet | CampR | 0.1 mM IPTG, 16 °C, 18 hours |
| pET16b-aass | *Vh*AasS | pET16b | AmpR | 0.5 mM IPTG, 37 °C, 4.5 hours |
| pSGL-005 | *Ec*AcpP (C-term Histag) | pET23a | AmpR | Auto-induction, 30 °C, 18 hours |
| pALGO-013 | ALGO-013 (C-term Histag) | pET28a | KanR | 1 mM IPTG, 18 °C, 18 hours |
| pALGO-023 | ALGO-013 (C-term Histag) | pET28a | KanR | 1 mM IPTG, 18 °C, 18 hours |
| pALGO-040 | ALGO-040 (C-term Histag) | pET28a | KanR | 1 mM IPTG, 18 °C, 18 hours |
| pALGO-044 | ALGO-044 (C-term Histag) | pET28a | KanR | 1 mM IPTG, 18 °C, 18 hours |
| pALGO-055 | ALGO-055 (C-term Histag) | pET28a | KanR | 1 mM IPTG, 18 °C, 18 hours |
| pALGO-057 | ALGO-057 (C-term Histag) | pET28a | KanR | 1 mM IPTG, 18 °C, 18 hours |
| pALGO-059 | ALGO-059 (C-term Histag) | pET28a | KanR | 1 mM IPTG, 18 °C, 18 hours |
| pCHALGO-012 | chALGO-012 (C-term Histag) | pET28a | KanR | 1 mM IPTG, 18 °C, 18 hours |
| pCHALGO-012 | chALGO-024 (C-term Histag) | pET28a | KanR | 1 mM IPTG, 18 °C |

**Table S3** Seed AcpP sequences used for homologue retrieval.

| Organism | Sequence |
|---|---|
| *Escherichia coli* | MSTIEERVKKIIGEQLGVKQEEVTNNASFVEDLGADSLDTVELVMALEEEFDTEIPDEEAEKITTVQ AAIDYINGHQA |
| *Pseudomonas putida* | MSTIEERVKKIVAEQLGVKEEEVTPEKSFVDDLGADSLDTVELVMALEEEFETEIPDEEAEKITTVQ AAIDYVNSHKA |
| *Bacteroides fragilis* | MSEIASRVKAIIVDKLGVEESEVTETASFTNDLGADSLDTVELIMEFEKEFGISIPDDQAEKIGTVQD AIAYIEEHAK |
| *Sphingomonas paucimobilis* | MSETADRVKKIVVEHLGVEADKVTEDASFIDDLGADSLDIVELVMAFEEEFGVEIPDDAAEKITTVK DAITYIDENKA |
| *Salmonella enterica* | MSTIEERVKKIIGEQLGVKQEEVTNNASFVEDLGADSLDTVELVMALEEEFDTEIPDEEAEKITTVQ AAIDYINSHQA |
| *Chlamydia trachomatis* | MSLEDDVKAIIVDQLGVSPEDVKVDSSFIEDLNADSLDLTELIMTLEEKFAFEISEDDAEQLRTVGD VIKYIQEHQN |
| *Helicobacter pylori* | MSLFEDIQAVIAEQLNVDAAQVTPEAEFVKDLGADSLDVVELIMALEEKFGIEIPDEQAEKIVNVGD VVKYIEDNKLA |
| *Legionella pneumophila* | MSTVEERVRKIVVEQLGVKEEELKNDASFVDDLGADSLDTVELVMALEEEFETEIPDEKAEKITTI QEAIDYIESNLNKEEA |
| *Vibrio cholerae* | MSNIEERVKKIIVEQLGVDEAEVKNESSFVEDLGADSLDTVELVMALEEEFDTEIPDEEAEKITTVQ AAIDYVTSNAQ |
| *Neisseria meningitidis* | MSNIEQQVKKIVAEQLGVNEADVKNESSFQDDLGADSLDTVELVMALEEAFGCEIPDEDAEKITT VQLAIDYINTHNG |
| *Acinetobacter baumannii* | MSTIEERVKKIVAEQLGVKEEEVTNSASFVEDLGADSLDTVELVMALEEEFETEIPDEKAEKITTVQ EAIDYIVAHQQ |
| *Campylobacter jejuni* | MATFDDVKAVVVEQLGIDADAVKMESKIIEDLGADSLDVVELIMALEEKFEVEIPDSDAEKLIKIEDV VNYIDNLKK |
| *Porphyromonas gingivalis* | MSEVEKKVIDLVVDKLNVEASEVTREASFSNDLGADSLDTVELMMNFEKEFNMSIPDDQAQEIKT VGDAIDYIEKNLK |
| *Caulobacter vibrioides* | MSDILERVRKIVIEHLDADPEKVTEKASFIDDLGADSLDNVELVMAFEEEFDIEIPDDAAEHIQTVG DAVKFITEKTA |
| *Bordetella parapertussis* | MESIEQRVKKIVAEQLGVNEAEIKNESSFLDDLGADSLDMVELVMALEDEFETEIPDEEAEKITTV QQAVDYINSHGKQ |
| *Burkholderia cepacia* | MDNIEQRVKKIVAEQLGVAEAEIKTEASFVNDLGADSLDTVELVMALEDEFGMEIPDEEAEKITTV QQAIDYARANVKA |
| *Yersinia pestis* | MSTIEERVKKIIVEQLGVKEDEVKNSASFVEDLGADSLDTVELVMALEEEFDTEIPDEEAEKITTVQ AAIDFINANQQ |

**Table S4** Descriptive statistics of sampled ALGO sequences (pI).

| Factor | N | Mean | StDev | 95% CI |
|---|---|---|---|---|
| r = 0.00 | 100 | 4.386 | 0.089 | (4.369, 4.404) |
| r = 0.60 | 100 | 4.320 | 0.106 | (4.299, 4.341) |
| r = 1.00 | 100 | 4.240 | 0.174 | (4.205, 4.274) |

**Table S5** Descriptive statistics of sampled ALGO sequences (GRAVY).

| Factor | N | Mean | StDev | 95% CI |
|---|---|---|---|---|
| r = 0.00 | 100 | -0.201 | 0.137 | (-0.228, -0.173) |
| r = 0.60 | 100 | -0.105 | 0.181 | (-0.141, -0.069) |
| r = 1.00 | 100 | -0.029 | 0.185 | (-0.066, 0.007) |

**Table S6** Descriptive statistics of sampled ALGO sequences (pLDDT).

| Factor | N | Mean | StDev | 95% CI |
|---|---|---|---|---|
| r = 0.00 | 100 | 93.356 | 0.932 | (93.170, 93.540) |
| r = 0.60 | 100 | 90.086 | 2.746 | (89.541, 90.631) |
| r = 1.00 | 100 | 86.823 | 4.216 | (85.986, 87.660) |

**Table S7** Welch’s ANOVA (pI).

| Source | DF Num | DF Den | F-Value | P-Value |
|---|---|---|---|---|
| Factor (pI) | 2 | 187.826 | 31.773 | 1.30E-12 |

**Table S8** Welch’s ANOVA (GRAVY).

| Source | DF Num | DF Den | F-Value | P-Value |
|---|---|---|---|---|
| Factor (GRAVY) | 2 | 193.849 | 28.882 | 1.05E-11 |

**Table S9** Welch’s ANOVA (pLDDT).

| Source | DF Num | DF Den | F-Value | P-Value |
|---|---|---|---|---|
| Factor (pLDDT) | 2 | 150.882 | 166.258 | <1.000 E-16 |

**Table S10** Games-Howell simultaneous tests for differences of means (pI)

| Difference of Levels | Difference of means | SE of Difference | 95% CI | T-Value | Adjusted P-Value |
|---|---|---|---|---|---|
| r = 0.60 – r = 0.00 | -0.066 | 0.014 | (-0.099, -0.034) | -4.790 | 1.963E-05 |
| r = 1.00 – r = 0.00 | -0.146 | 0.020 | (-0.193, -0.100) | -7.508 | 9.951E-06 |
| r = 1.00 - r = 0.60 | -0.080 | 0.020 | (-0.129, -0.032) | -3.951 | 3.485E-04 |

**Table S11** Games-Howell simultaneous tests for differences of means (GRAVY).

| Difference of Levels | Difference of means | SE of Difference | 95% CI | T-Value | Adjusted P-Value |
|---|---|---|---|---|---|
| r = 0.60 – r = 0.00 | 0.095 | 0.023 | (0.042, 0.149) | 4.191 | 1.362E-04 |
| r = 1.00 – r = 0.00 | 0.171 | 0.023 | (0.117, 0.226) | 7.437 | 9.951E-06 |
| r = 1.00 - r = 0.60 | 0.076 | 0.026 | (0.015, 0.137) | 2.933 | 1.045E-02 |

**Table S12** Games-Howell simultaneous tests for differences of means (pLDDT).

| Difference of Levels | Difference of means | SE of Difference | 95% CI | T-Value | Adjusted P-Value |
|---|---|---|---|---|---|
| r = 0.60 – r = 0.00 | -3.269 | 0.290 | (-3.958, -2.581) | -11.28 | 9.951E-06 |
| r = 1.00 – r = 0.00 | -6.533 | 0.432 | (-7.558, -5.507) | -15.13 | 9.951E-06 |
| r = 1.00 - r = 0.60 | -3.263 | 0.503 | (-4.452, -2.075) | -6.49 | 9.953E-06 |

**Table S13** ALGO sequences sampled for experimental testing.

| Protein | Sequence | %ID (vs *Ec*AcpP) | GRAVY | pI | pLDDT |
|---|---|---|---|---|---|
| ALGO-013 | MSTLPDKVKKIVADQLGVKEPKVKNHASFIQDLGADSLDTVELVMSMEEDFDVEIPDDTAEKISSVAQAIDYVTDHSA | 64.10 | -0.25 | 4.41 | 93.96 |
| ALGO-023 | MMSTNDKVEKIVLNELGVGETEVQLDAIFVDDLGADSLDSVELIMTLENSFDIQIPDDHAETITTVQFAVSYATAKSK | 51.28 | 0.03 | 4.30 | 94.02 |
| ALGO-040 | MNDVSQRVQRVLGEQLGVVETEVVTSAAFVEDLGADSLDTVELVMSLEEAFNFNIPDEHAEDITQVQSAVHYINEQSA | 61.54 | -0.08 | 4.20 | 94.72 |
| ALGO-044 | MSEIIKKVKKIVVQELGVQVNEVHESSAFVQDLGADSLDFVELIMSFEEQFHTAIPDEDAEKIITVAQAIHYINNNSG | 57.69 | -0.01 | 4.54 | 94.26 |
| ALGO-055 | MNPLEQRVKTIIVQELGVNEDVVINDASFVRDLGADSLDSVELVMALEKEFSIQIPDEQAEKIIQVSAAIDYAEKAAK | 61.54 | 0.00 | 4.46 | 93.52 |
| ALGO-057 | MNELFRKVQAIVVENLGVEGVKVTNESAFANDLGADSLDQVELLMAIEEQFDCDIPDEEAEDIITVADAILYIGASSQ | 53.85 | 0.13 | 4.33 | 93.91 |
| ALGO-059 | MSNLDQRVIDIIVQELGVPPKEVKSEASFIKDLGADSLDTVELIMSIEEDFNVEIPDEDAEHITTVASVLNYLNEHSN | 56.41 | -0.15 | 4.27 | 93.74 |
| $^{ch}$ALGO-009 | MNPLDQRVITIIVQELGVPPKEVKSDASFVKDLGADSLDTVELIMSIEKEFSVQIPDEDAEHITTVSAVLDYANKHAN | 57.69 | -0.10 | 4.18 | 92.71 |
| $^{ch}$ALGO-012 | MSNLDQRVIDIIVQELGVNPDVVKSEASFIKDLGADSLDTVELIMSIEKDFSIQIPDEDAEHIIQVSAALNYAEEHAN | 51.28 | -0.03 | 3.84 | 94.48 |
| $^{ch}$ALGO-024 | MSPLDQRVIDIIVQELGVNPDVVKSEASFIKDLGADSLDTVELIMALEKDFSIQIPDEDAEKIITVAAVIDYLNEHAN | 58.97 | 0.14 | 3.82 | 94.27 |
| $^{ch}$ALGO-044 | MNNLEQRVKDIIVQELGVNEDEVISDASFIKDLGADSLDTVELIMALEEDFNVEIPDEDAEKITTVAAAINYAEKHSN | 64.10 | -0.235 | 3.78 | 93.56 |
| $^{ch}$ALGO-097 | MSNLEQRVKTIIVQELGVPEKVVINDASFVKDLGADSLDTVELIMSLEKEFNIEIPDEDAEHIITVASVINYANKASK | 60.26 | 0.013 | 4.26 | 91.85 |

**Table S14** LC/ESI-MS (*Ec*AcpP and ALGO sequences).

| Protein | Empirical Formula | Theoretical MW / Da | Actual MW / Da | Comment |
|---|---|---|---|---|
| *apo-Ec*AcpP | $C_{420}H_{658}N_{114}O_{146}S_{2}$ | 9704.50 | 9573.39 ± 0.09 | -$Met_1$ |
| *holo-Ec*AcpP | $C_{431}H_{679}N_{116}O_{152}S_{3}P$ | 10044.83 | 9913.78 ± 0.04 | -$Met_1$ |
| $C_{12}$-*Ec*AcpP | $C_{443}H_{702}N_{116}O_{153}S_{3}P$ | 10228.14 | 10096.08 ± 0.14 | -$Met_1$ |
| *apo*-ALGO-013 | $C_{418}H_{657}N_{113}O_{139}S_{3}$ | 9585.53 | 9454.33 ± 0.02 | -$Met_1$ |
| *apo*-ALGO-040 | $C_{417}H_{642}N_{118}O_{142}S_{2}$ | 9644.37 | 9512.88 ± 0.25<br>9644.40 ± 0.18 | Partial loss of $Met_1$ |
| *apo*-ALGO-055 | $C_{425}H_{679}N_{117}O_{136}S_{2}$ | 9667.74 | 9667.43 ± 0.00 | |
| *holo*-ALGO-055 | $C_{436}H_{700}N_{119}O_{142}S_{3}P$ | 10008.08 | 10007.91 ± 0.12 | |
| $C_{12}$-ALGO-055 | $C_{448}H_{722}N_{119}O_{143}S_{3}P$ | 10190.38 | 10190.28 ± 0.00 | |
| *apo*-ALGO-059 | $C_{426}H_{665}N_{115}O_{143}S_{2}$ | 9749.63 | 9618.46 ± 0.12 | -$Met_1$ |
| *holo*-ALGO-059 | $C_{437}H_{686}N_{117}O_{149}S_{3}P$ | 10089.96 | 9958.44 ± 0.13 | -$Met_1$ |
| $C_{12}$-ALGO-059 | $C_{449}H_{708}N_{117}O_{150}S_{3}P$ | 10272.26 | 10140.79 ± 0.12 | -$Met_1$ |
| *apo*-$^{ch}$ALGO-012 | $C_{423}H_{662}N_{116}O_{140}S_{2}$ | 9676.64 | 9545.40 ± 0.02 | -$Met_1$ |
| *holo*- $^{ch}$ALGO-012 | $C_{434}H_{683}N_{118}O_{146}S_{3}P$ | 10016.91 | 9885.66 ± 0.01 | -$Met_1$ |
| $C_{12}$- $^{ch}$ALGO-012 | $C_{446}H_{704}N_{118}O_{147}S_{3}P$ | 10198.21 | 10067.91 ± 0.02 | -$Met_1$ |
| *apo*- $^{ch}$ALGO-024 | $C_{427}H_{675}N_{113}O_{137}S_{2}$ | 9647.76 | 9647.70 ± 0.02 | |
| *holo*- $^{ch}$ALGO-024 | $C_{438}H_{696}N_{115}O_{143}S_{3}P$ | 9988.04 | 9987.96 ± 0.02 | |
| $C_{12}$- $^{ch}$ALGO-024 | $C_{450}H_{718}N_{115}O_{144}S_{3}P$ | 10170.34 | 10170.16 ± 0.04 | |

**Table S15** Disordered-ordered classification of proteins studied by CD spectroscopy. Values were computed using BeStSel server using mean residue ellipticity as input.

| Protein | PTM | 197 nm | 206 nm | 233 nm | Prediction |
|---|---|---|---|---|---|
| *Ec*AcpP | *apo-* | 3.89 | -3.33 | -1.52 | ordered |
| | *holo-* | 3.77 | -3.11 | -1.23 | ordered |
| | $C_{12}$- | 5.79 | -4.16 | -1.74 | ordered |
| ALGO-055 | *apo-* | -4.01 | -5.19 | -1.58 | disordered |
| | *holo-* | -2.25 | -2.73 | -0.73 | disordered |
| | $C_{12}$- | 1.37 | -4.16 | -1.66 | ordered |
| ALGO-059 | *apo-* | -4.46 | -4.38 | -1.11 | disordered |
| | *holo-* | -2.74 | -2.66 | -0.66 | disordered |
| | $C_{12}$- | 1.38 | -4.61 | -1.62 | ordered |

**Table S16** Estimated percent helical content of proteins studied by CD spectroscopy across PTM states. Values were computed using BeStSel server using mean residue ellipticity as input.

| Protein | ***apo-*** | ***holo-*** | **$C_{12}$-** |
|---|---|---|---|
| *Ec*AcpP | 28.1 | 22.2 | 35.0 |
| ALGO-055 | 15.6 | 5.8 | 21.1 |
| ALGO-059 | 12.2 | 4.0 | 28.2 |

**Table S17** Positional analysis of ALGO-055 with comparison to *Ec*AcpP.

| Position | Residue | vdW / Å | $H_{KD}$ | pI | Rarity (vs input MSA) | Equivalent (*Ec*AcpP) | BLOSUM62 (vs. *Ec*AcpP) | Grantham Distance (vs. *Ec*AcpP) |
|---|---|---|---|---|---|---|---|---|
| 1 | M | 162.9 | 1.90 | 5.74 | 0.749 | M | 5 | 0 |
| 2 | N | 122.4 | -3.50 | 5.41 | 0.012 | S | 1 | 46 |
| 3 | P | 121.6 | -1.60 | 6.30 | 0.001 | T | -1 | 38 |
| 4 | L | 163.1 | 3.80 | 5.98 | 0.072 | I | 2 | 5 |
| 5 | E | 138.8 | -3.50 | 3.22 | 0.585 | E | 5 | 0 |
| 6 | Q | 146.9 | -3.50 | 5.65 | 0.162 | E | 2 | 29 |
| 7 | R | 190.3 | -4.50 | 10.76 | 0.728 | R | 5 | 0 |
| 8 | V | 138.2 | 4.20 | 5.96 | 0.933 | V | 4 | 0 |
| 9 | K | 165.1 | -3.90 | 9.74 | 0.814 | K | 5 | 0 |
| 10 | T | 119.6 | -0.70 | 5.60 | 0.007 | K | -1 | 78 |
| 11 | I | 163.0 | 4.50 | 6.02 | 0.898 | I | 4 | 0 |
| 12 | I | 163.0 | 4.50 | 6.02 | 0.394 | I | 4 | 0 |
| 13 | V | 138.2 | 4.20 | 5.96 | 0.545 | G | -3 | 109 |
| 14 | Q | 146.9 | -3.50 | 5.65 | 0.004 | E | 2 | 29 |
| 15 | E | 138.8 | -3.50 | 3.22 | 0.004 | Q | 2 | 29 |
| 16 | L | 163.1 | 3.80 | 5.98 | 0.996 | L | 4 | 0 |
| 17 | G | 63.8 | -0.40 | 5.97 | 0.876 | G | 6 | 0 |
| 18 | V | 138.2 | 4.20 | 5.96 | 0.915 | V | 4 | 0 |
| 19 | N | 122.4 | -3.50 | 5.41 | 0.142 | K | 0 | 94 |
| 20 | E | 138.8 | -3.50 | 3.22 | 0.655 | Q | 2 | 29 |
| 21 | D | 114.4 | -3.50 | 2.77 | 0.234 | E | 2 | 45 |
| 22 | V | 138.2 | 4.20 | 5.96 | 0.001 | E | -2 | 121 |
| 23 | V | 138.2 | 4.20 | 5.96 | 0.867 | V | 4 | 0 |
| 24 | I | 163.0 | 4.50 | 6.02 | 0.012 | T | -1 | 89 |
| 25 | N | 122.4 | -3.50 | 5.41 | 0.376 | N | 6 | 0 |
| 26 | D | 114.4 | -3.50 | 2.77 | 0.100 | N | 1 | 23 |
| 27 | A | 89.3 | 1.80 | 6.00 | 0.652 | A | 4 | 0 |
| 28 | S | 89.0 | -0.80 | 5.68 | 0.838 | S | 4 | 0 |
| 29 | F | 190.8 | 2.80 | 5.48 | 0.983 | F | 6 | 0 |
| 30 | V | 138.2 | 4.20 | 5.96 | 0.619 | V | 4 | 0 |
| 31 | R | 190.3 | -4.50 | 10.76 | 0.001 | E | 0 | 54 |
| 32 | D | 114.4 | -3.50 | 2.77 | 0.999 | D | 6 | 0 |
| 33 | L | 163.1 | 3.80 | 5.98 | 0.999 | L | 4 | 0 |
| 34 | G | 63.8 | -0.40 | 5.97 | 0.957 | G | 6 | 0 |
| 35 | A | 89.3 | 1.80 | 6.00 | 0.984 | A | 4 | 0 |
| 36 | D | 114.4 | -3.50 | 2.77 | 0.999 | D | 6 | 0 |
| 37 | S | 89.0 | -0.80 | 5.68 | 0.999 | S | 4 | 0 |
| 38 | L | 163.1 | 3.80 | 5.98 | 0.999 | L | 4 | 0 |
| 39 | D | 114.4 | -3.50 | 2.77 | 1.000 | D | 6 | 0 |
| 40 | S | 89.0 | -0.80 | 5.68 | 0.001 | T | 1 | 58 |
| 41 | V | 138.2 | 4.20 | 5.96 | 0.971 | V | 4 | 0 |
| 42 | E | 138.8 | -3.50 | 3.22 | 1.000 | E | 5 | 0 |
| 43 | L | 163.1 | 3.80 | 5.98 | 0.992 | L | 4 | 0 |
| 44 | V | 138.2 | 4.20 | 5.96 | 0.784 | V | 4 | 0 |
| 45 | M | 162.9 | 1.90 | 5.74 | 0.998 | M | 5 | 0 |

| | | | | | | | | |
|---|---|---|---|---|---|---|---|---|
| 46 | A | 89.3 | 1.80 | 6.00 | 0.848 | A | 4 | 0 |
| 47 | L | 163.1 | 3.80 | 5.98 | 0.695 | L | 4 | 0 |
| 48 | E | 138.8 | -3.50 | 3.22 | 0.998 | E | 5 | 0 |
| 49 | K | 165.1 | -3.90 | 9.74 | 0.113 | E | 1 | 56 |
| 50 | E | 138.8 | -3.50 | 3.22 | 0.850 | E | 5 | 0 |
| 51 | F | 190.8 | 2.80 | 5.48 | 0.996 | F | 6 | 0 |
| 52 | S | 89.0 | -0.80 | 5.68 | 0.025 | D | 0 | 65 |
| 53 | I | 163.0 | 4.50 | 6.02 | 0.258 | T | -1 | 89 |
| 54 | Q | 146.9 | -3.50 | 5.65 | 0.005 | E | 2 | 29 |
| 55 | I | 163.0 | 4.50 | 6.02 | 0.987 | I | 4 | 0 |
| 56 | P | 121.6 | -1.60 | 6.30 | 0.950 | P | 7 | 0 |
| 57 | D | 114.4 | -3.50 | 2.77 | 0.973 | D | 6 | 0 |
| 58 | E | 138.8 | -3.50 | 3.22 | 0.696 | E | 5 | 0 |
| 59 | Q | 146.9 | -3.50 | 5.65 | 0.153 | E | 2 | 29 |
| 60 | A | 89.3 | 1.80 | 6.00 | 0.991 | A | 4 | 0 |
| 61 | E | 138.8 | -3.50 | 3.22 | 0.984 | E | 5 | 0 |
| 62 | K | 165.1 | -3.90 | 9.74 | 0.841 | K | 5 | 0 |
| 63 | I | 163.0 | 4.50 | 6.02 | 0.910 | I | 4 | 0 |
| 64 | I | 163.0 | 4.50 | 6.02 | 0.018 | T | -1 | 89 |
| 65 | Q | 146.9 | -3.50 | 5.65 | 0.001 | T | -1 | 42 |
| 66 | V | 138.2 | 4.20 | 5.96 | 0.948 | V | 4 | 0 |
| 67 | S | 89.0 | -0.80 | 5.68 | 0.009 | Q | 0 | 68 |
| 68 | A | 89.3 | 1.80 | 6.00 | 0.142 | A | 4 | 0 |
| 69 | A | 89.3 | 1.80 | 6.00 | 0.869 | A | 4 | 0 |
| 70 | I | 163.0 | 4.50 | 6.02 | 0.669 | I | 4 | 0 |
| 71 | D | 114.4 | -3.50 | 2.77 | 0.559 | D | 6 | 0 |
| 72 | Y | 194.6 | -1.30 | 5.66 | 0.832 | Y | 7 | 0 |
| 73 | A | 89.3 | 1.80 | 6.00 | 0.018 | I | -1 | 94 |
| 74 | E | 138.8 | -3.50 | 3.22 | 0.309 | N | 0 | 42 |
| 75 | K | 165.1 | -3.90 | 9.74 | 0.108 | G | -2 | 127 |
| 76 | A | 89.3 | 1.80 | 6.00 | 0.030 | H | -2 | 86 |
| 77 | A | 89.3 | 1.80 | 6.00 | 0.079 | Q | -1 | 91 |
| 78 | K | 165.1 | -3.90 | 9.74 | 0.137 | A | -1 | 106 |

**Table S18** Positional analysis of ALGO-059 with comparison to *Ec*AcpP.

| Position | Residue | vdW / Å | $H_{KD}$ | pI | Rarity (vs input MSA) | Equivalent (*Ec*AcpP) | BLOSUM62 (vs. *Ec*AcpP) | Grantham Distance (vs. *Ec*AcpP) |
|---|---|---|---|---|---|---|---|---|
| 1 | M | 162.9 | 1.90 | 5.74 | 0.749 | M | 5 | 0 |
| 2 | S | 89.0 | -0.80 | 5.68 | 0.624 | S | 4 | 0 |
| 3 | N | 122.4 | -3.50 | 5.41 | 0.206 | T | 0 | 65 |
| 4 | L | 163.1 | 3.80 | 5.98 | 0.072 | I | 2 | 5 |
| 5 | D | 114.4 | -3.50 | 2.77 | 0.016 | E | 2 | 45 |
| 6 | Q | 146.9 | -3.50 | 5.65 | 0.162 | E | 2 | 29 |
| 7 | R | 190.3 | -4.50 | 10.76 | 0.728 | R | 5 | 0 |
| 8 | V | 138.2 | 4.20 | 5.96 | 0.933 | V | 4 | 0 |
| 9 | I | 163.0 | 4.50 | 6.02 | 0.032 | K | -3 | 102 |
| 10 | D | 114.4 | -3.50 | 2.77 | 0.050 | K | -1 | 101 |
| 11 | I | 163.0 | 4.50 | 6.02 | 0.898 | I | 4 | 0 |
| 12 | I | 163.0 | 4.50 | 6.02 | 0.394 | I | 4 | 0 |
| 13 | V | 138.2 | 4.20 | 5.96 | 0.545 | G | -3 | 109 |
| 14 | Q | 146.9 | -3.50 | 5.65 | 0.004 | E | 2 | 29 |
| 15 | E | 138.8 | -3.50 | 3.22 | 0.004 | Q | 2 | 29 |
| 16 | L | 163.1 | 3.80 | 5.98 | 0.996 | L | 4 | 0 |
| 17 | G | 63.8 | -0.40 | 5.97 | 0.876 | G | 6 | 0 |
| 18 | V | 138.2 | 4.20 | 5.96 | 0.915 | V | 4 | 0 |
| 19 | P | 121.6 | -1.60 | 6.30 | 0.006 | K | -1 | 103 |
| 20 | P | 121.6 | -1.60 | 6.30 | 0.084 | Q | -1 | 76 |
| 21 | K | 165.1 | -3.90 | 9.74 | 0.013 | E | 1 | 56 |
| 22 | E | 138.8 | -3.50 | 3.22 | 0.611 | E | 5 | 0 |
| 23 | V | 138.2 | 4.20 | 5.96 | 0.867 | V | 4 | 0 |
| 24 | K | 165.1 | -3.90 | 9.74 | 0.347 | T | -1 | 78 |
| 25 | S | 89.0 | -0.80 | 5.68 | 0.047 | N | 1 | 46 |
| 26 | E | 138.8 | -3.50 | 3.22 | 0.568 | N | 0 | 42 |
| 27 | A | 89.3 | 1.80 | 6.00 | 0.652 | A | 4 | 0 |
| 28 | S | 89.0 | -0.80 | 5.68 | 0.838 | S | 4 | 0 |
| 29 | F | 190.8 | 2.80 | 5.48 | 0.983 | F | 6 | 0 |
| 30 | I | 163.0 | 4.50 | 6.02 | 0.241 | V | 3 | 29 |
| 31 | K | 165.1 | -3.90 | 9.74 | 0.032 | E | 1 | 56 |
| 32 | D | 114.4 | -3.50 | 2.77 | 0.999 | D | 6 | 0 |
| 33 | L | 163.1 | 3.80 | 5.98 | 0.999 | L | 4 | 0 |
| 34 | G | 63.8 | -0.40 | 5.97 | 0.957 | G | 6 | 0 |
| 35 | A | 89.3 | 1.80 | 6.00 | 0.984 | A | 4 | 0 |
| 36 | D | 114.4 | -3.50 | 2.77 | 0.999 | D | 6 | 0 |
| 37 | S | 89.0 | -0.80 | 5.68 | 0.999 | S | 4 | 0 |
| 38 | L | 163.1 | 3.80 | 5.98 | 0.999 | L | 4 | 0 |
| 39 | D | 114.4 | -3.50 | 2.77 | 1.000 | D | 6 | 0 |
| 40 | T | 119.6 | -0.70 | 5.60 | 0.784 | T | 5 | 0 |
| 41 | V | 138.2 | 4.20 | 5.96 | 0.971 | V | 4 | 0 |
| 42 | E | 138.8 | -3.50 | 3.22 | 1.000 | E | 5 | 0 |
| 43 | L | 163.1 | 3.80 | 5.98 | 0.992 | L | 4 | 0 |
| 44 | I | 163.0 | 4.50 | 6.02 | 0.209 | V | 3 | 29 |

| | | | | | | | | |
|---|---|---|---|---|---|---|---|---|
| 45 | M | 162.9 | 1.90 | 5.74 | 0.998 | M | 5 | 0 |
| 46 | S | 89.0 | -0.80 | 5.68 | 0.007 | A | 1 | 99 |
| 47 | I | 163.0 | 4.50 | 6.02 | 0.001 | L | 2 | 5 |
| 48 | E | 138.8 | -3.50 | 3.22 | 0.998 | E | 5 | 0 |
| 49 | E | 138.8 | -3.50 | 3.22 | 0.845 | E | 5 | 0 |
| 50 | D | 114.4 | -3.50 | 2.77 | 0.001 | E | 2 | 45 |
| 51 | F | 190.8 | 2.80 | 5.48 | 0.996 | F | 6 | 0 |
| 52 | N | 122.4 | -3.50 | 5.41 | 0.112 | D | 1 | 23 |
| 53 | V | 138.2 | 4.20 | 5.96 | 0.119 | T | 0 | 69 |
| 54 | E | 138.8 | -3.50 | 3.22 | 0.869 | E | 5 | 0 |
| 55 | I | 163.0 | 4.50 | 6.02 | 0.987 | I | 4 | 0 |
| 56 | P | 121.6 | -1.60 | 6.30 | 0.950 | P | 7 | 0 |
| 57 | D | 114.4 | -3.50 | 2.77 | 0.973 | D | 6 | 0 |
| 58 | E | 138.8 | -3.50 | 3.22 | 0.696 | E | 5 | 0 |
| 59 | D | 114.4 | -3.50 | 2.77 | 0.186 | E | 2 | 45 |
| 60 | A | 89.3 | 1.80 | 6.00 | 0.991 | A | 4 | 0 |
| 61 | E | 138.8 | -3.50 | 3.22 | 0.984 | E | 5 | 0 |
| 62 | H | 157.5 | -3.20 | 7.59 | 0.012 | K | -1 | 32 |
| 63 | I | 163.0 | 4.50 | 6.02 | 0.910 | I | 4 | 0 |
| 64 | T | 119.6 | -0.70 | 5.60 | 0.548 | T | 5 | 0 |
| 65 | T | 119.6 | -0.70 | 5.60 | 0.892 | T | 5 | 0 |
| 66 | V | 138.2 | 4.20 | 5.96 | 0.948 | V | 4 | 0 |
| 67 | A | 89.3 | 1.80 | 6.00 | 0.010 | Q | -1 | 91 |
| 68 | S | 89.0 | -0.80 | 5.68 | 0.032 | A | 1 | 99 |
| 69 | V | 138.2 | 4.20 | 5.96 | 0.124 | A | 0 | 64 |
| 70 | L | 163.1 | 3.80 | 5.98 | 0.010 | I | 2 | 5 |
| 71 | N | 122.4 | -3.50 | 5.41 | 0.057 | D | 1 | 23 |
| 72 | Y | 194.6 | -1.30 | 5.66 | 0.832 | Y | 7 | 0 |
| 73 | L | 163.1 | 3.80 | 5.98 | 0.056 | I | 2 | 5 |
| 74 | N | 122.4 | -3.50 | 5.41 | 0.250 | N | 6 | 0 |
| 75 | E | 138.8 | -3.50 | 3.22 | 0.164 | G | -2 | 98 |
| 76 | H | 157.5 | -3.20 | 7.59 | 0.342 | H | 8 | 0 |
| 77 | S | 89.0 | -0.80 | 5.68 | 0.050 | Q | 0 | 68 |
| 78 | N | 122.4 | -3.50 | 5.41 | 0.009 | A | -2 | 111 |

128 **Table S19** Positional comparison between ALGO-055 and ALGO-059

| Position | ALGO-055 | ALGO-059 | BLOSUM62 | Grantham Distance |
|---|---|---|---|---|
| 1 | M | M | 5 | 0 |
| 2 | N | S | 1 | 46 |
| 3 | P | N | -2 | 91 |
| 4 | L | L | 4 | 0 |
| 5 | E | D | 2 | 45 |
| 6 | Q | Q | 5 | 0 |
| 7 | R | R | 5 | 0 |
| 8 | V | V | 4 | 0 |
| 9 | K | I | -3 | 102 |
| 10 | T | D | -1 | 85 |
| 11 | I | I | 4 | 0 |
| 12 | I | I | 4 | 0 |
| 13 | V | V | 4 | 0 |
| 14 | Q | Q | 5 | 0 |
| 15 | E | E | 5 | 0 |
| 16 | L | L | 4 | 0 |
| 17 | G | G | 6 | 0 |
| 18 | V | V | 4 | 0 |
| 19 | N | P | -2 | 91 |
| 20 | E | P | -1 | 93 |
| 21 | D | K | -1 | 101 |
| 22 | V | E | -2 | 121 |
| 23 | V | V | 4 | 0 |
| 24 | I | K | -3 | 102 |
| 25 | N | S | 1 | 46 |
| 26 | D | E | 2 | 45 |
| 27 | A | A | 4 | 0 |
| 28 | S | S | 4 | 0 |
| 29 | F | F | 6 | 0 |
| 30 | V | I | 3 | 29 |
| 31 | R | K | 2 | 26 |
| 32 | D | D | 6 | 0 |
| 33 | L | L | 4 | 0 |
| 34 | G | G | 6 | 0 |
| 35 | A | A | 4 | 0 |
| 36 | D | D | 6 | 0 |
| 37 | S | S | 4 | 0 |
| 38 | L | L | 4 | 0 |
| 39 | D | D | 6 | 0 |
| 40 | S | T | 1 | 58 |
| 41 | V | V | 4 | 0 |
| 42 | E | E | 5 | 0 |
| 43 | L | L | 4 | 0 |
| 44 | V | I | 3 | 29 |
| 45 | M | M | 5 | 0 |

| | | | | |
|---|---|---|---|---|
| 46 | A | S | 1 | 99 |
| 47 | L | I | 2 | 5 |
| 48 | E | E | 5 | 0 |
| 49 | K | E | 1 | 56 |
| 50 | E | D | 2 | 45 |
| 51 | F | F | 6 | 0 |
| 52 | S | N | 1 | 46 |
| 53 | I | V | 3 | 29 |
| 54 | Q | E | 2 | 29 |
| 55 | I | I | 4 | 0 |
| 56 | P | P | 7 | 0 |
| 57 | D | D | 6 | 0 |
| 58 | E | E | 5 | 0 |
| 59 | Q | D | 0 | 61 |
| 60 | A | A | 4 | 0 |
| 61 | E | E | 5 | 0 |
| 62 | K | H | -1 | 32 |
| 63 | I | I | 4 | 0 |
| 64 | I | T | -1 | 89 |
| 65 | Q | T | -1 | 42 |
| 66 | V | V | 4 | 0 |
| 67 | S | A | 1 | 99 |
| 68 | A | S | 1 | 99 |
| 69 | A | V | 0 | 64 |
| 70 | I | L | 2 | 5 |
| 71 | D | N | 1 | 23 |
| 72 | Y | Y | 7 | 0 |
| 73 | A | L | -1 | 96 |
| 74 | E | N | 0 | 42 |
| 75 | K | E | 1 | 56 |
| 76 | A | H | -2 | 86 |
| 77 | A | S | 1 | 99 |
| 78 | K | N | 0 | 94 |

**Table S20** Analysis of amino acid variants found exclusively in unsuccessful ALGO sequences.

| Position | Variant | Count | Rarity (vs input MSA) | Equivalent (*Ec*AcpP) | BLOSUM62 (vs *Ec*AcpP) | Grantham Distance (vs *Ec*AcpP) |
|---|---|---|---|---|---|---|
| 2 | M | 1 | 0.1066 | S | -1 | 135 |
| 3 | E | 2 | 0.0711 | T | 1 | 58 |
| | S | 1 | 0.1703 | | -1 | 65 |
| | D | 1 | 0.2303 | | -1 | 85 |
| 4 | T | 1 | 0.0531 | I | -1 | 89 |
| | V | 1 | 0.2109 | | 3 | 29 |
| 5 | P | 1 | 0.0005 | E | -3 | 140 |
| | N | 1 | 0.0028 | | -3 | 134 |
| | I | 1 | 0.0078 | | 0 | 80 |
| | S | 1 | 0.0134 | | 0 | 42 |
| | F | 1 | 0.0964 | | -1 | 93 |
| 6 | R | 1 | 0.0005 | E | 1 | 56 |
| | K | 1 | 0.0166 | | 2 | 45 |
| | D | 2 | 0.1657 | | 0 | 54 |
| 7 | K | 4 | 0.1357 | R | 2 | 26 |
| 9 | E | 1 | 0.0009 | K | 1 | 56 |
| | Q | 2 | 0.0249 | | 1 | 53 |
| 10 | R | 1 | 0.0078 | K | -1 | 106 |
| | A | 1 | 0.1269 | | 2 | 26 |
| 11 | V | 1 | 0.0868 | I | 3 | 29 |
| 12 | L | 1 | 0.0005 | I | 3 | 29 |
| | V | 4 | 0.5976 | | 2 | 5 |
| 13 | L | 1 | 0.0037 | G | 0 | 60 |
| | A | 1 | 0.2903 | | -4 | 138 |
| 14 | N | 1 | 0.0028 | E | 0 | 42 |
| | D | 1 | 0.1375 | | 2 | 45 |
| 15 | N | 1 | 0.0014 | Q | 0 | 46 |
| 19 | V | 1 | 0.0005 | K | 1 | 53 |
| | G | 1 | 0.0032 | | -2 | 97 |
| | Q | 1 | 0.0037 | | -2 | 127 |
| | E | 1 | 0.1592 | | 1 | 56 |
| 20 | G | 1 | 0.0037 | Q | -2 | 87 |
| | V | 1 | 0.0212 | | -2 | 96 |
| 21 | P | 1 | 0.0014 | E | -1 | 65 |
| | V | 1 | 0.0023 | | 0 | 42 |
| | T | 2 | 0.0171 | | -1 | 93 |
| | N | 1 | 0.0268 | | -2 | 121 |
| 22 | K | 2 | 0.1384 | E | 1 | 56 |
| 24 | H | 1 | 0.0018 | T | -1 | 42 |
| | Q | 1 | 0.0171 | | 0 | 69 |
| | V | 1 | 0.066 | | -2 | 47 |
| 25 | T | 1 | 0.042 | N | 0 | 65 |
| | L | 1 | 0.0452 | | 0 | 42 |
| | E | 1 | 0.1343 | | -3 | 153 |
| 26 | H | 1 | 0.0046 | N | 1 | 46 |

| | | | | | | |
|---|---|---|---|---|---|---|
| | S | 2 | 0.1029 | | 1 | 68 |
| 27 | S | 2 | 0.3096 | A | 1 | 99 |
| 28 | I | 1 | 0.0009 | S | 1 | 99 |
| | A | 3 | 0.012 | | -2 | 142 |
| 30 | A | 1 | 0.006 | V | 0 | 64 |
| 31 | Q | 2 | 0.0014 | E | 0 | 42 |
| | N | 1 | 0.1444 | | 2 | 45 |
| | D | 1 | 0.1444 | | 2 | 29 |
| 40 | F | 1 | 0.0005 | T | -2 | 103 |
| | Q | 1 | 0.0028 | | -1 | 42 |
| 44 | L | 1 | 0.0023 | V | 1 | 32 |
| 46 | T | 1 | 0.0245 | A | 0 | 58 |
| 47 | M | 1 | 0.0111 | L | 0 | 22 |
| | F | 1 | 0.2921 | | 2 | 15 |
| 49 | N | 1 | 0.0009 | E | 0 | 42 |
| 50 | S | 1 | 0.0005 | E | -1 | 107 |
| | Q | 2 | 0.0097 | | 0 | 80 |
| | A | 1 | 0.0208 | | 2 | 29 |
| 52 | H | 1 | 0.0009 | D | -1 | 81 |
| 53 | F | 1 | 0.0175 | T | -1 | 149 |
| | C | 1 | 0.1957 | | -2 | 103 |
| 54 | N | 1 | 0.0032 | E | -1 | 107 |
| | D | 1 | 0.0065 | | 0 | 42 |
| | A | 1 | 0.0443 | | 2 | 45 |
| 58 | D | 2 | 0.2635 | E | 2 | 45 |
| 59 | T | 1 | 0.0005 | E | -1 | 65 |
| | H | 2 | 0.0042 | | 0 | 40 |
| 62 | D | 2 | 0.0014 | K | -1 | 78 |
| | T | 1 | 0.0554 | | -1 | 101 |
| 64 | S | 1 | 0.0535 | T | 1 | 58 |
| 65 | S | 1 | 0.0475 | T | 1 | 58 |
| 68 | F | 1 | 0.0018 | A | -2 | 113 |
| | Q | 2 | 0.2843 | | -2 | 126 |
| | D | 1 | 0.3899 | | -1 | 91 |
| 70 | V | 2 | 0.2796 | I | 3 | 29 |
| 71 | L | 1 | 0.0005 | D | 0 | 65 |
| | H | 2 | 0.0009 | | -1 | 81 |
| | S | 1 | 0.0655 | | -4 | 172 |
| 73 | V | 1 | 0.2635 | I | 3 | 29 |
| 74 | G | 1 | 0.0078 | N | 0 | 65 |
| | T | 2 | 0.0951 | | 0 | 80 |
| 75 | D | 1 | 0.0406 | G | 0 | 60 |
| | N | 1 | 0.084 | | 0 | 80 |
| | A | 2 | 0.3009 | | -1 | 94 |
| 76 | S | 1 | 0.0129 | H | -1 | 89 |
| | Q | 1 | 0.0152 | | -1 | 32 |
| | K | 1 | 0.0452 | | 1 | 68 |

| | | | | | | |
|---|---|---|---|---|---|---|
| | N | 1 | 0.2884 | | 0 | 24 |
| 78 | G | 1 | 0.0097 | A | 0 | 60 |
| | Q | 1 | 0.0125 | | -1 | 91 |

#### Sequence Analysis (Fig. S1-3)

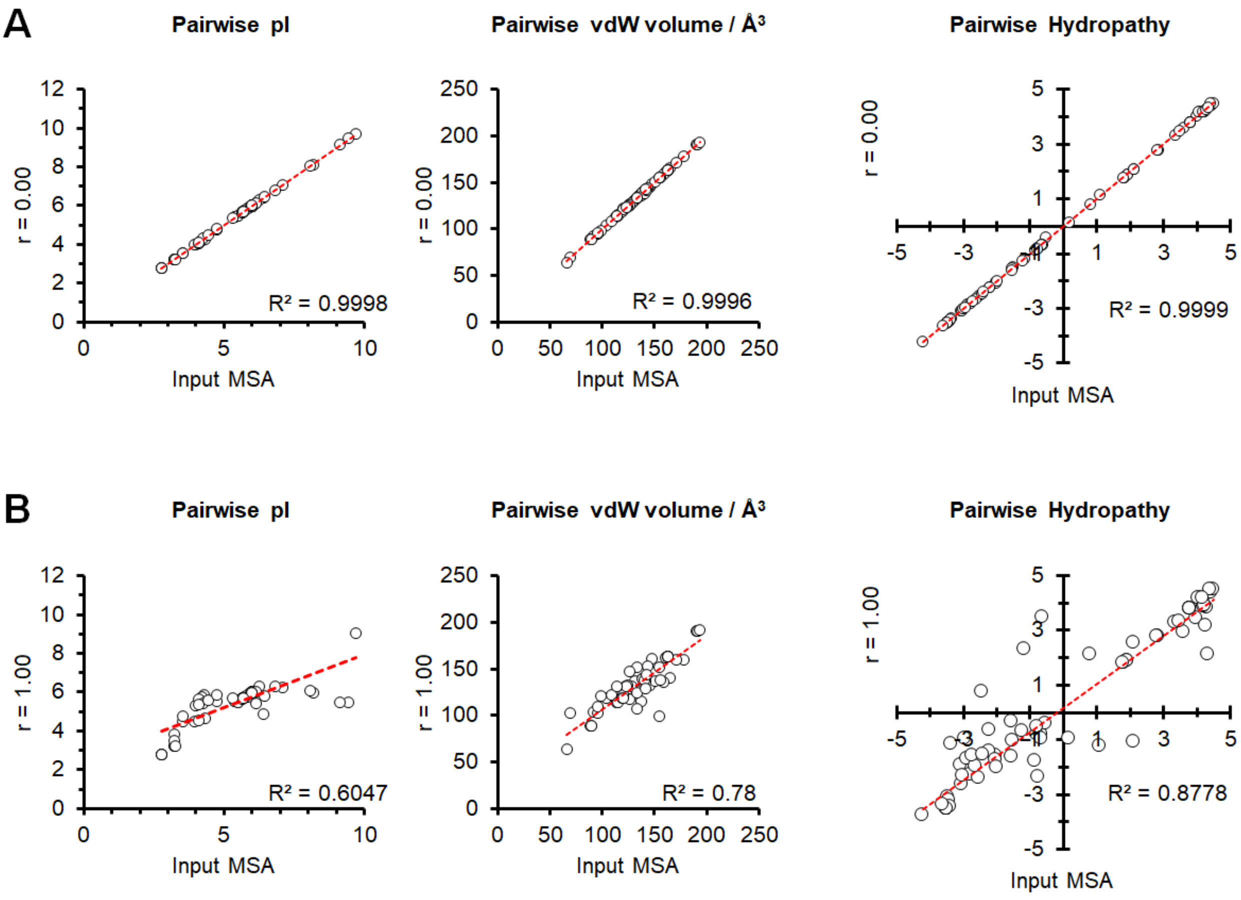


**Fig. S1** Example pairwise correlation plots for sequence sets r = 0.00 (A) and r = 1.00 (B).

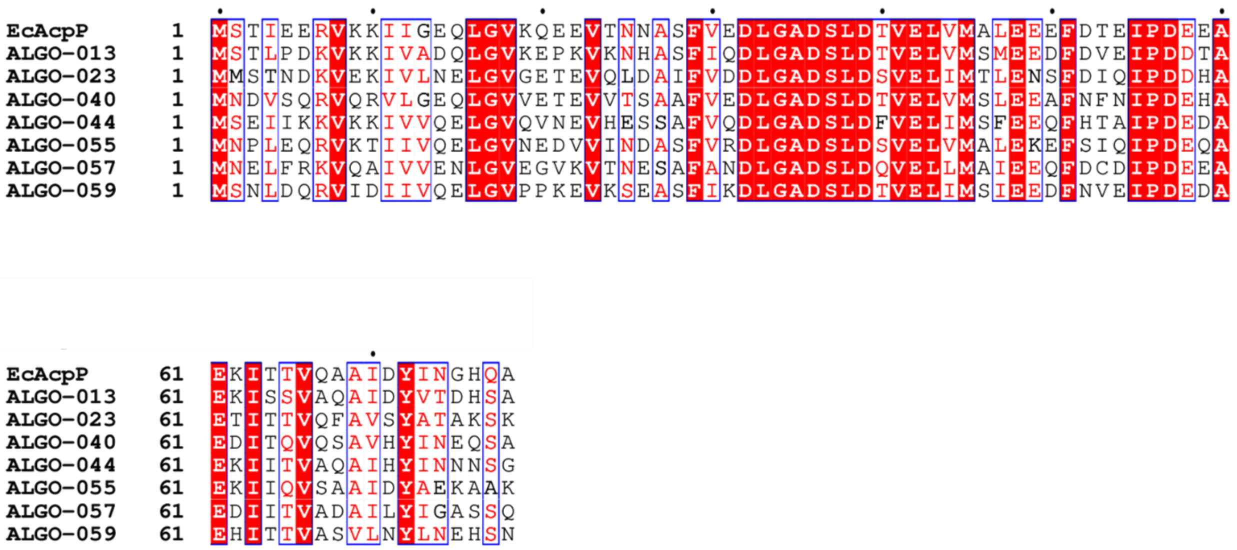


**Fig. S2** Multiple sequence alignment of ALGO-CP candidates and *Ec*AcpP.

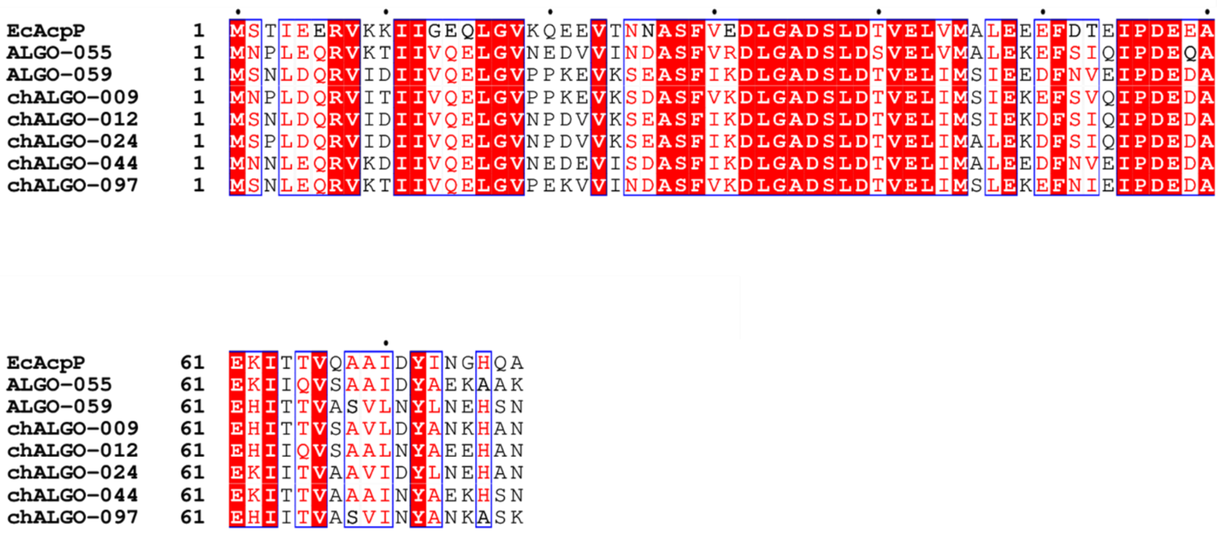


**Fig. S3** Multiple sequence alignment of chALGO-CP chimeras, parental ALGO-055 and ALGO-059, and *Ec*AcpP.

#### Purification Data (Figs. S4-S14)

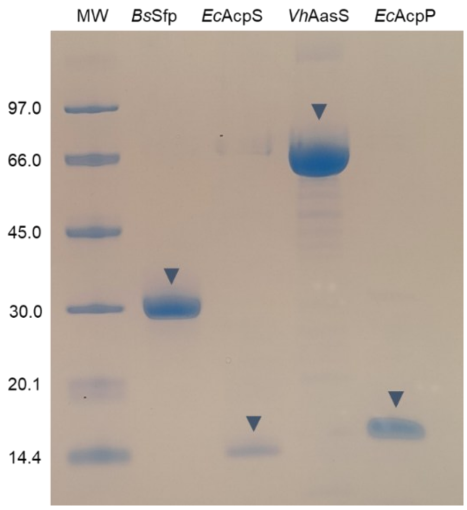


**Fig. S4** Annotated SDS-PAGE gel photograph showing purified *Bs*Sfp, *Ec*AcpS, *Vh*AasS and *Ec*AcpP. An Amersham low molecular weight calibration kit was used as the protein ladder (lane MW).

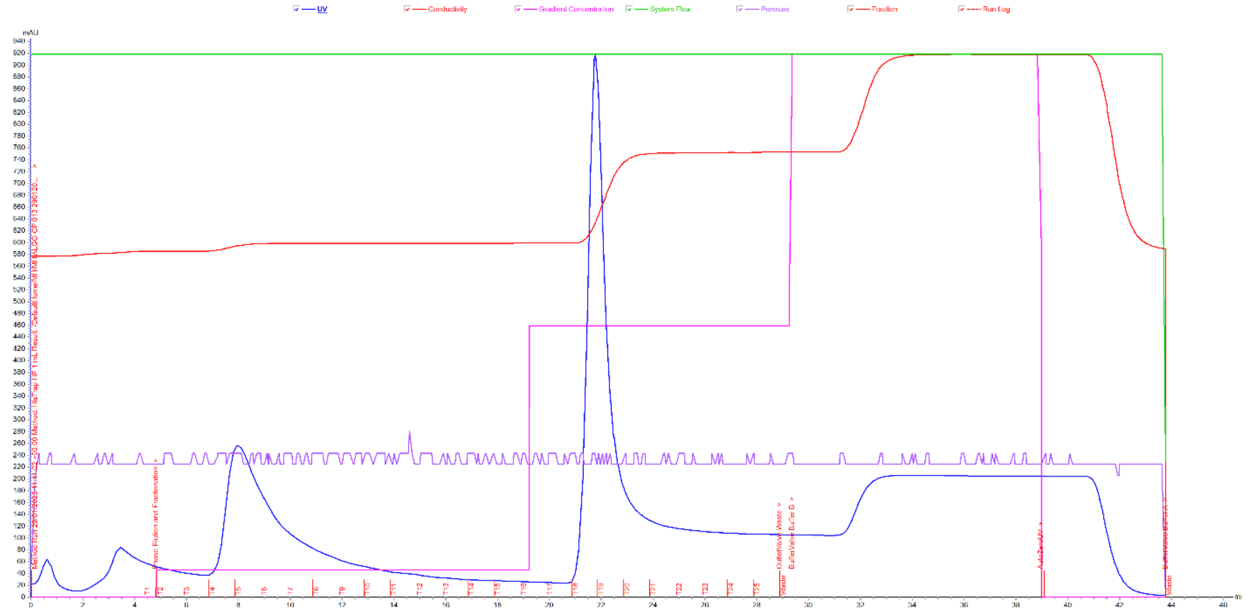


**Fig. S5** $A_{280}$ chromatogram of ALGO-013 purification by $Ni^{2+}$-affinity chromatography.

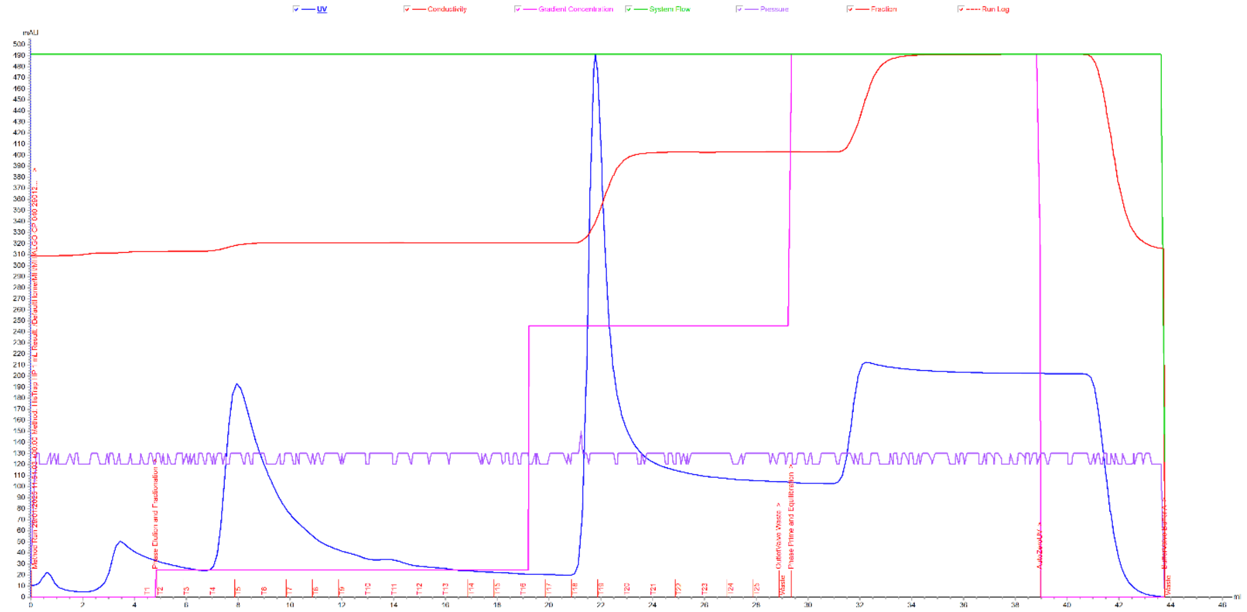


**Fig. S6** $A_{280}$ chromatogram of ALGO-040 purification by $Ni^{2+}$-affinity chromatography.

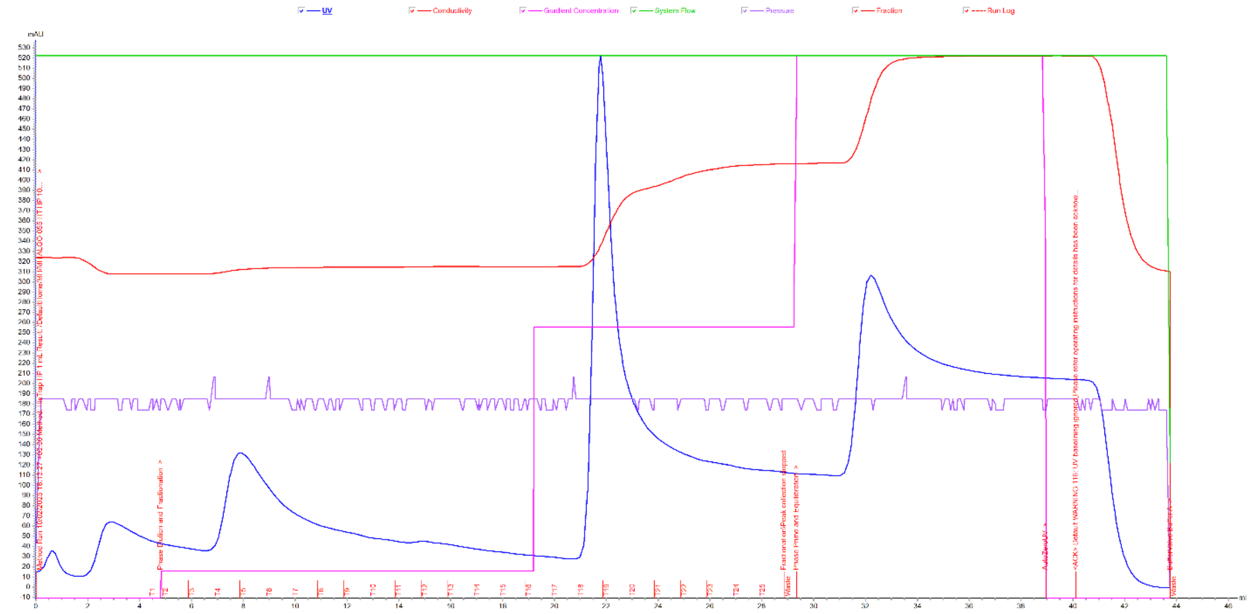


**Fig. S7** $A_{280}$ chromatogram of ALGO-055 purification by $Ni^{2+}$-affinity chromatography.

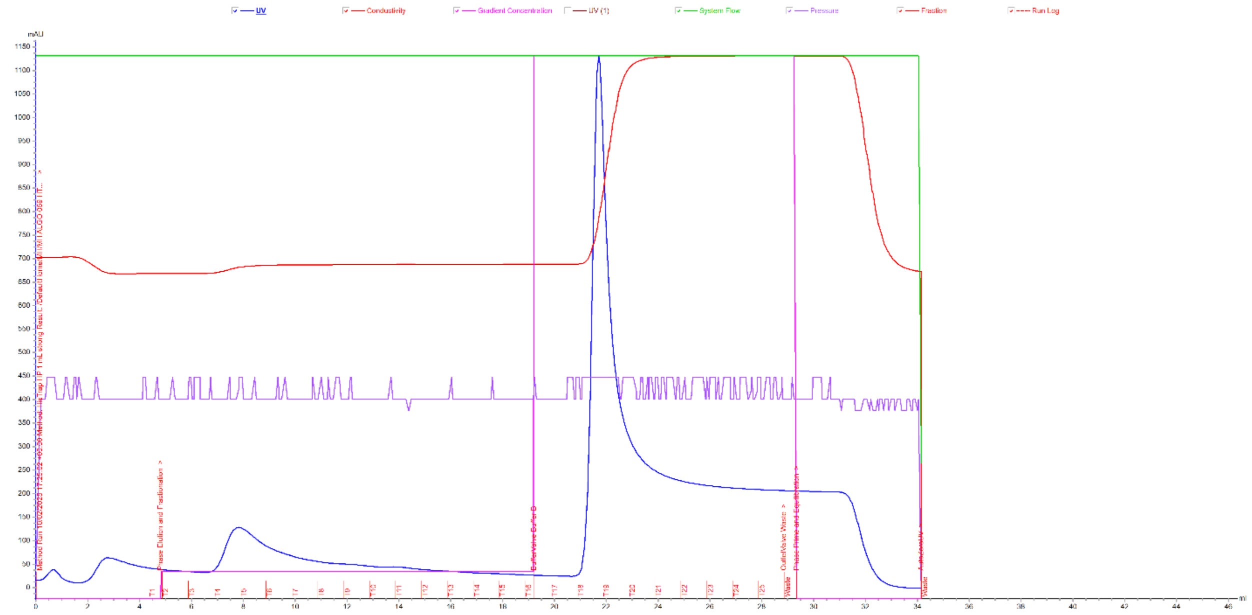


**Fig. S8** $A_{280}$ chromatogram of ALGO-059 purification by $Ni^{2+}$-affinity chromatography.

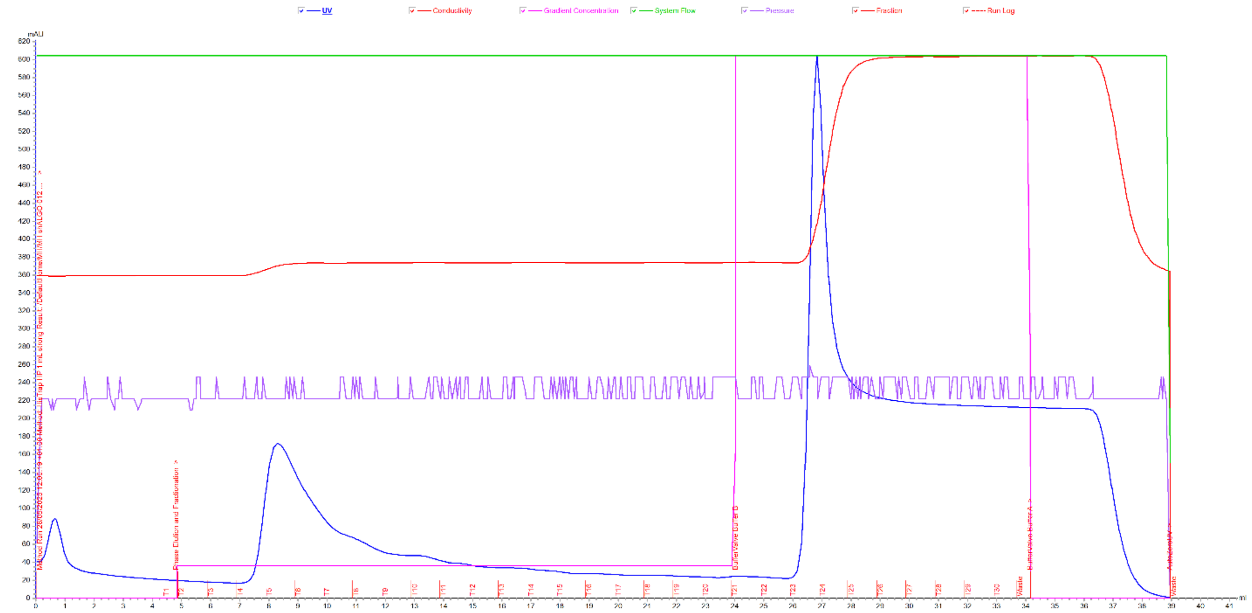


**Fig. S9** $A_{280}$ chromatogram of $^{ch}$ALGO-012 purification by $Ni^{2+}$-affinity chromatography.

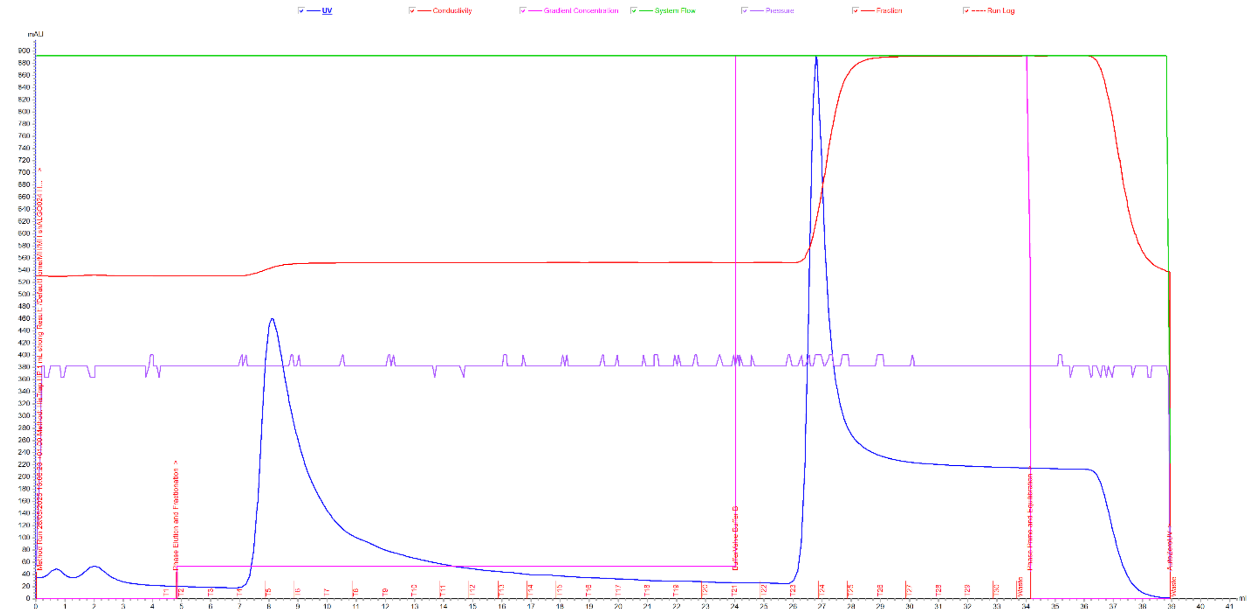


**Fig. S10** $A_{280}$ chromatogram of $^{ch}$ALGO-024 purification by $Ni^{2+}$-affinity chromatography.

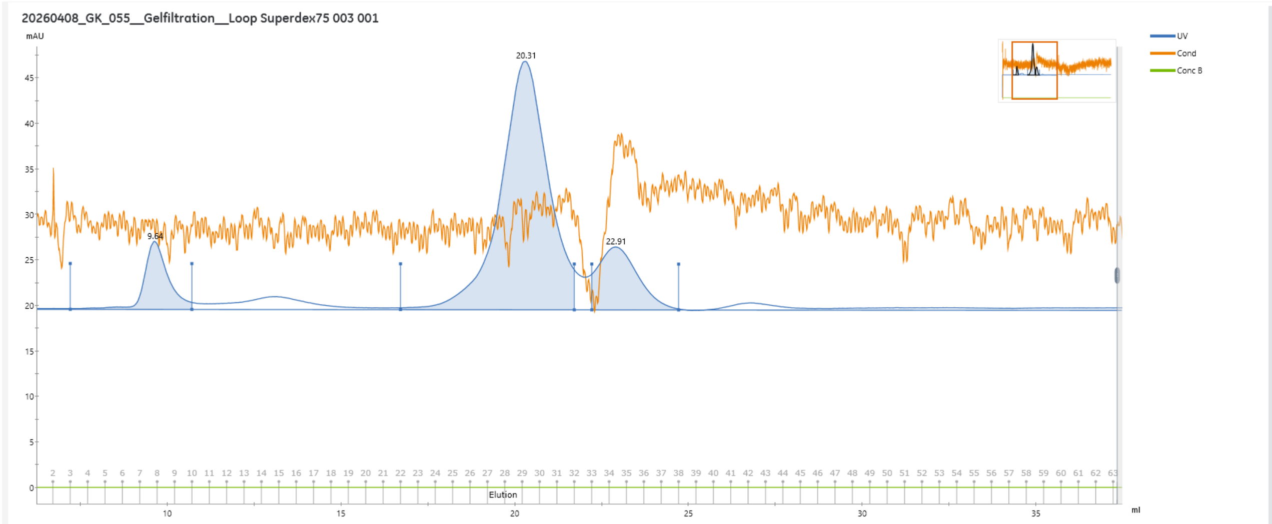


Peak Table - UV

| Peak | Retention ml | Area ml*mAU | Area % | Ext coeff. mg ml⁻¹ cm⁻¹ | Fraction(s) | Volume ml | Conductivity mS/cm |
|---|---|---|---|---|---|---|---|
| Peak A | 9.635 | 5.942 | 10.2 | | 3 - 9 | 3.500 | 5.28 |
| Peak B | 20.310 | 43.23 | 74.21 | | 22 - 31 | 5.000 | 5.28 |
| Peak C | 22.907 | 9.086 | 15.6 | | 33 - 37 | 2.499 | 5.28 |

**Fig. S11** $A_{280}$ chromatogram of purified $C_{12}$-*Ec*AcpP using Superdex S75 SEC (20.31 mL).

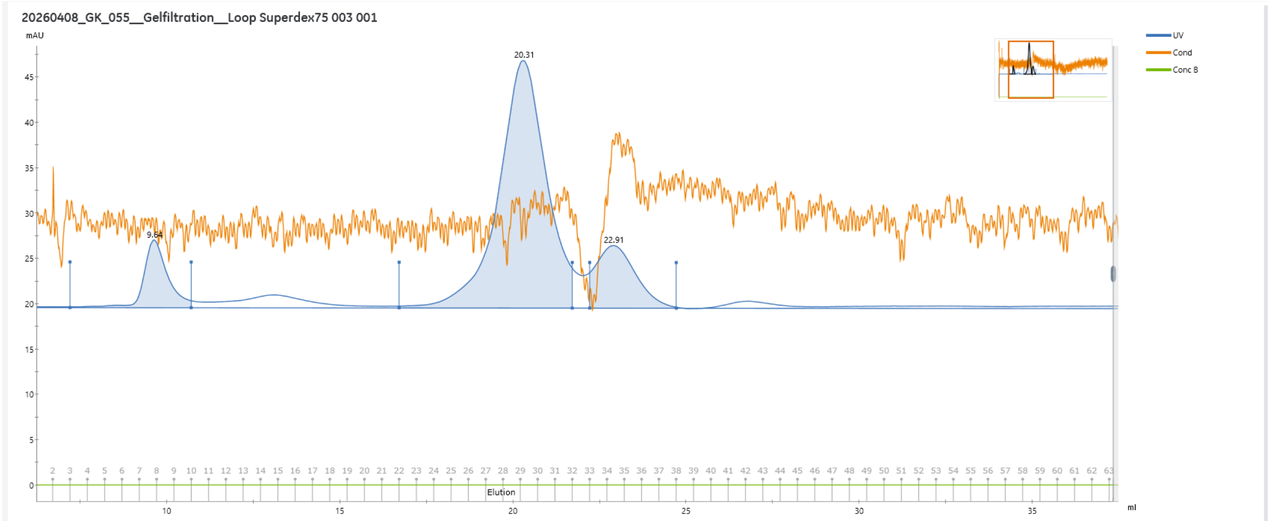


Peak Table - UV

| Peak | Retention ml | Area ml*mAU | Area % | Ext coeff. mg ml⁻¹ cm⁻¹ | Fraction(s) | Volume ml | Conductivity mS/cm |
|---|---|---|---|---|---|---|---|
| Peak A | 9.635 | 5.942 | 10.2 | | 3 - 9 | 3.500 | 5.28 |
| Peak B | 20.310 | 43.23 | 74.21 | | 22 - 31 | 5.000 | 5.28 |
| Peak C | 22.907 | 9.086 | 15.6 | | 33 - 37 | 2.499 | 5.28 |

**Fig. S12** $A_{280}$ chromatogram of purified $C_{12}$-ALGO-055 using Superdex S75 SEC (20.31 mL).

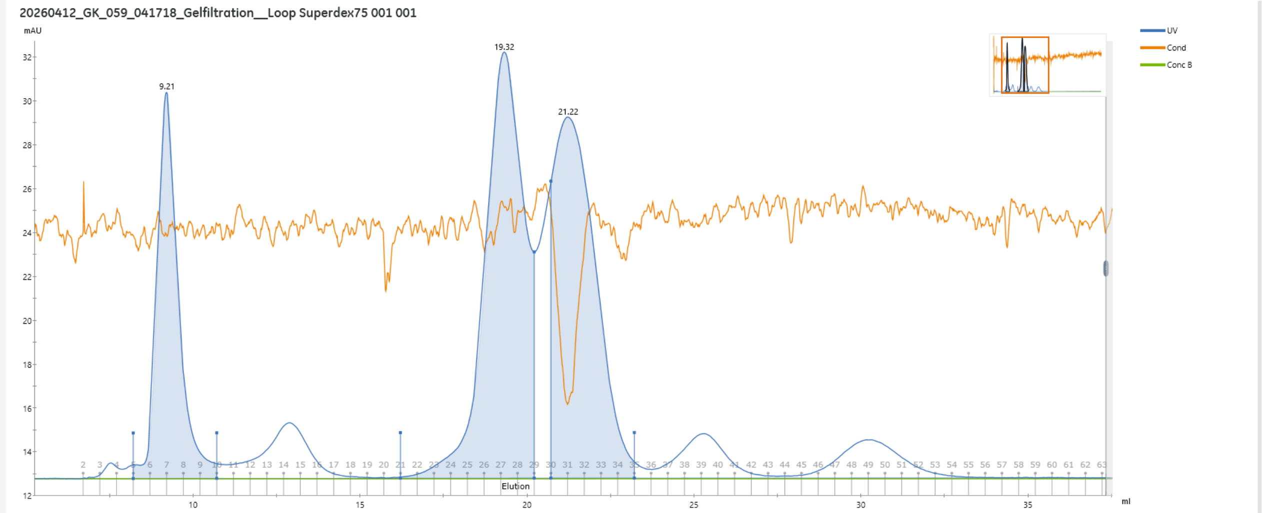


Peak Table - UV

| Peak | Retention (ml) | Area (ml*mAU) | Area % | Ext coeff. (mg ml⁻¹ cm⁻¹) | Fraction(s) | Volume (ml) | Conductivity (mS/cm) |
|---|---|---|---|---|---|---|---|
| Peak A | 9.207 | 12.85 | 20.46 | | 5 - 9 | 2.501 | 5.28 |
| Peak B | 19.319 | 26.13 | 41.58 | | 21 - 28 | 4.004 | 5.28 |
| Peak C | 21.221 | 23.85 | 37.96 | | 30 - 34 | 2.499 | 5.27 |

**Fig. S13** $A_{280}$ chromatogram of purified $C_{12}$-ALGO-059 using Superdex S75 SEC (20.31 mL).

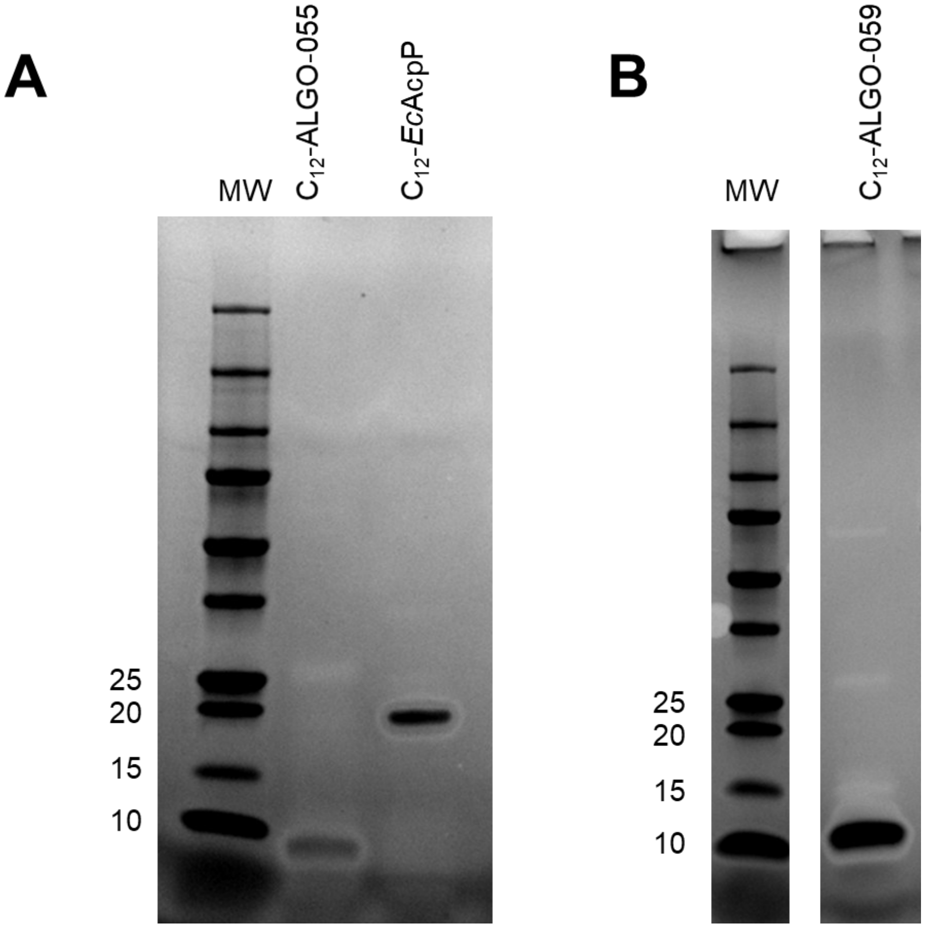


**Fig. S14** SDS-PAGE of purified acylated proteins (A) $C_{12}$-*Ec*AcpP and $C_{12}$-ALGO-055 (B) $C_{12}$-ALGO-059. Gels were imaged using a FluorChem M FM1059.

168

#### LC/ESI-MS data (Figs. S11-S31)

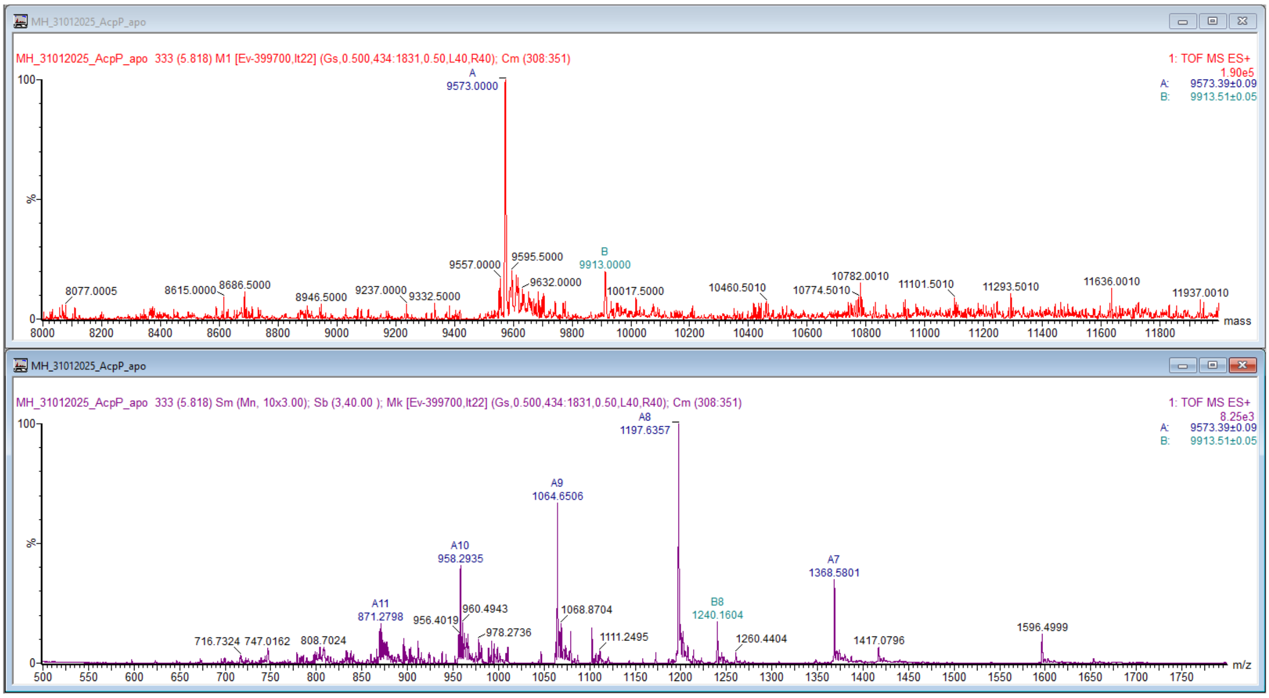


**Fig. S15** *apo-Ec*AcpP charge envelope (purple) and deconvoluted mass (red).

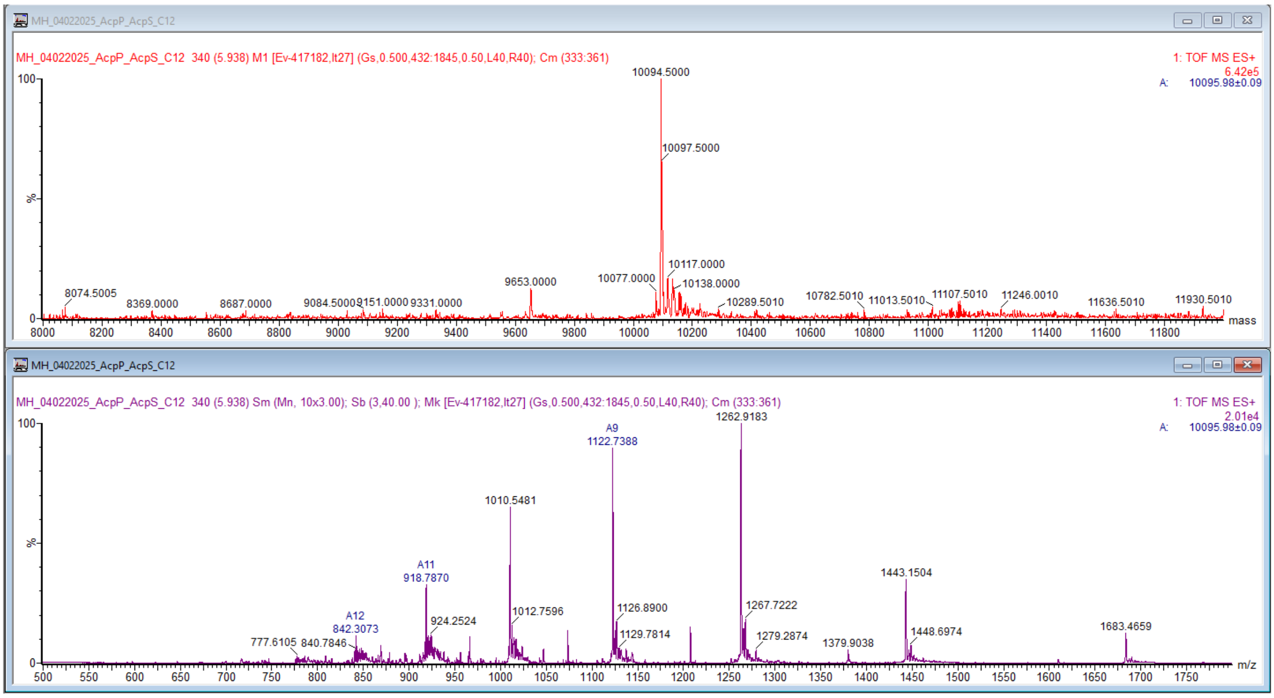


**Fig. S16** *holo-Ec*AcpP charge envelope (purple) and deconvoluted mass (red).

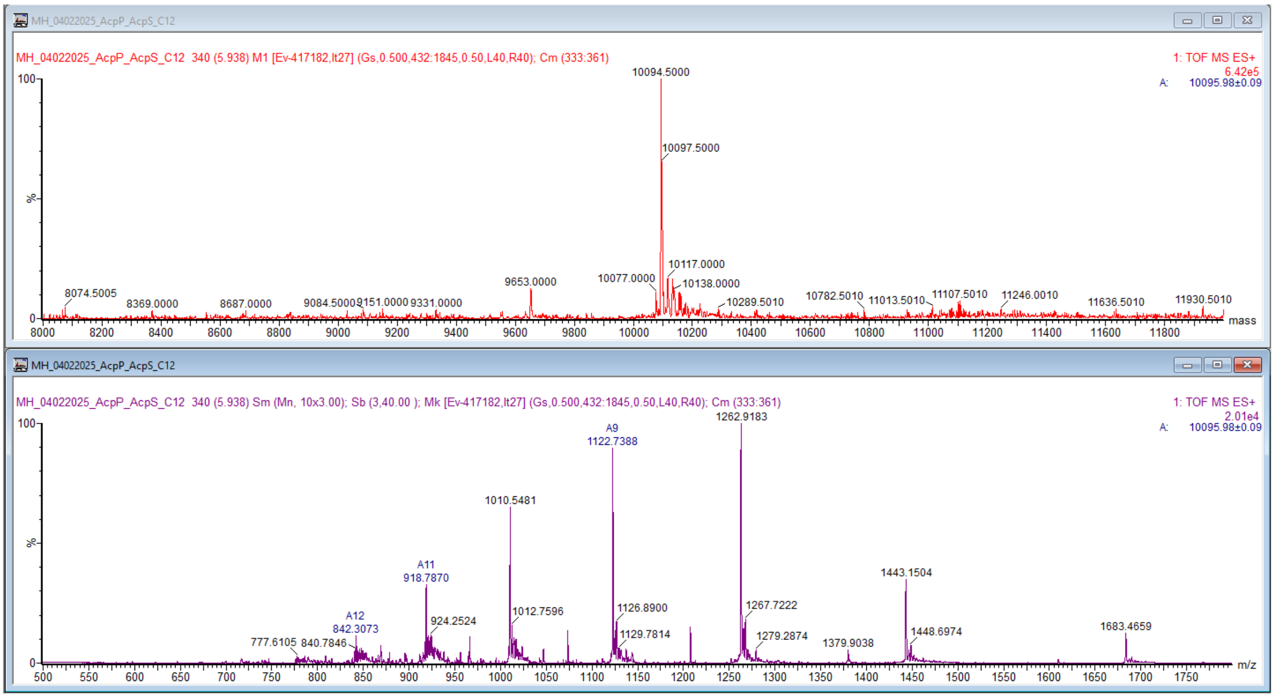


**Fig. S17** C12-*Ec*AcpP charge envelope (purple) and deconvoluted mass (red).

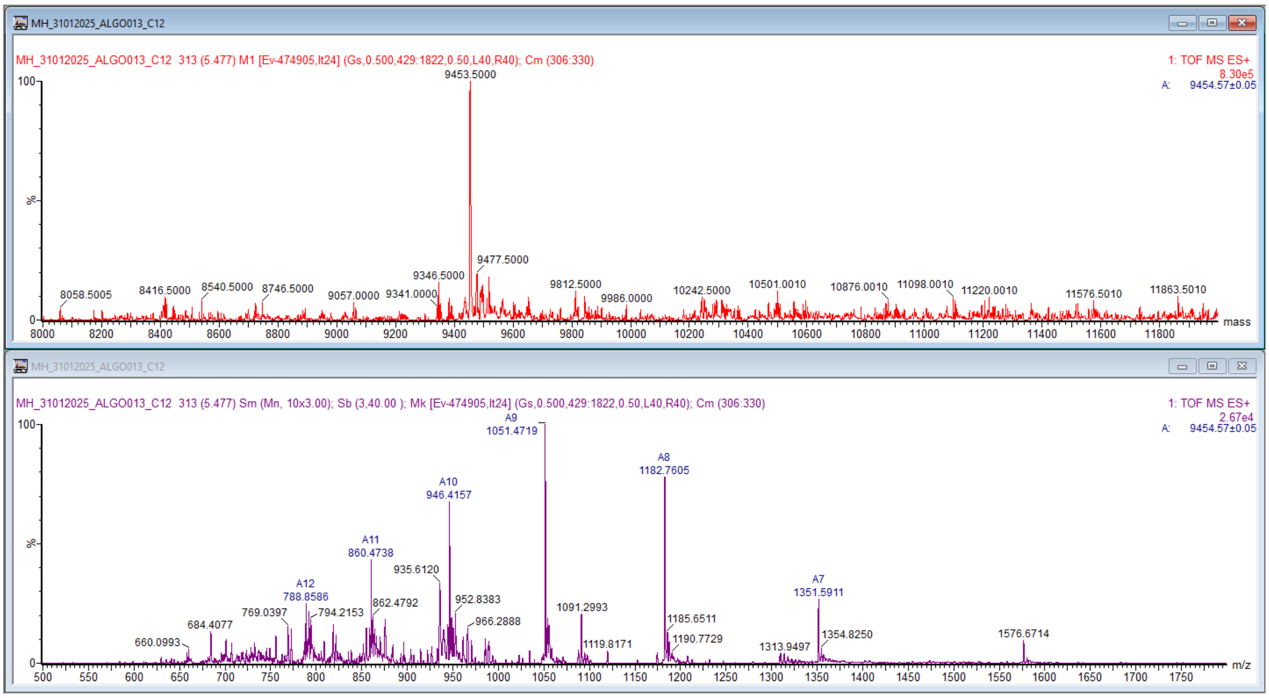


**Fig. S18** *apo*-ALGO-013 charge envelope (purple) and deconvoluted mass (red).

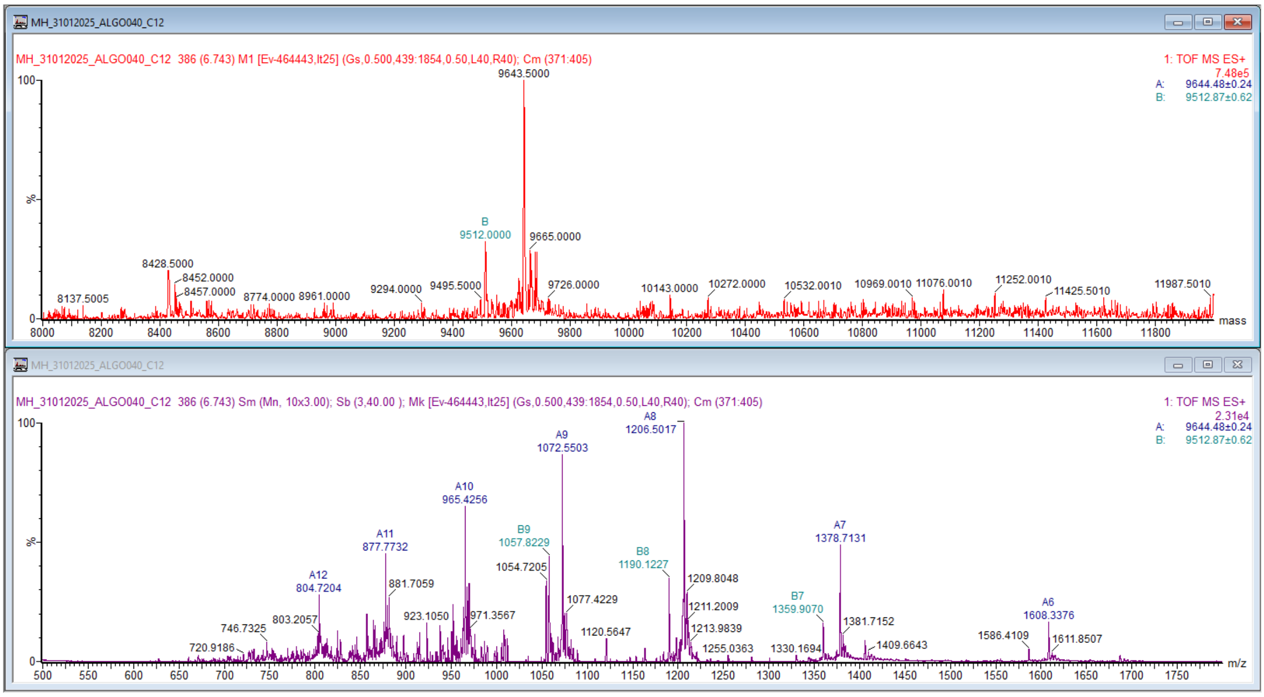


**Fig. S19** *apo*-ALGO-040 charge envelope (purple) and deconvoluted mass (red).

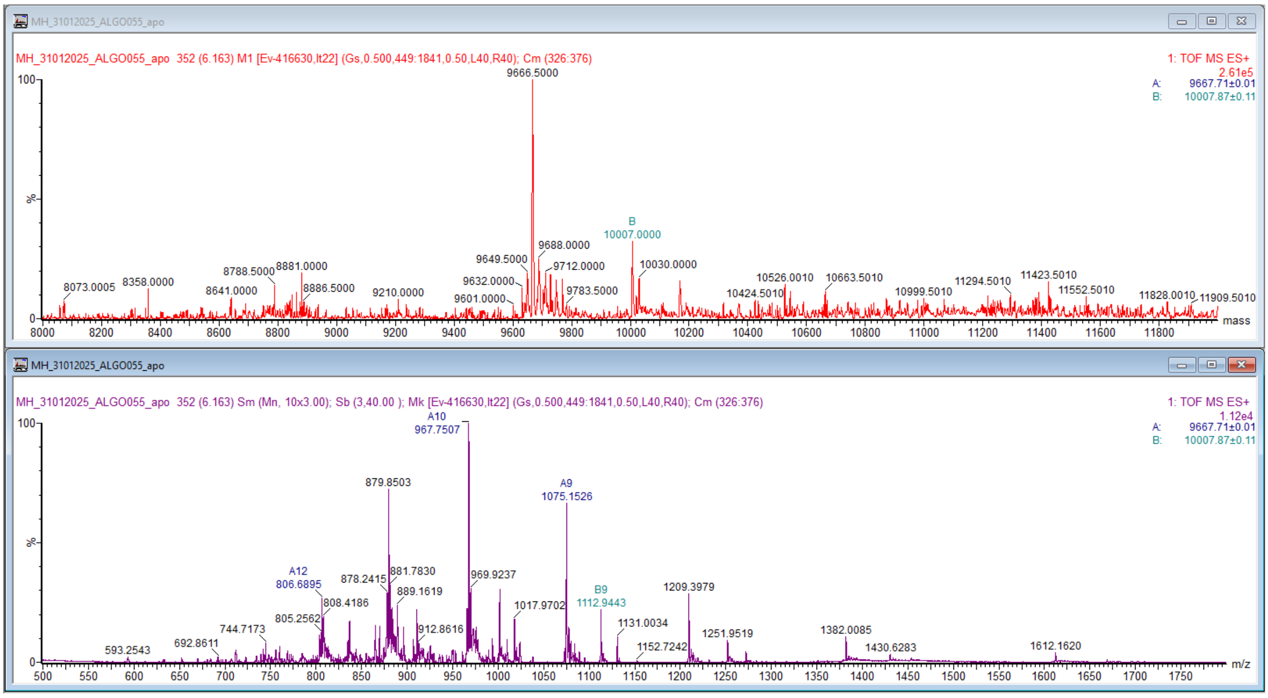


**Fig. S20** *apo*-ALGO-055 charge envelope (purple) and deconvoluted mass (red).

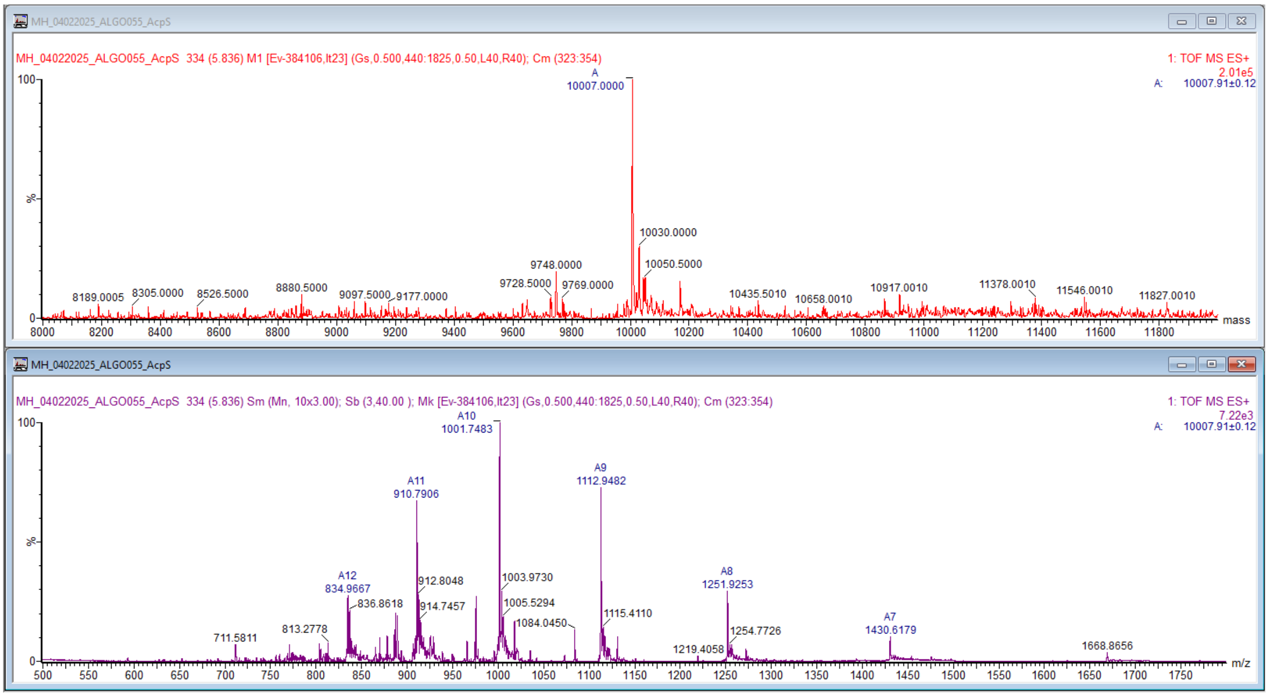


**Fig. S21** *holo*-ALGO-055 charge envelope (purple) and deconvoluted mass (red).

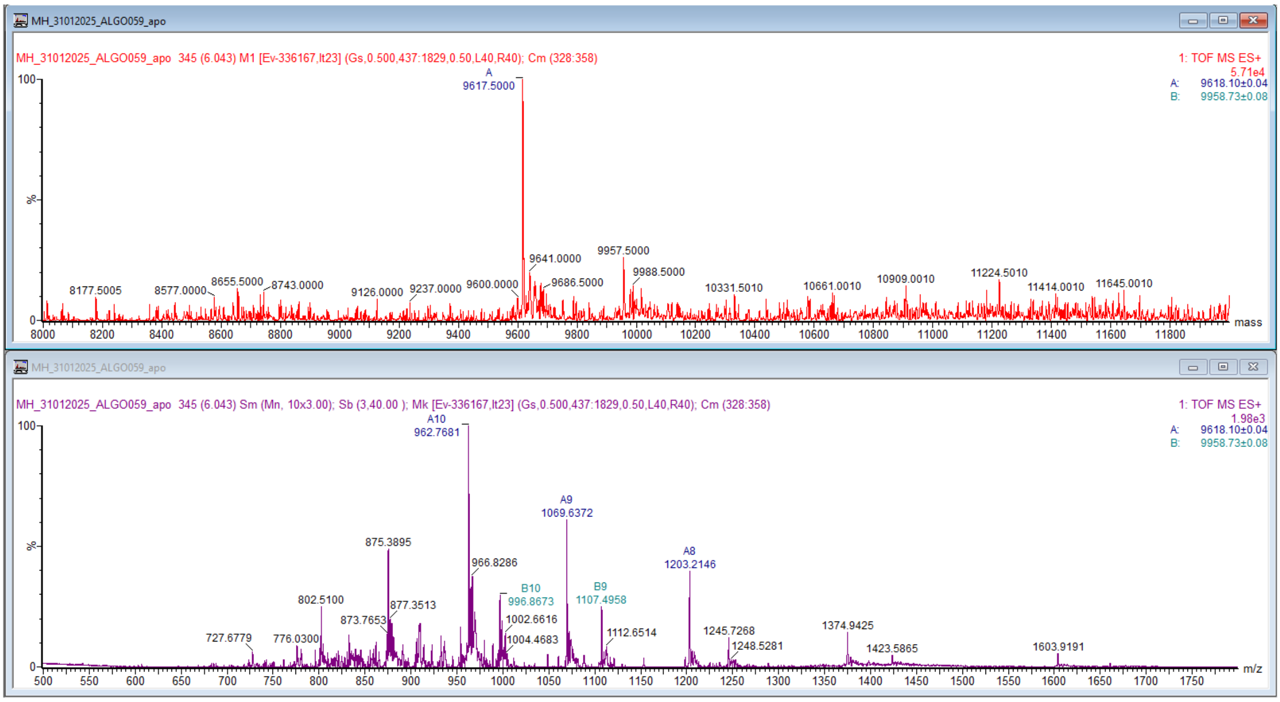


**Fig. S22** *apo*-ALGO-059 charge envelope (purple) and deconvoluted mass (red).

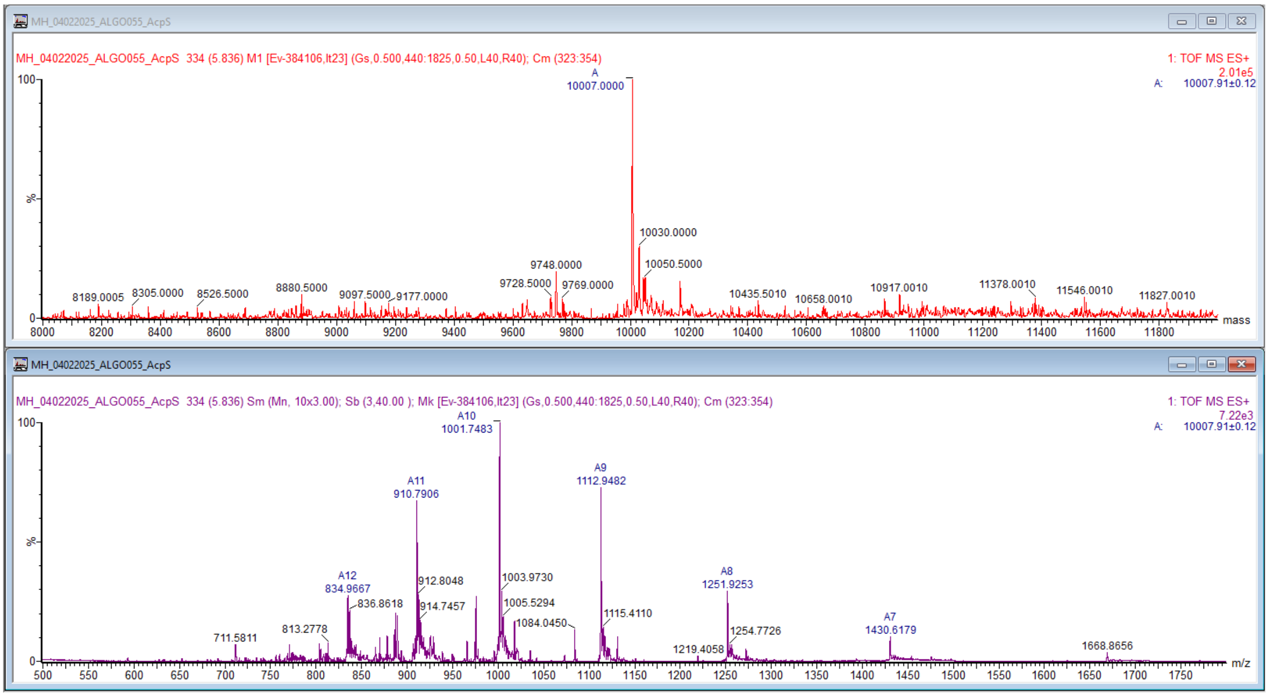


**Fig. S23** *holo*-ALGO-059 charge envelope (purple) and deconvoluted mass (red).

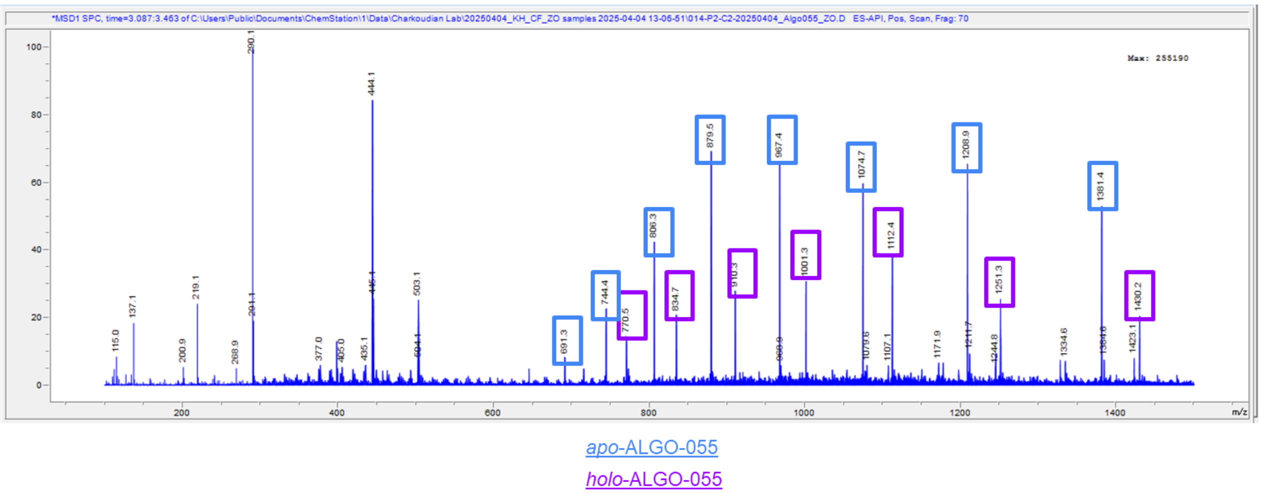


**Fig. S24** Attempted apo→holo conversion of immobilised ALGO-055 using *Ec*AcpS. Recorded using an Agilent Technologies InfinityLab G6125B single quadrupole MS.

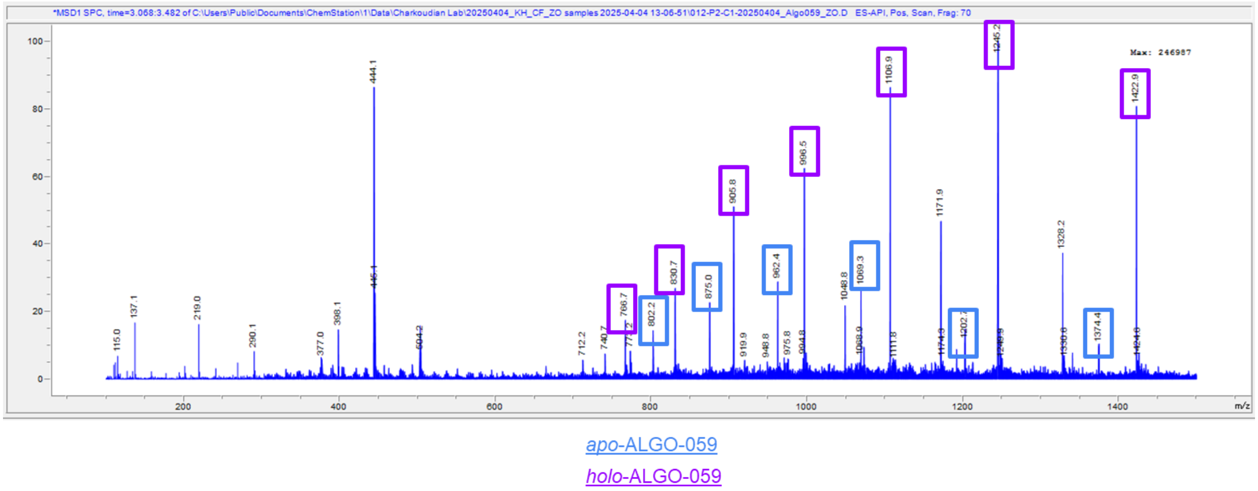


**Fig. S25** Attempted apo→holo conversion of immobilised ALGO-059 using *Ec*AcpS. Recorded using an Agilent Technologies InfinityLab G6125B single quadrupole MS.

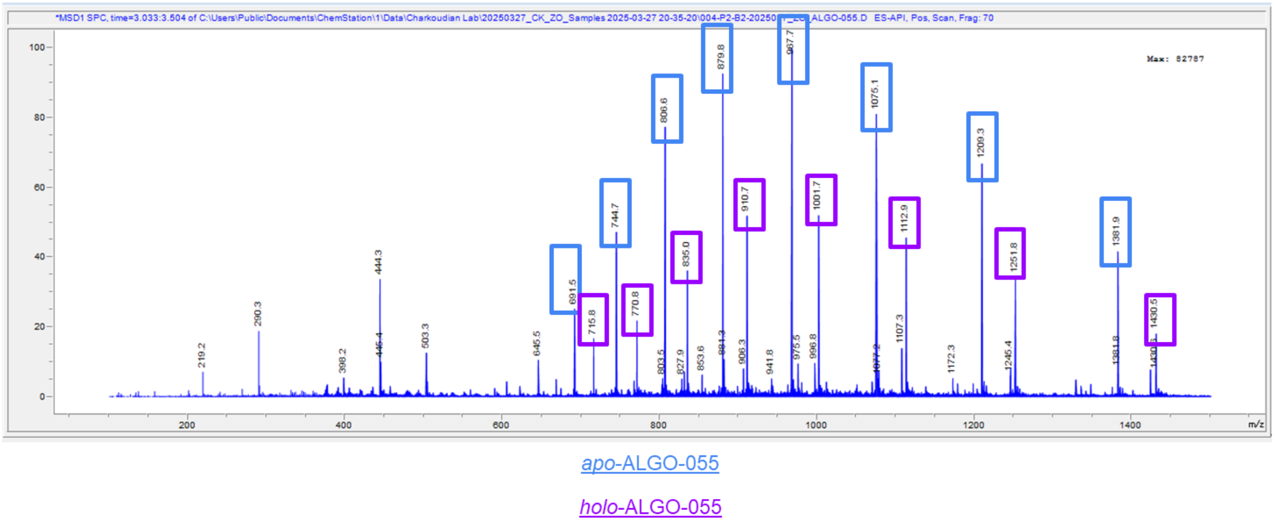


**Fig. S26** Attempted apo→holo conversion of ALGO-055 using *Bs*Sfp. Recorded using an Agilent Technologies InfinityLab G6125B single quadrupole MS.

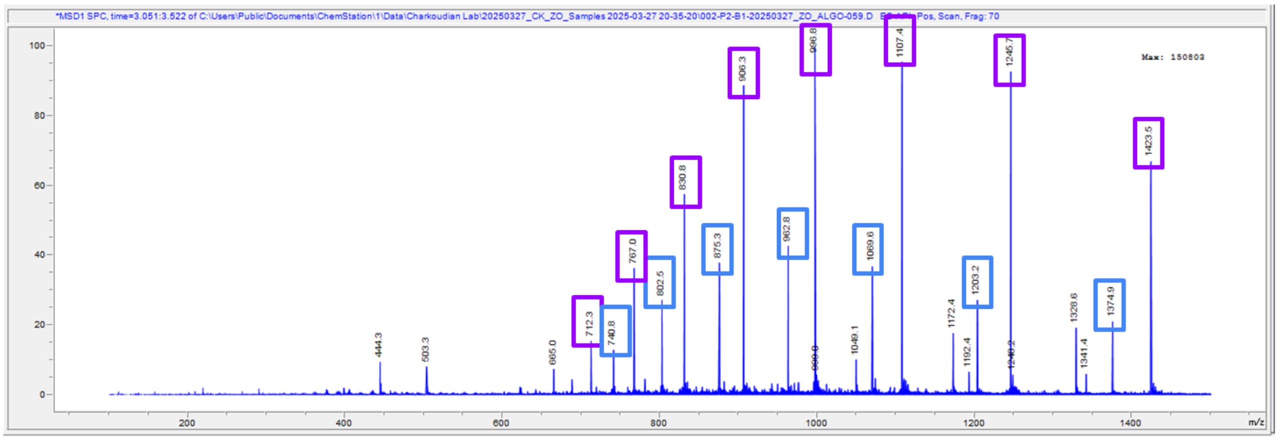


apo-ALGO-059

holo-ALGO-059

**Fig. S27** Attempted apo→holo conversion of ALGO-059 using *Bs*Sfp. Recorded using an Agilent Technologies InfinityLab G6125B single quadrupole MS.

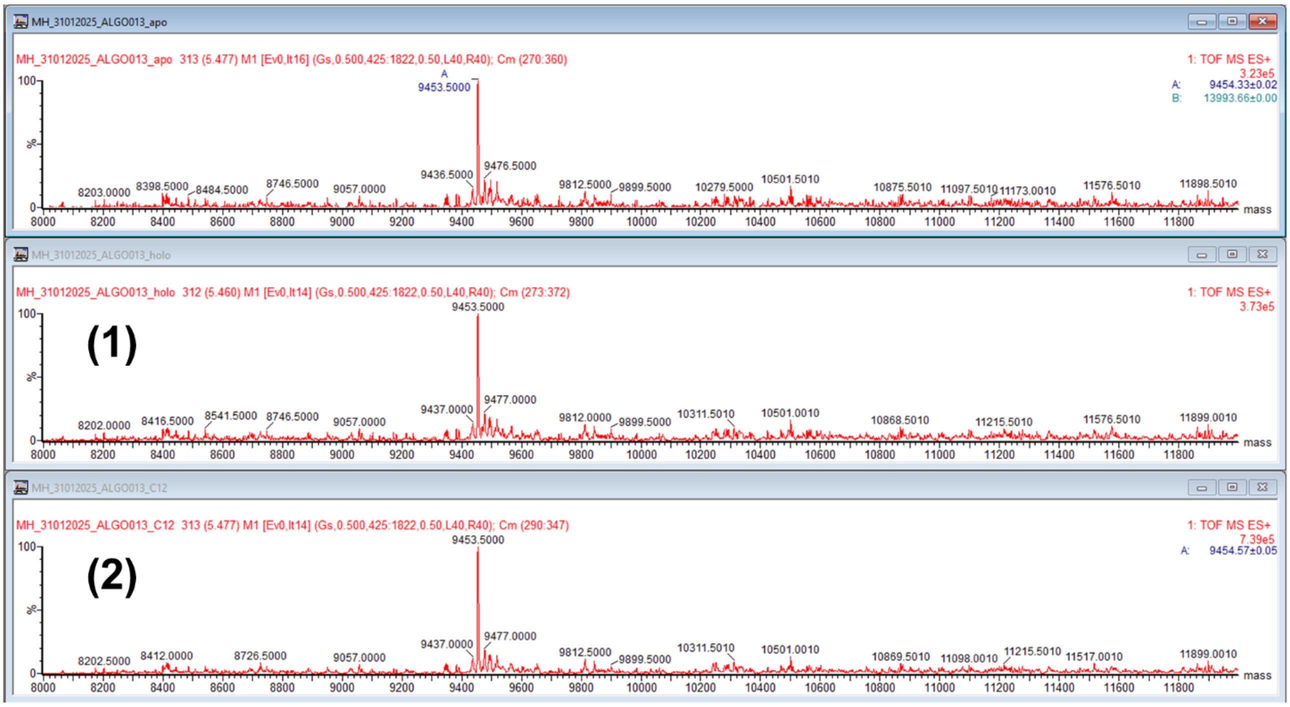


**Fig. S28** Unsuccessful apo→holo→acyl conversion of ALGO-013 using *Ec*AcpS (1) and VhAasS (2).

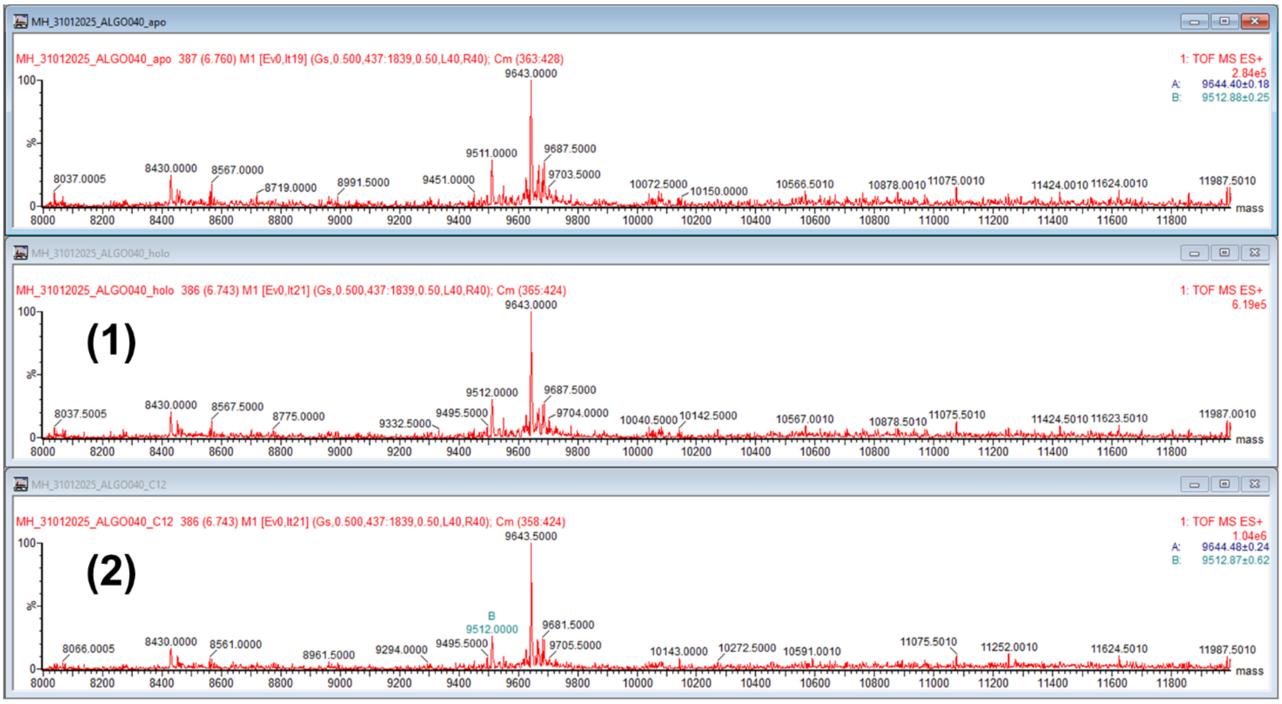


**Fig. S29** Unsuccessful apo→holo→acyl conversion of ALGO-040 using *Ec*AcpS (1) and VhAasS (2).

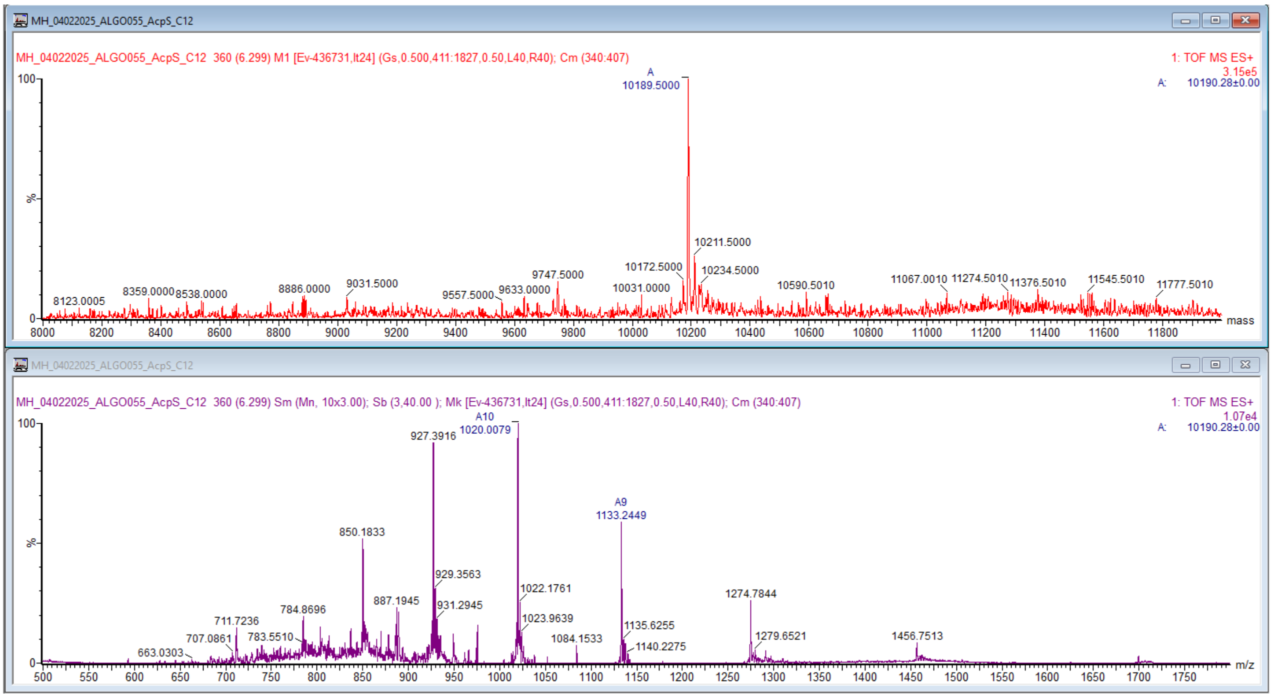


**Fig. S30** $C_{12}$-ALGO-055 charge envelope.

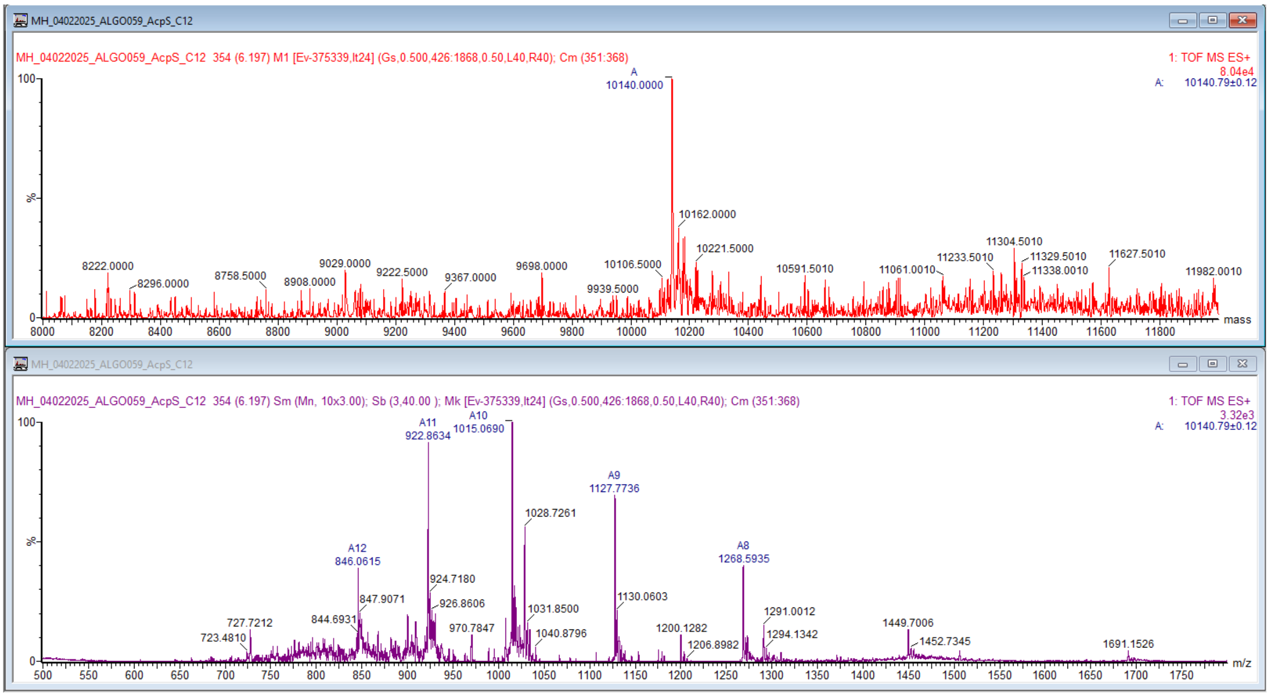


**Fig. S31** C12-ALGO-059 charge envelope.

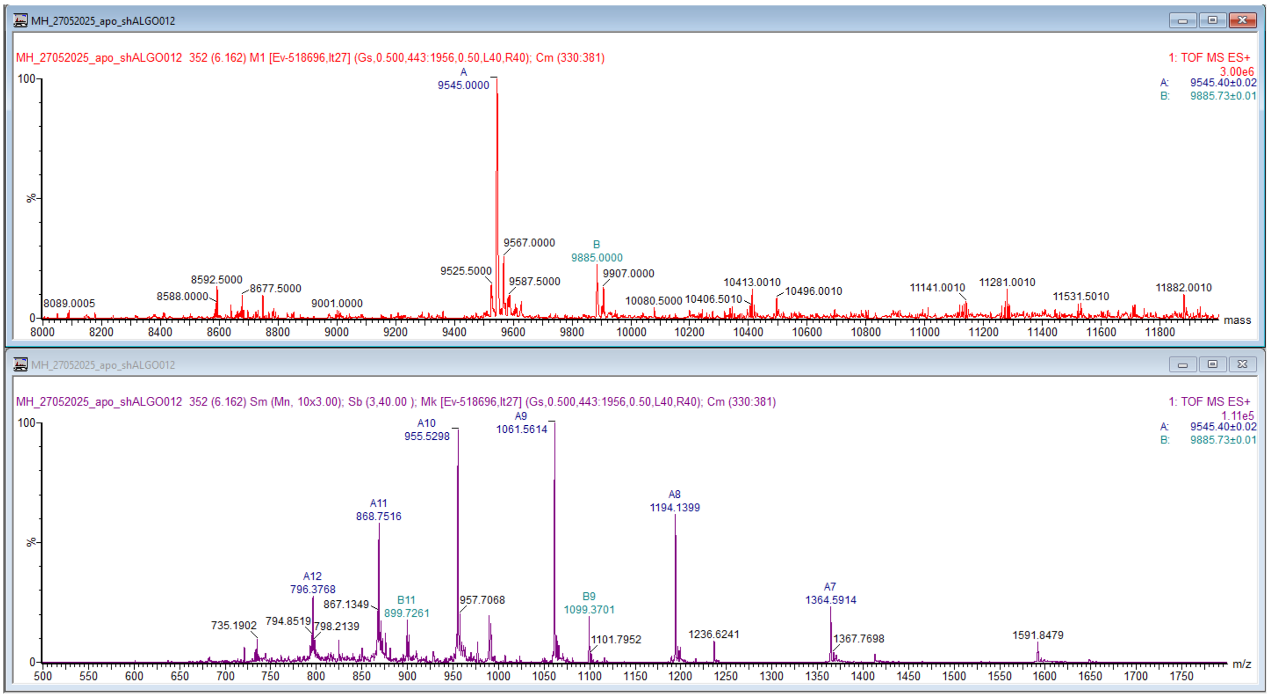


**Fig. S32** *apo*-[ch]ALGO-012 charge envelope.

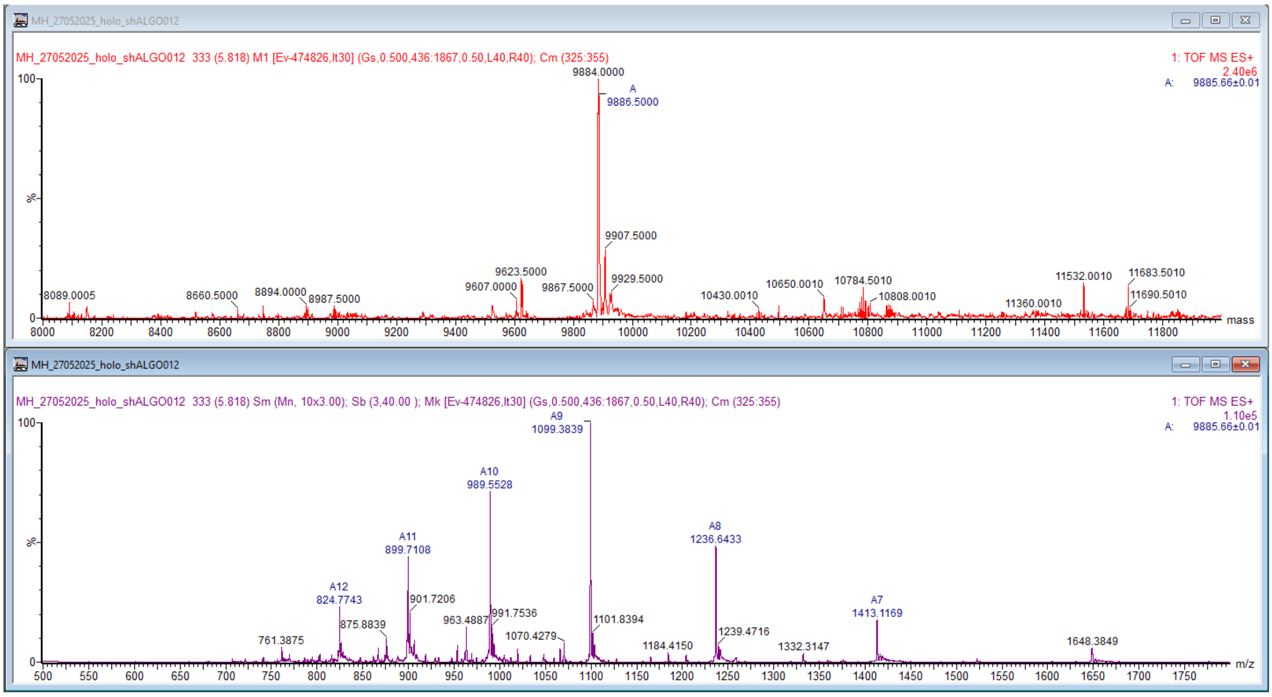


**Fig. S33** *holo*-$^{ch}$ALGO-012 charge envelope.

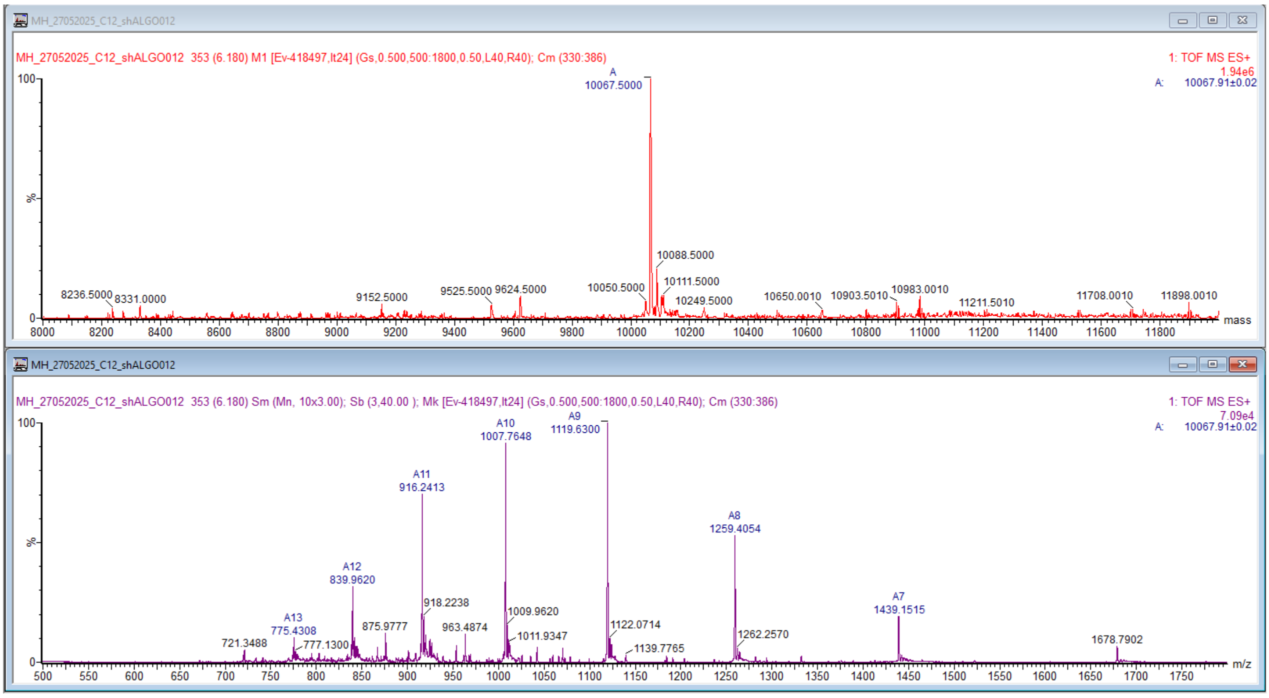


**Fig. S34** $C_{12}$-$^{ch}$ALGO-012 charge envelope.

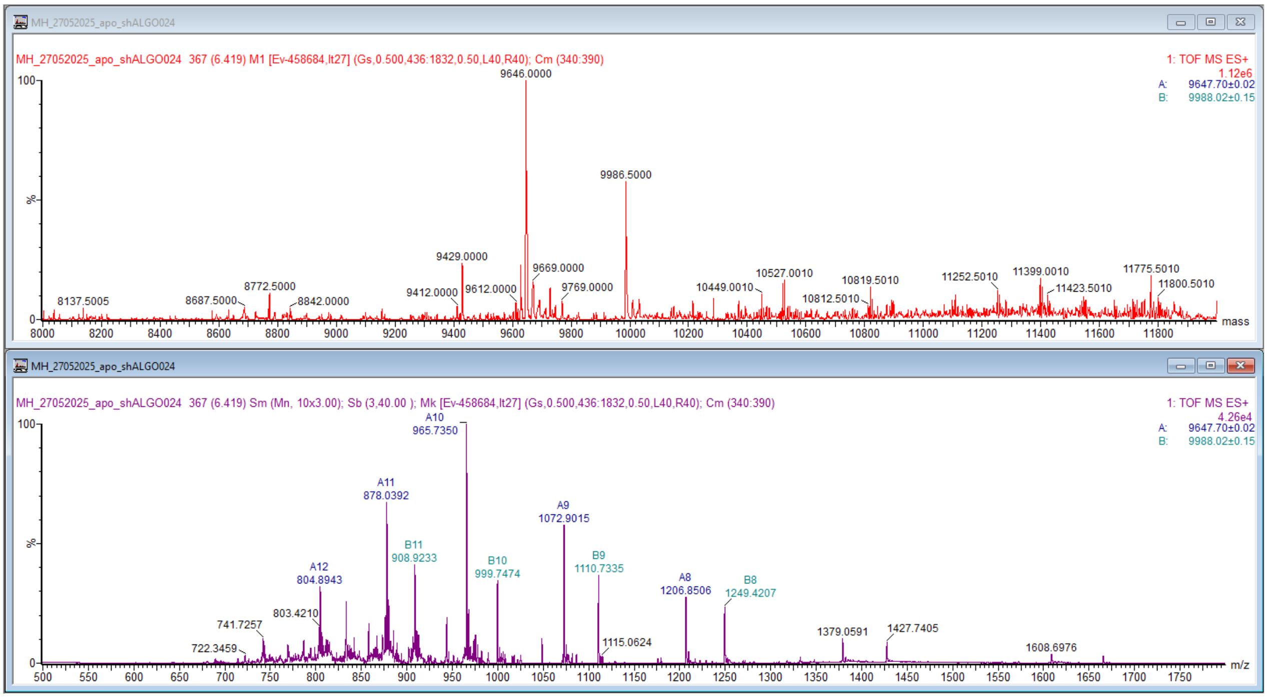


**Fig. S35** *apo*-$^{ch}$ALGO-024 charge envelope.

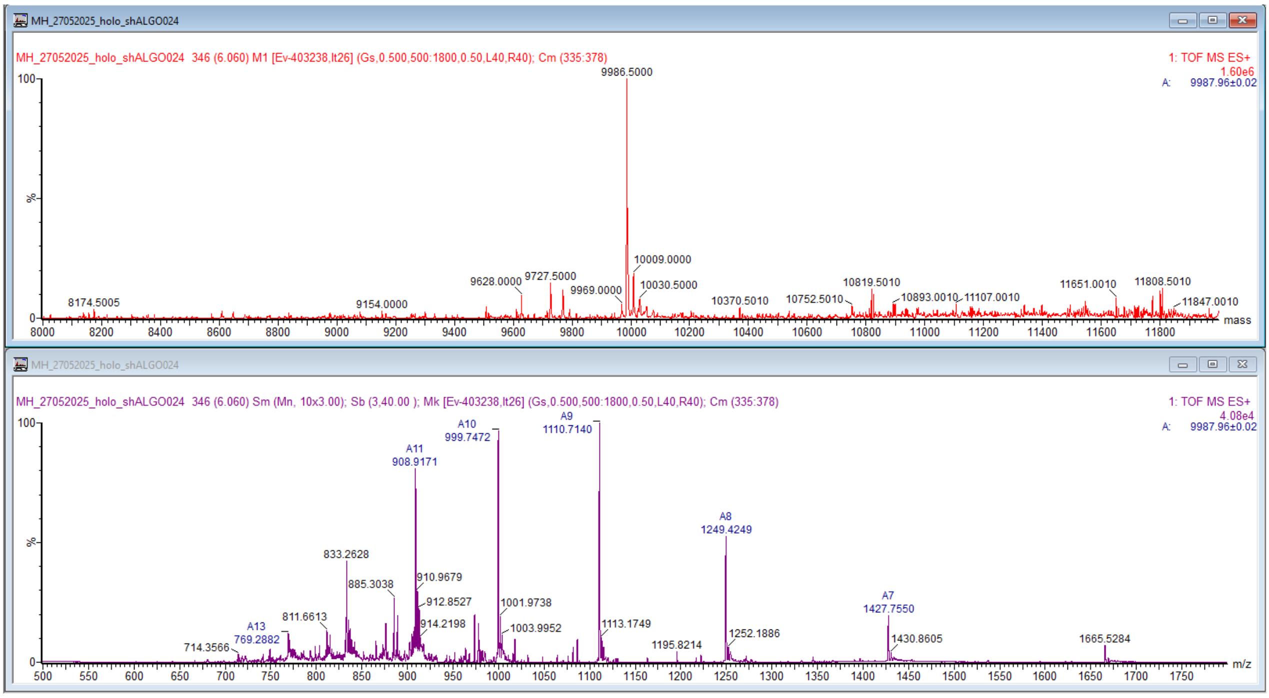


**Fig. S36** *holo*-*Ec*AcpP charge envelope.

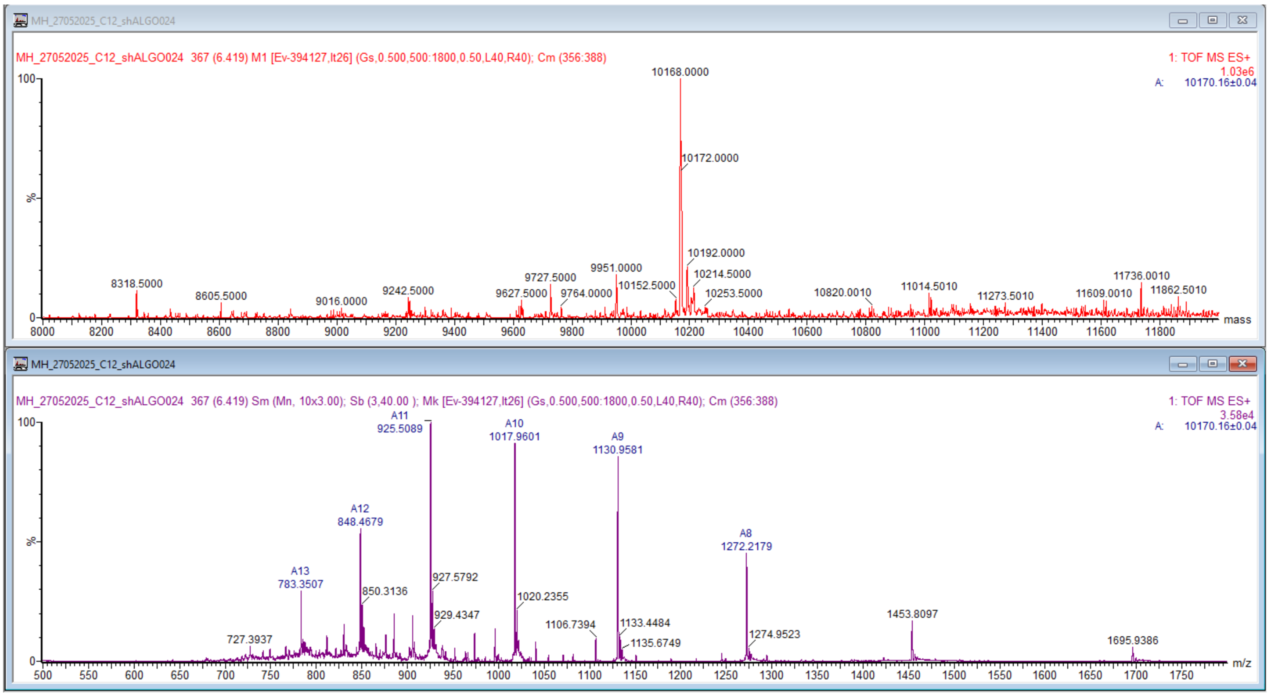


**Fig. S37** $C_{12}$-$^{ch}$ALGO-024 charge envelope.

#### Structural modelling and MDS (Figs. S32-S41)

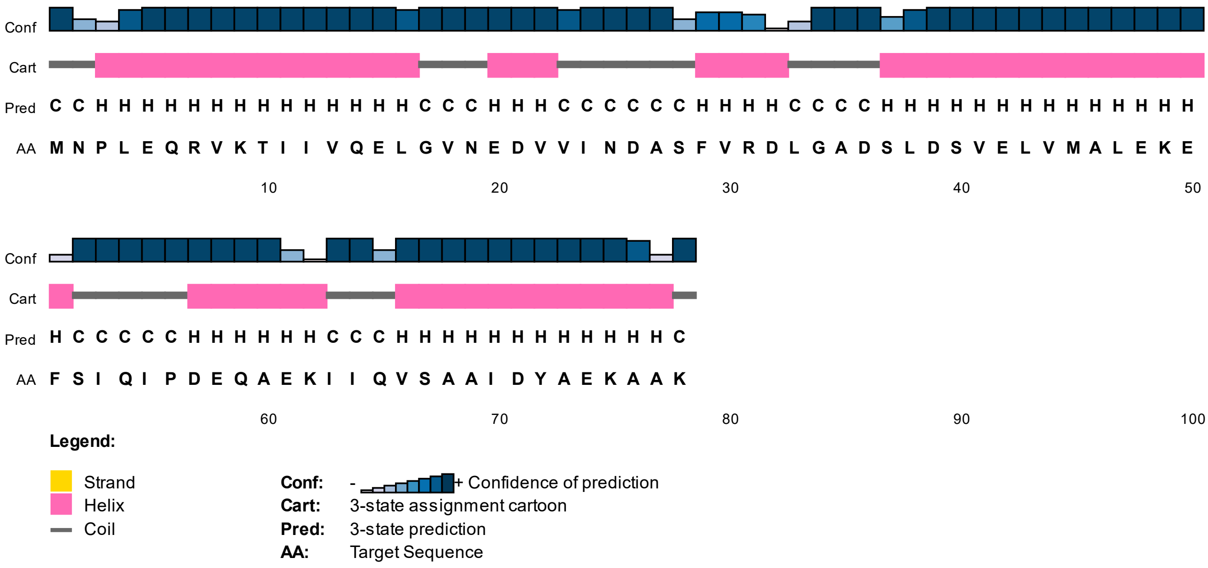


**Fig. S38** PSIPRED secondary structure prediction from ALGO-055 primary sequence.

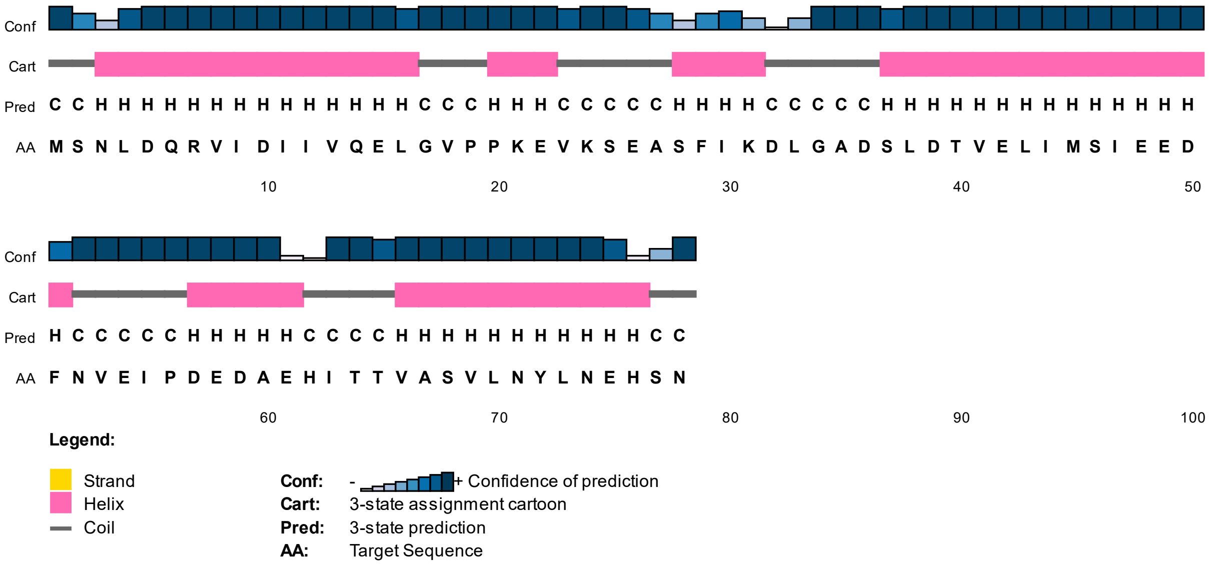


**Fig. S39** PSIPRED secondary structure prediction from ALGO-059 primary sequence.

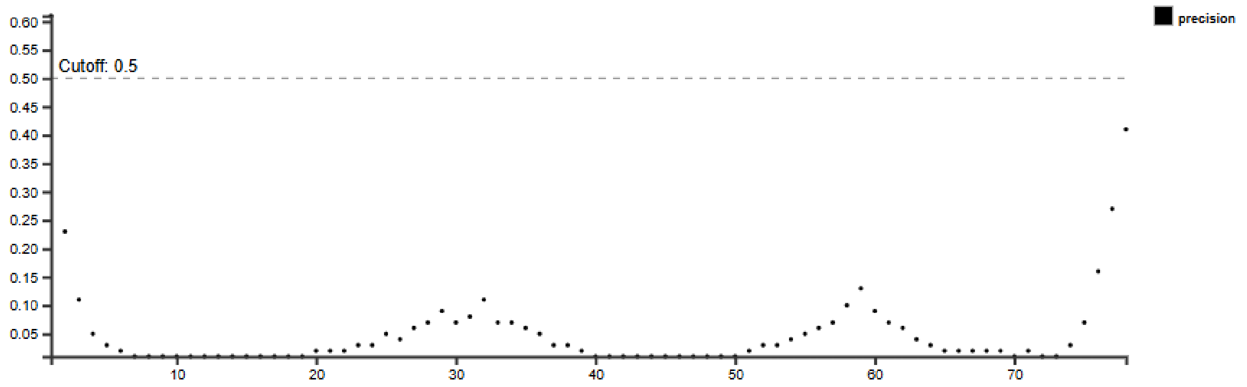


**Fig. S40** DISOPRED3 disorder prediction from ALGO-059 primary sequence. Cut-off for protein disorder classification = 0.5.

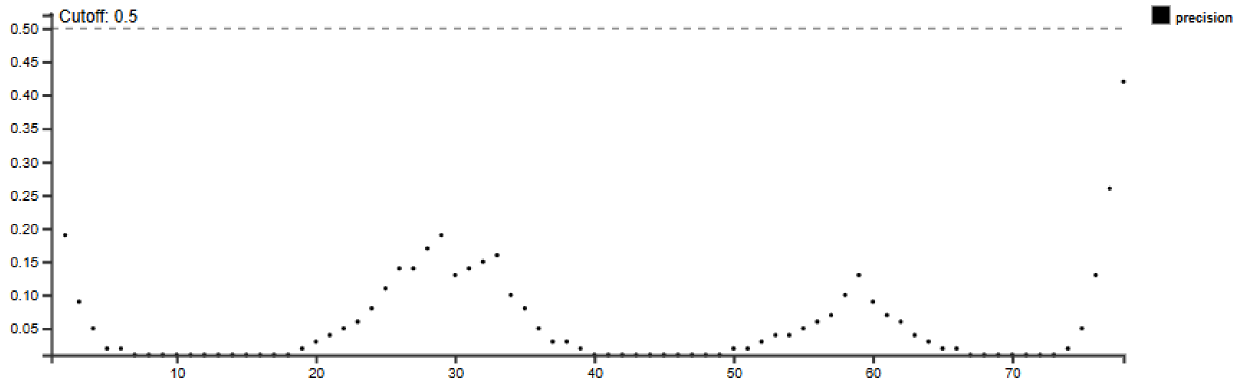


**Fig. S41** DISOPRED3 disorder prediction from ALGO-059 primary sequence. Cut-off for protein disorder classification = 0.5.

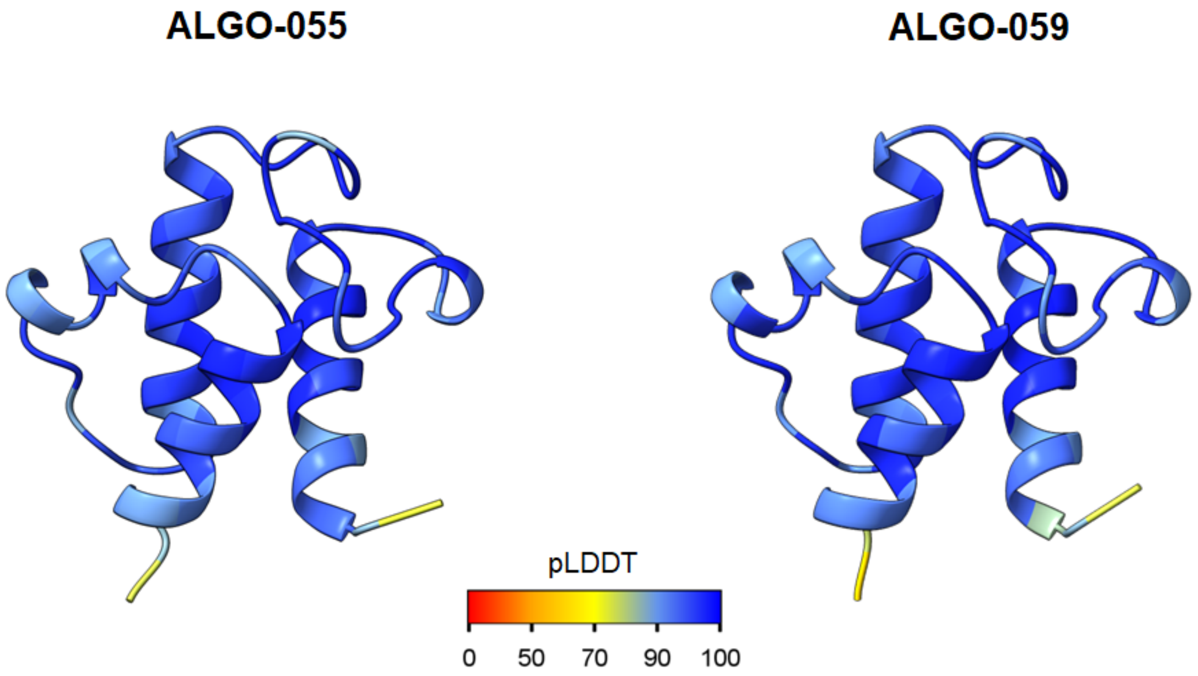


**Fig. S42** AlphaFold3 predicted structural models of ALGO-055 and ALGO-059.

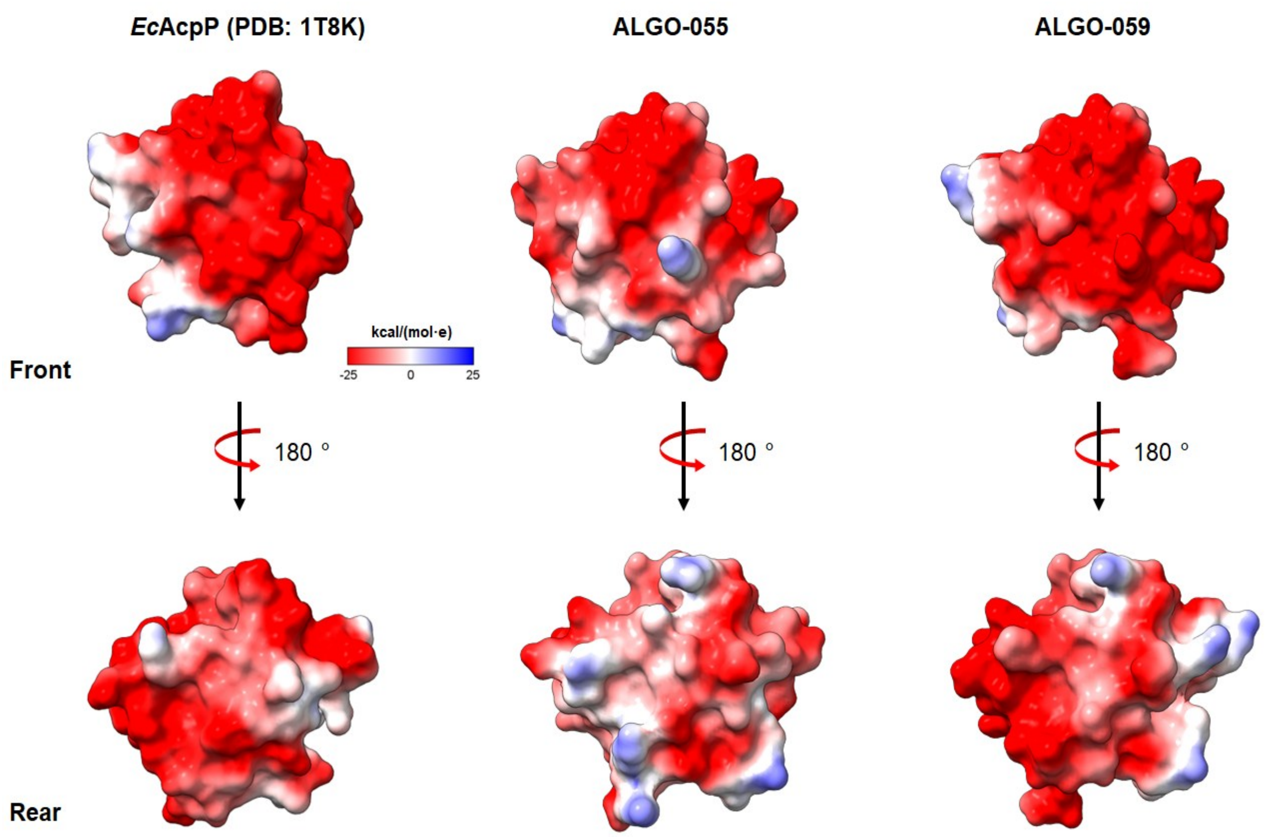


**Fig. S43** Coulombic potential maps of *Ec*AcpP, alongisde AlphaFold3 models of ALGO-055 and ALGO059.

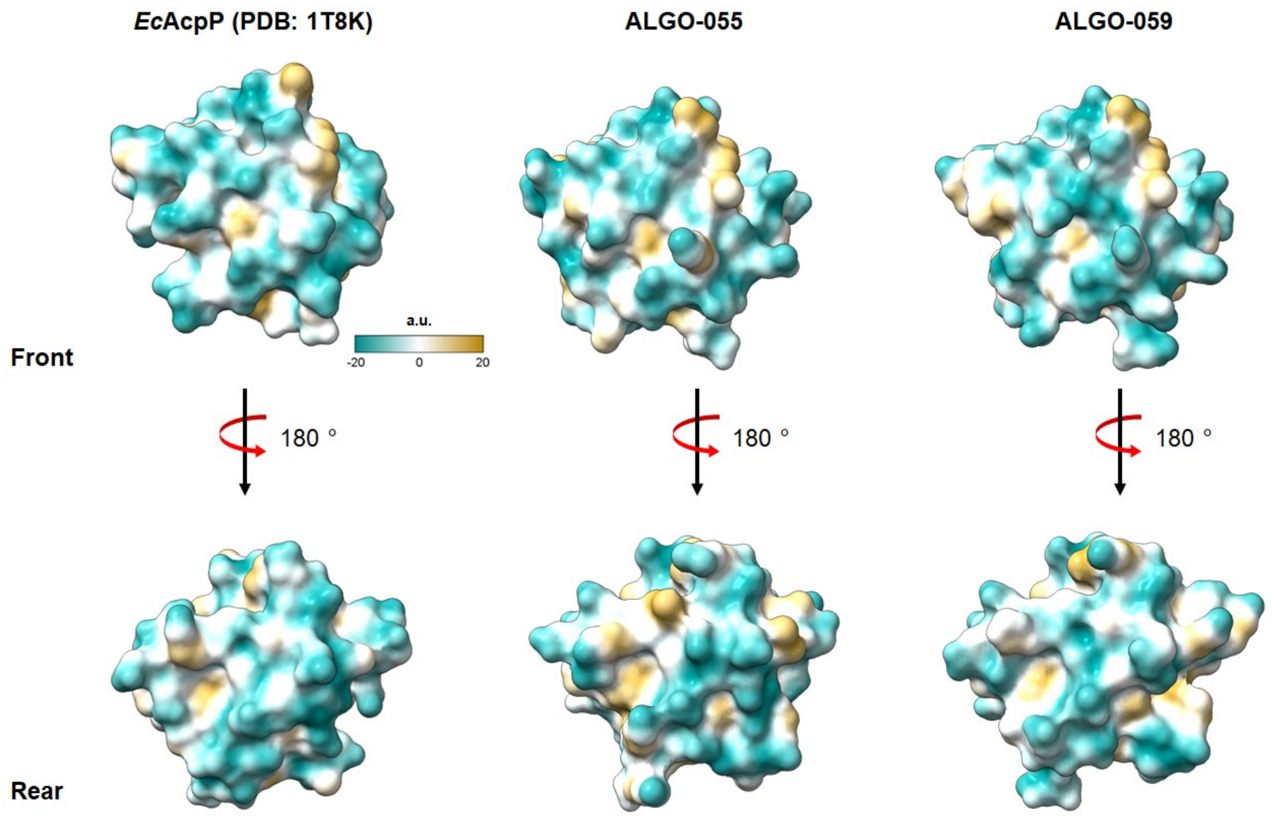


**Fig. S44** Molecular lipophilicity potential maps of *Ec*AcpP, alongisde AlphaFold3 models of ALGO-055 and ALGO059.

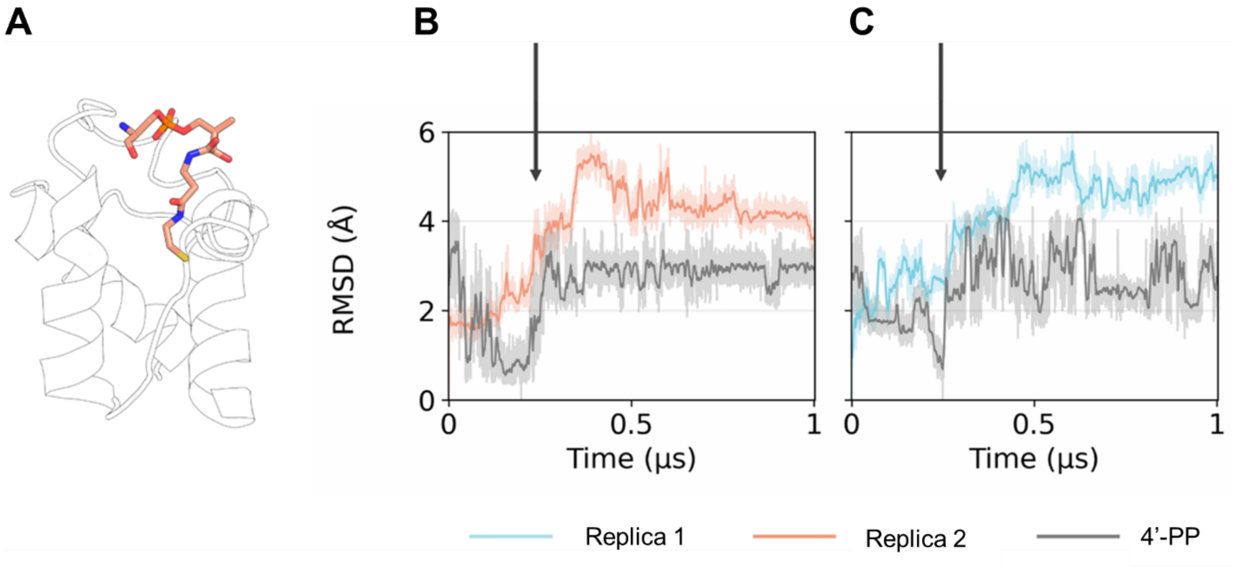


**Fig. S45** Correlation between 4’-PP and protein RMSD in selected *holo-* ACP replicas. Arrows indicate the point of 4’-PP displacement from the ACP binding pocket. (A) A representative frame extracted from MD simulation, showing how the 4’-PP group positions itself inside the binding pocket. (B) *holo-Ec*AcpP, Replica 2 (C) *holo*-ALGO-059, Replica 1.

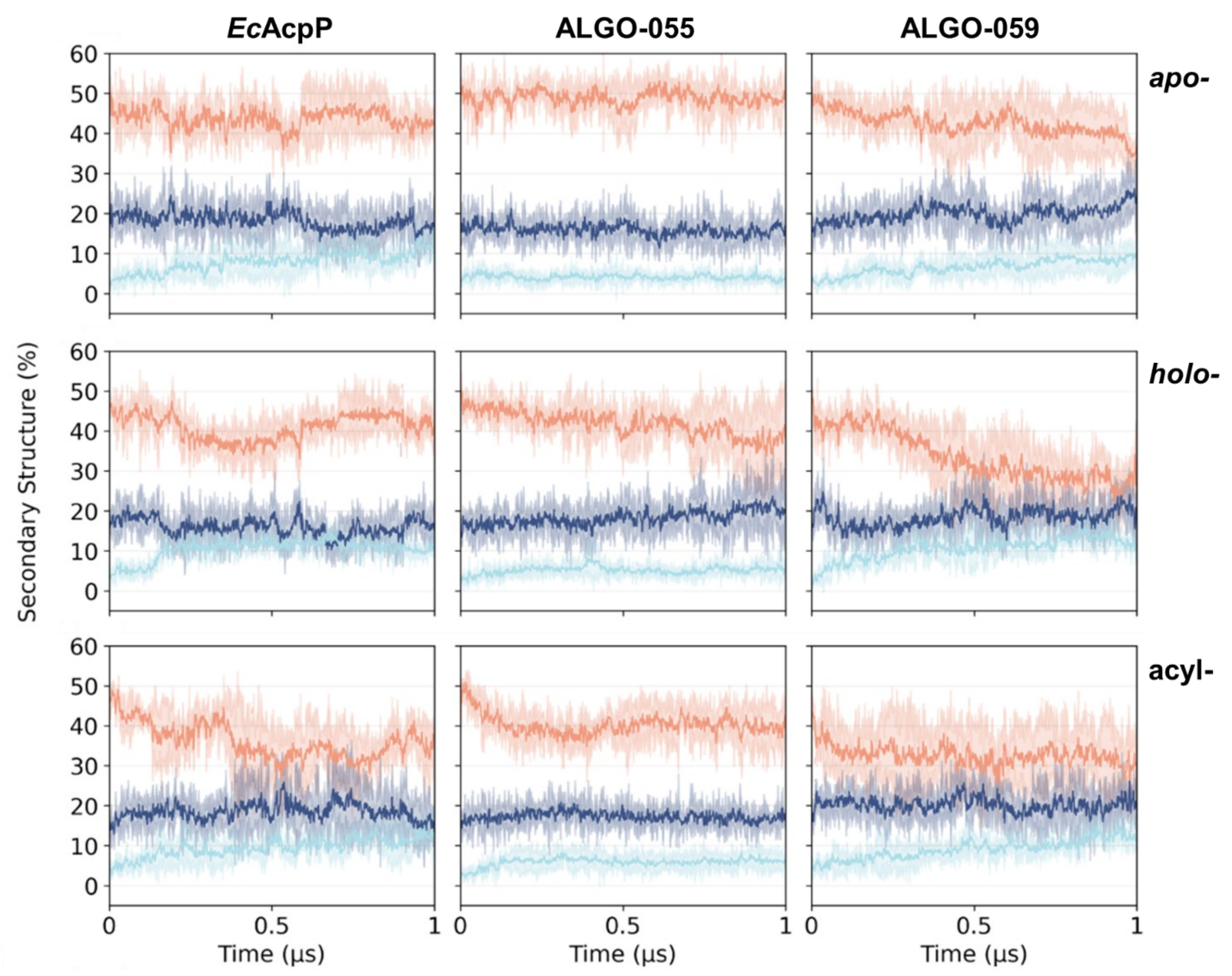


**Fig. S46** Backbone DSSP profiles of MD trajectories. MD trajectories of *Ec*AcpP, ALGO-055, and ALGO-059 models were analysed in *apo*- (top), *holo*- (middle) and $C_{12}$-acylated (bottom) forms.

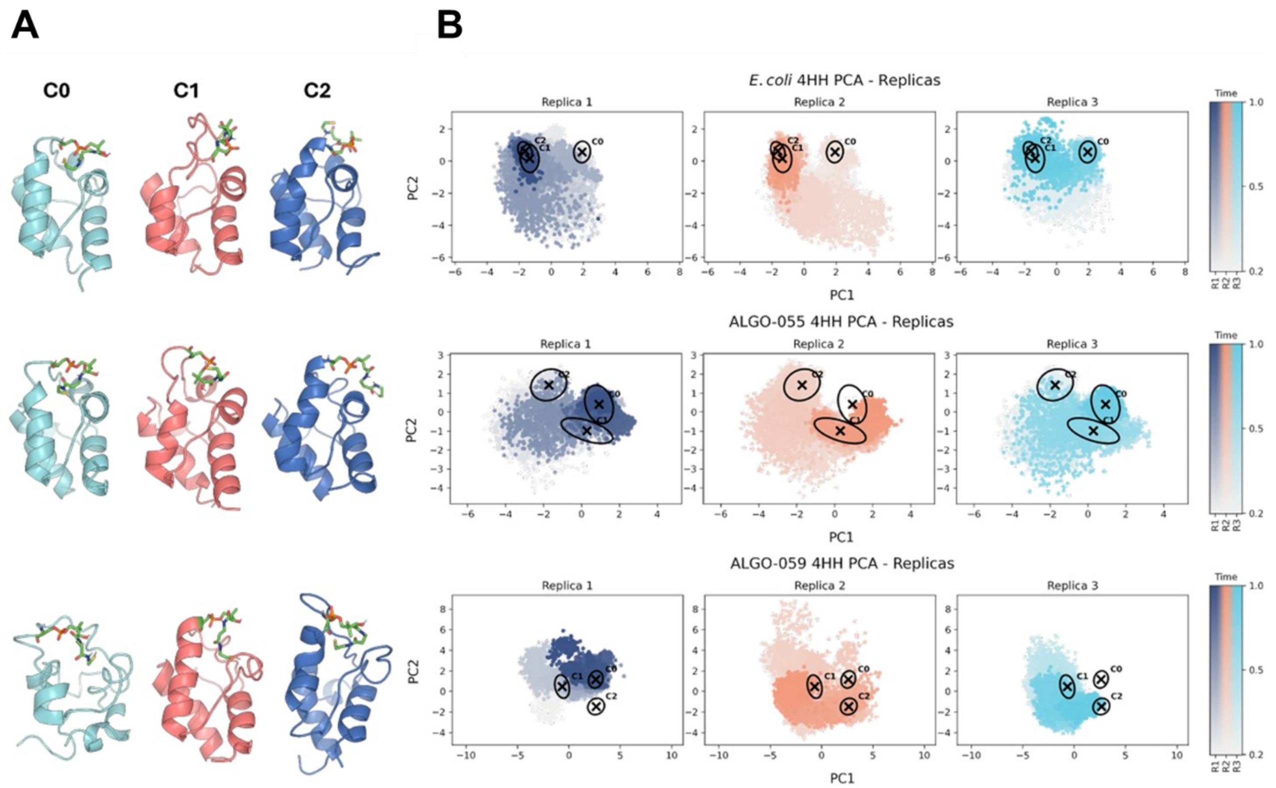


**Fig. S47** Cluster analysis and PCA projections of *holo*-ACP trajectories. (A) Representative structures of the three main centroids (C0, C1, C2), clusters are marked as black circles. The 4HH moiety is represented as sticks in green. (B) PCA projections onto the PC1-PC2 plane for each replicate of the four ACP variants. Trajectories evolution is represented by colormaps. Clusters are marked as black circles.

#### Additional CD data (Fig. S48)

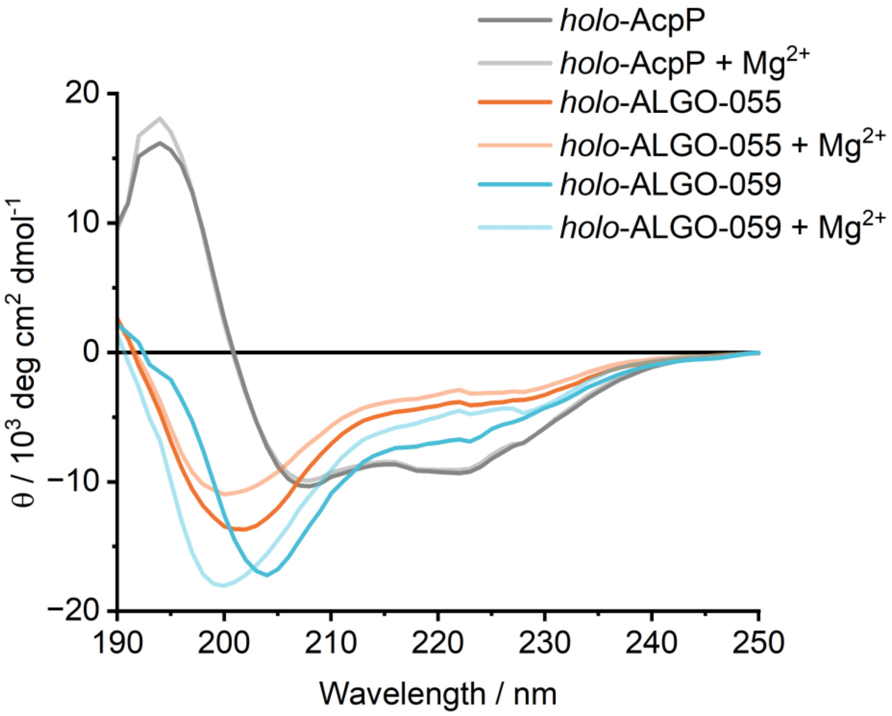


**Fig. S48** CD spectra of *holo*- proteins before and after $MgCl_2$ supplementation.

#### Additional sequence data (Fig. S49)

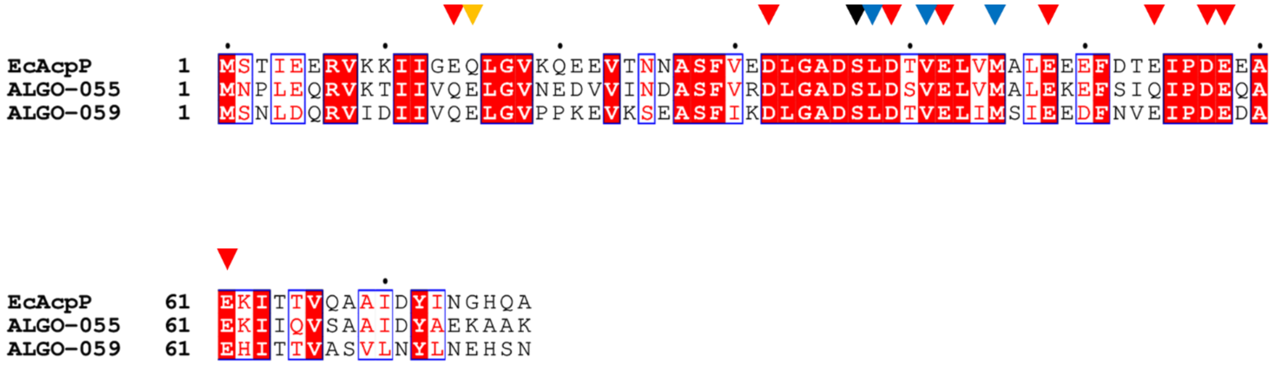


**Fig. S49** MSA of *Ec*AcpP, ALGO-055 and ALGO-059, highlighting conserved contact residues. *Ec*AcpP residues involved in PPIs with *Ec*AcpS and *Vh*AasS, based on reported crystallographic and NMR data, are shown as triangles. Black triangle (position 37) highlights the position of the invariant serine. Red triangles show important acidic contact residues. Blue triangles highlight known hydrophobic contact residues that interact with *Vh*AasS. Yellow triangle (position 15) highlights Q→E variation in ALGO-055 and ALGO-059 that could still plausibly engage in PPIs.

#### Raw SDS-PAGE Images

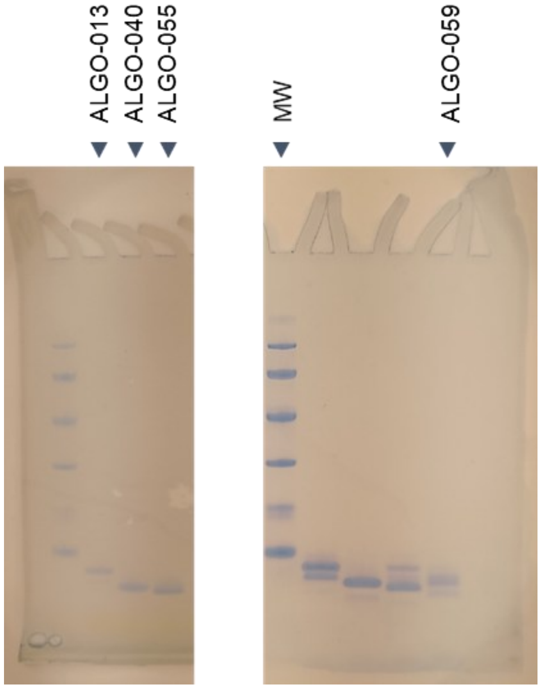


Raw SDS-PAGE gel photographs used for the composite image shown in Fig. 3B. Triangles indicate the lanes cropped.

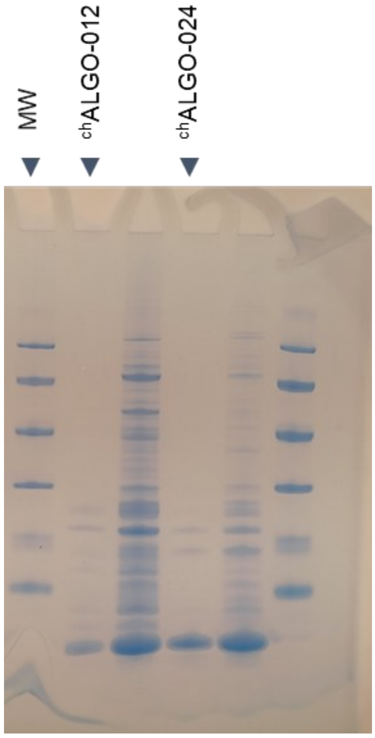


Raw SDS-PAGE gel photograph used for the image shown in Fig. 3G. Triangles indicate the lanes cropped.

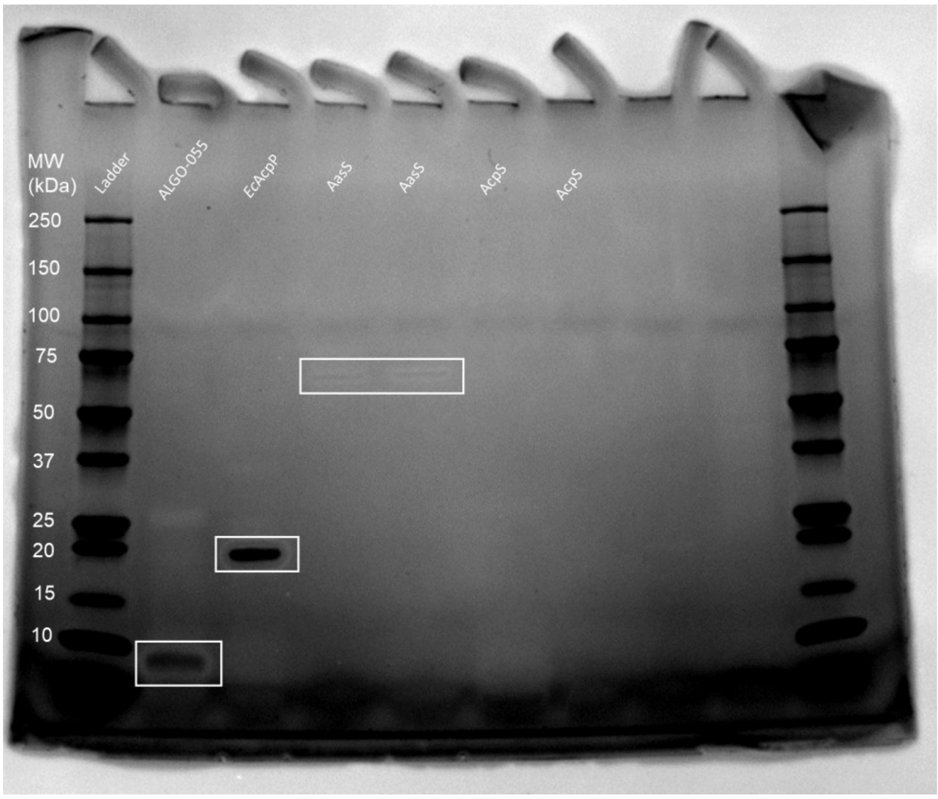

Raw SDS-PAGE gel photograph used for the image shown in Fig. S14A.

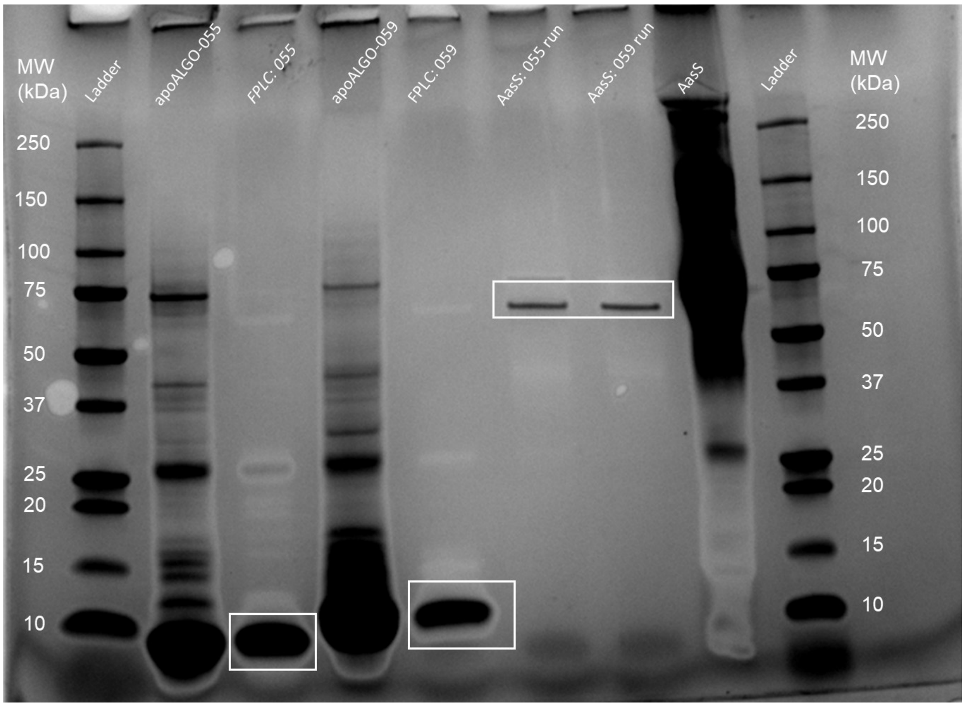

Raw SDS-PAGE gel photograph used for the image shown in Fig. S14B.

### Plasmid Maps

#### ALGO013

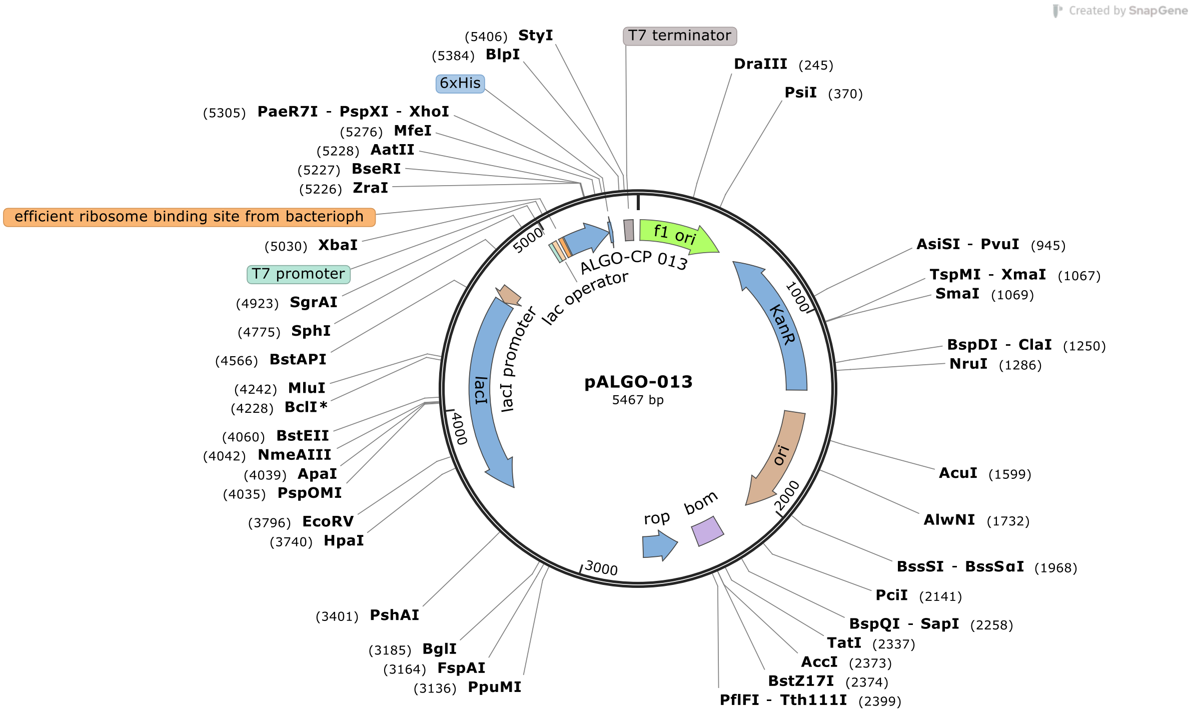

### pALGO-023

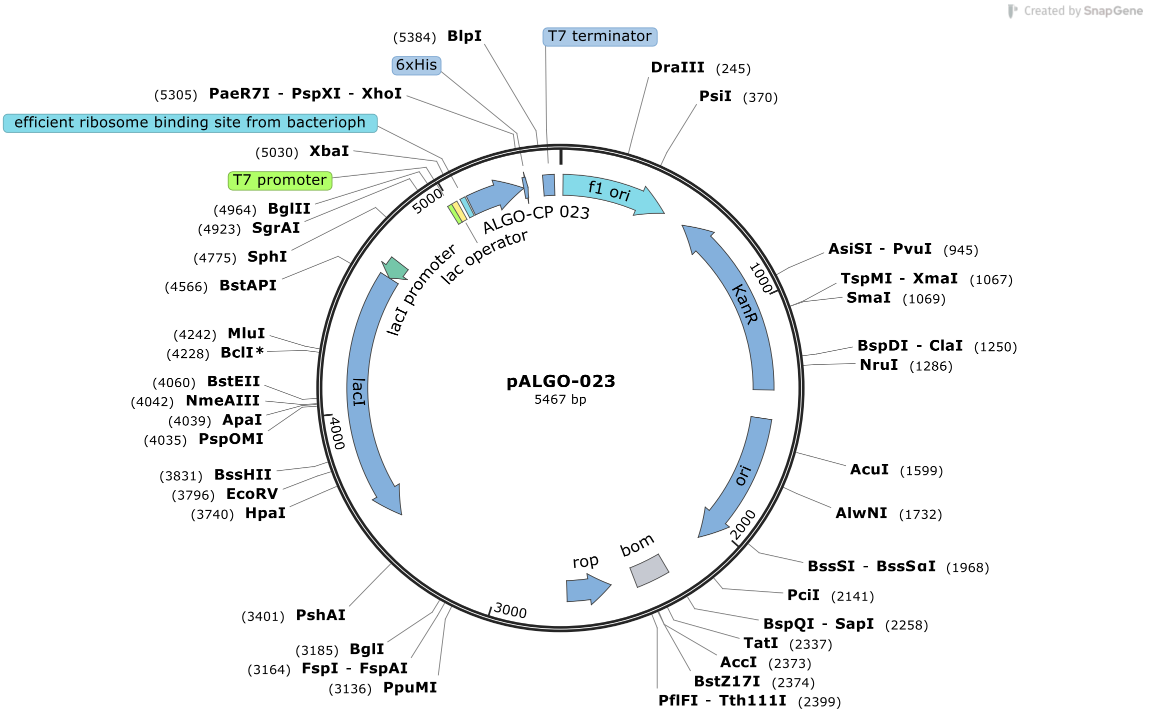

Created by SnapGene
pALGO-023
5467 bp
(5384) BlpI
T7 terminator
6xHis
(5305) PaeR7I - PspXI - XhoI
efficient ribosome binding site from bacterioph
(5030) XbaI
T7 promoter
(4964) BglII
(4923) SgrAI
(4775) SphI
(4566) BstAPI
(4242) MluI
(4228) BclI*
(4060) BstEII
(4042) NmeAIII
(4039) ApaI
(4035) PspOMI
(3831) BssHII
(3796) EcoRV
(3740) HpaI
(3401) PshAI
(3185) BglI
(3164) FspI - FspAI
(3136) PpuMI
DraIII (245)
PsiI (370)
AsiSI - PvuI (945)
TspMI - XmaI (1067)
SmaI (1069)
BspDI - ClaI (1250)
NruI (1286)
AcuI (1599)
AlwNI (1732)
BssSI - BssSαI (1968)
PciI (2141)
BspQI - SapI (2258)
TatI (2337)
AccI (2373)
BstZ17I (2374)
PflFI - Tth111I (2399)
f1 ori
KanR
ori
bom
rop
lacI
lacI promoter
lac operator
ALGO-CP 023
1000
2000
3000
4000
5000

#### pALGO-040

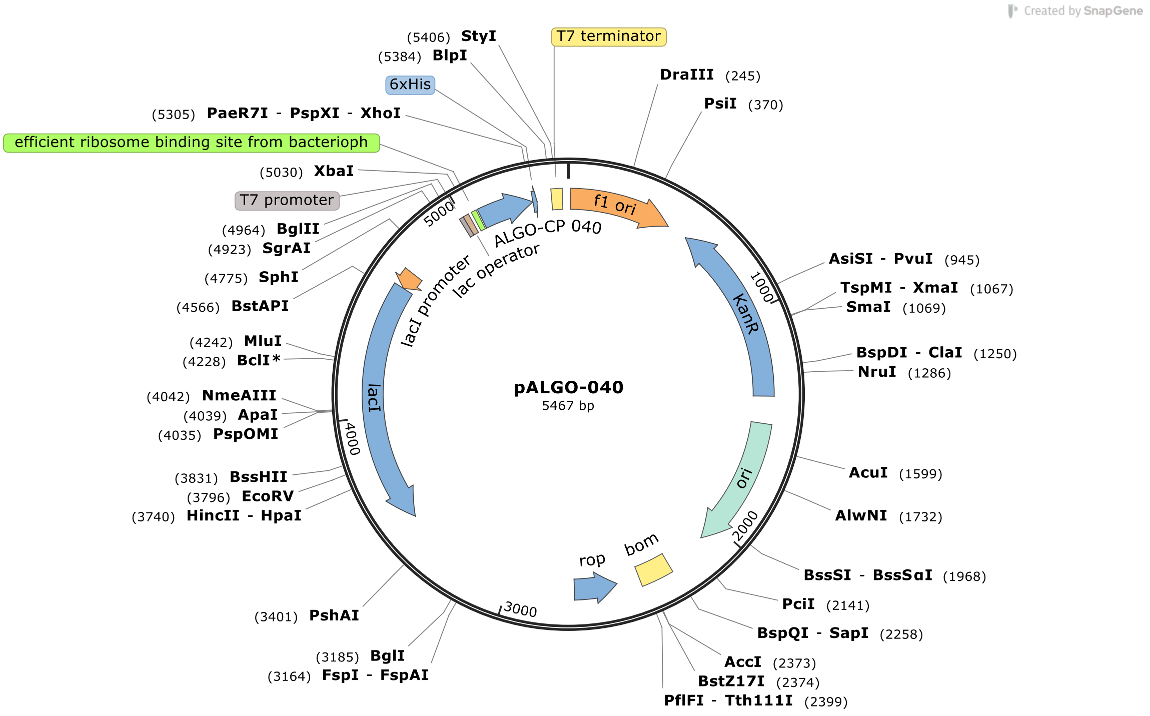

Created by SnapGene
(5406) StyI
(5384) BlpI
6xHis
(5305) PaeR7I - PspXI - XhoI
efficient ribosome binding site from bacterioph
(5030) XbaI
T7 promoter
(4964) BglII
(4923) SgrAI
(4775) SphI
(4566) BstAPI
(4242) MluI
(4228) BclI*
(4042) NmeAIII
(4039) ApaI
(4035) PspOMI
(3831) BssHII
(3796) EcoRV
(3740) HincII - HpaI
(3401) PshAI
(3185) BglI
(3164) FspI - FspAI
T7 terminator
DraIII (245)
PsiI (370)
AsiSI - PvuI (945)
TspMI - XmaI (1067)
SmaI (1069)
BspDI - ClaI (1250)
NruI (1286)
AcuI (1599)
AlwNI (1732)
BssSI - BssSαI (1968)
PciI (2141)
BspQI - SapI (2258)
AccI (2373)
BstZ17I (2374)
PflFI - Tth111I (2399)
f1 ori
ALGO-CP 040
lac operator
lacI promoter
lacI
KanR
ori
rop
bom
1000
2000
3000
4000
5000
pALGO-040
5467 bp

#### ppALGO-044

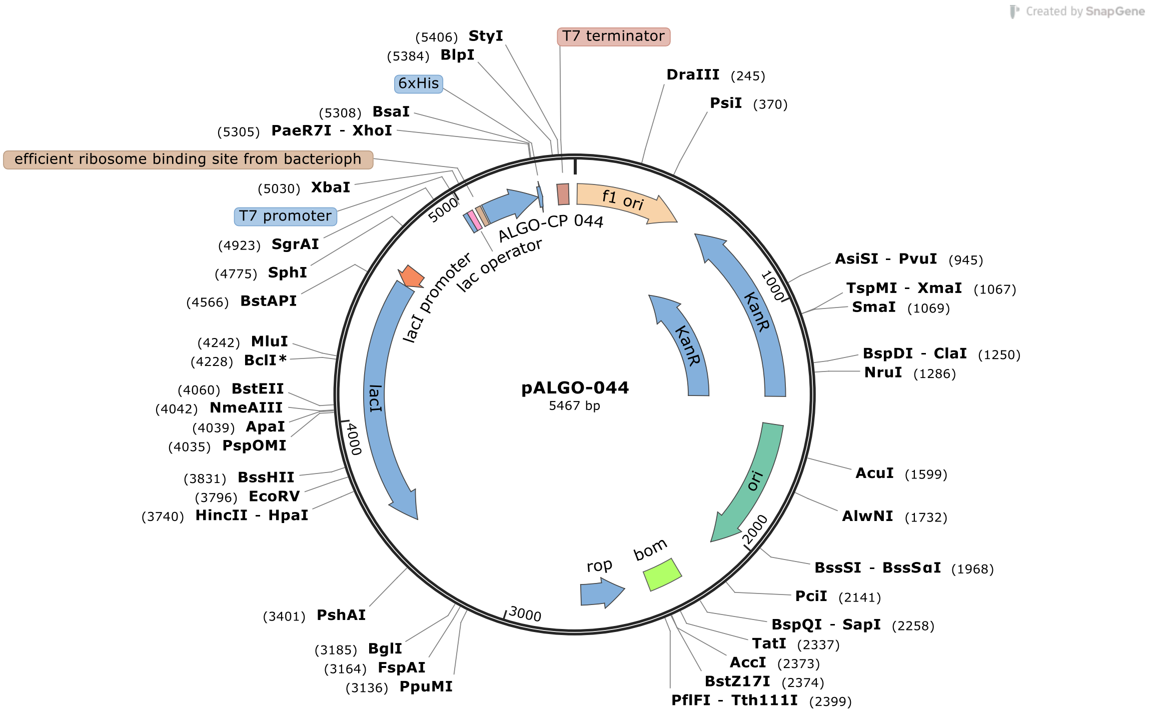

Created by SnapGene
pALGO-044
5467 bp
T7 terminator
f1 ori
KanR
KanR
ori
bom
rop
lacI
lacI promoter
lac operator
ALGO-CP 044
6xHis
efficient ribosome binding site from bacterioph
T7 promoter
1000
2000
3000
4000
5000
(5406) StyI
(5384) BlpI
(5308) BsaI
(5305) PaeR7I - XhoI
(5030) XbaI
(4923) SgrAI
(4775) SphI
(4566) BstAPI
(4242) MluI
(4228) BclI*
(4060) BstEII
(4042) NmeAIII
(4039) ApaI
(4035) PspOMI
(3831) BssHII
(3796) EcoRV
(3740) HincII - HpaI
(3401) PshAI
(3185) BglI
(3164) FspAI
(3136) PpuMI
DraIII (245)
PsiI (370)
AsiSI - PvuI (945)
TspMI - XmaI (1067)
SmaI (1069)
BspDI - ClaI (1250)
NruI (1286)
AcuI (1599)
AlwNI (1732)
BssSI - BssSαI (1968)
PciI (2141)
BspQI - SapI (2258)
TatI (2337)
AccI (2373)
BstZ17I (2374)
PflFI - Tth111I (2399)

#### pALGO-055

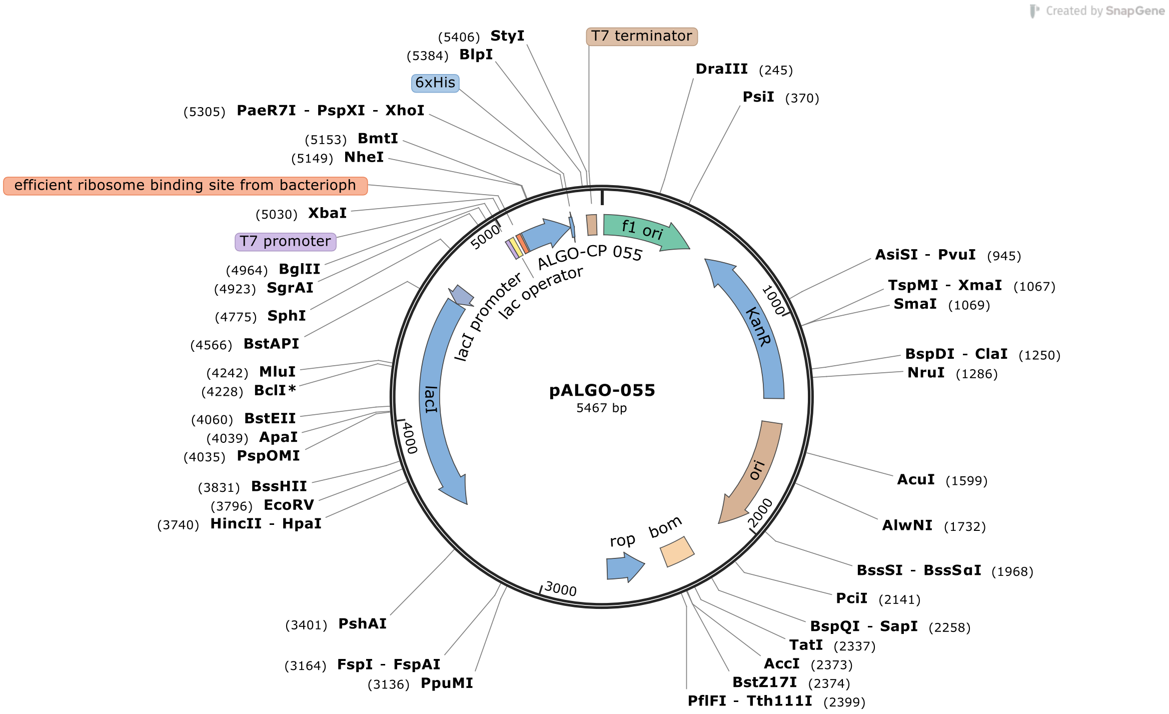
Created by SnapGene
pALGO-055
5467 bp
(5406) StyI
(5384) BlpI
6xHis
(5305) PaeR7I - PspXI - XhoI
(5153) BmtI
(5149) NheI
efficient ribosome binding site from bacterioph
(5030) XbaI
T7 promoter
(4964) BglII
(4923) SgrAI
(4775) SphI
(4566) BstAPI
(4242) MluI
(4228) BclI*
(4060) BstEII
(4039) ApaI
(4035) PspOMI
(3831) BssHII
(3796) EcoRV
(3740) HincII - HpaI
(3401) PshAI
(3164) FspI - FspAI
(3136) PpuMI
T7 terminator
DraIII (245)
PsiI (370)
AsiSI - PvuI (945)
TspMI - XmaI (1067)
SmaI (1069)
BspDI - ClaI (1250)
NruI (1286)
AcuI (1599)
AlwNI (1732)
BssSI - BssSαI (1968)
PciI (2141)
BspQI - SapI (2258)
TatI (2337)
AccI (2373)
BstZ17I (2374)
PflFI - Tth111I (2399)
f1 ori
KanR
ori
bom
rop
lacI
lacI promoter
lac operator
ALGO-CP 055
1000
2000
3000
4000
5000

#### pALGO-057

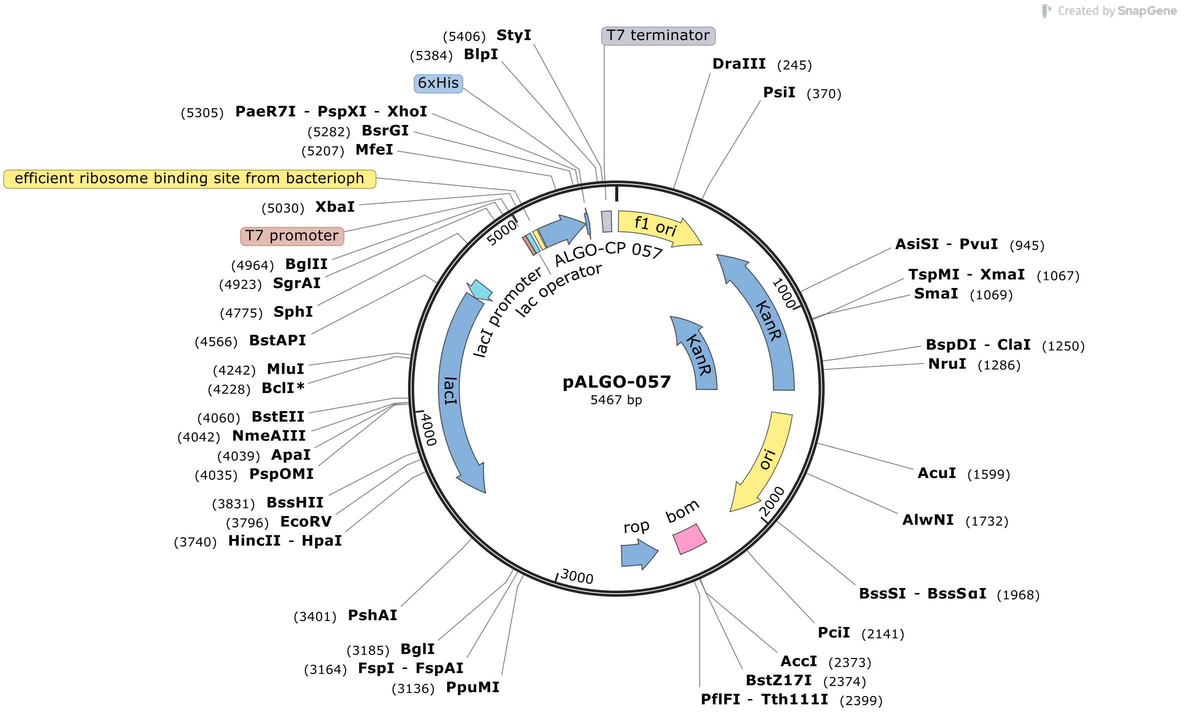
Created by SnapGene
(5406) StyI
(5384) BlpI
6xHis
(5305) PaeR7I - PspXI - XhoI
(5282) BsrGI
(5207) MfeI
efficient ribosome binding site from bacterioph
(5030) XbaI
T7 promoter
(4964) BglII
(4923) SgrAI
(4775) SphI
(4566) BstAPI
(4242) MluI
(4228) BclI*
(4060) BstEII
(4042) NmeAIII
(4039) ApaI
(4035) PspOMI
(3831) BssHII
(3796) EcoRV
(3740) HincII - HpaI
(3401) PshAI
(3185) BglI
(3164) FspI - FspAI
(3136) PpuMI
T7 terminator
DraIII (245)
PsiI (370)
AsiSI - PvuI (945)
TspMI - XmaI (1067)
SmaI (1069)
BspDI - ClaI (1250)
NruI (1286)
AcuI (1599)
AlwNI (1732)
BssSI - BssSαI (1968)
PciI (2141)
AccI (2373)
BstZ17I (2374)
PflFI - Tth111I (2399)
f1 ori
ALGO-CP 057
lac operator
lacI promoter
KanR
KanR
lacI
ori
rop
bom
5000
1000
2000
3000
4000
pALGO-057
5467 bp

**pALGO-059**

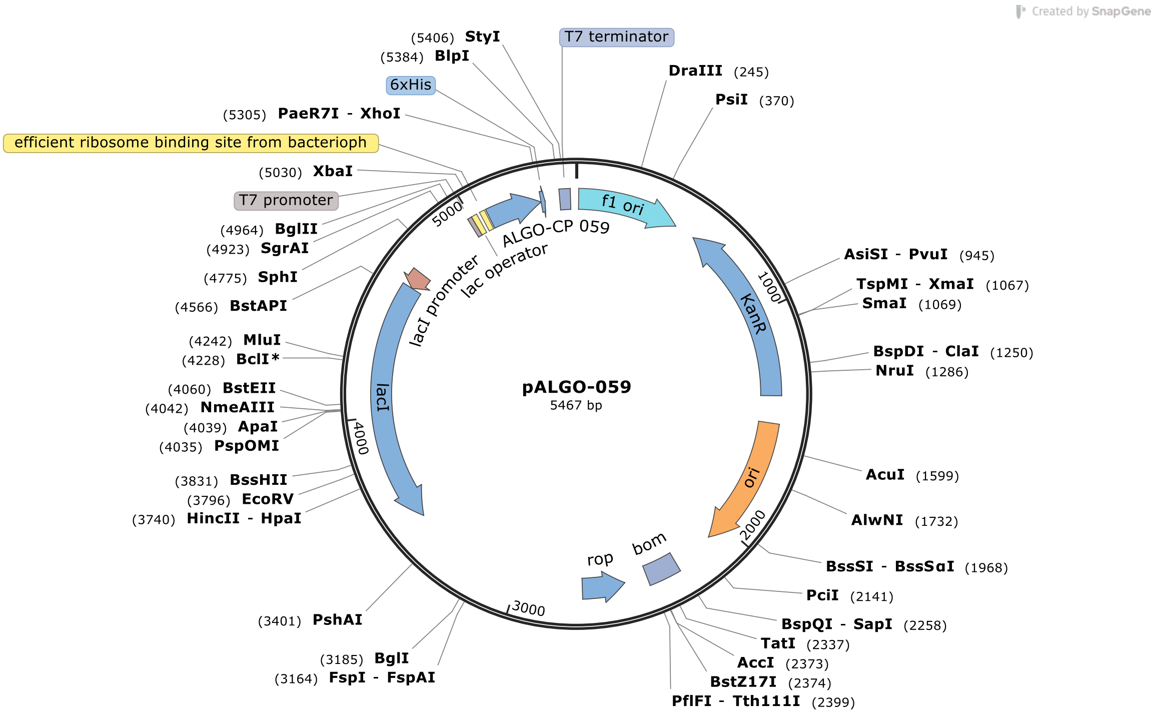

Created by SnapGene
(5406) StyI
(5384) BlpI
T7 terminator
DraIII (245)
PsiI (370)
6xHis
(5305) PaeR7I - XhoI
efficient ribosome binding site from bacterioph
(5030) XbaI
T7 promoter
(4964) BglII
(4923) SgrAI
(4775) SphI
(4566) BstAPI
(4242) MluI
(4228) BclI*
(4060) BstEII
(4042) NmeAIII
(4039) ApaI
(4035) PspOMI
(3831) BssHII
(3796) EcoRV
(3740) HincII - HpaI
(3401) PshAI
(3185) BglI
(3164) FspI - FspAI
AsiSI - PvuI (945)
TspMI - XmaI (1067)
SmaI (1069)
BspDI - ClaI (1250)
NruI (1286)
AcuI (1599)
AlwNI (1732)
BssSI - BssSαI (1968)
PciI (2141)
BspQI - SapI (2258)
TatI (2337)
AccI (2373)
BstZ17I (2374)
PflFI - Tth111I (2399)
f1 ori
ALGO-CP 059
lac operator
lacI promoter
lacI
KanR
ori
rop
bom
1000
2000
3000
4000
5000
pALGO-059
5467 bp

pCHALGO-009

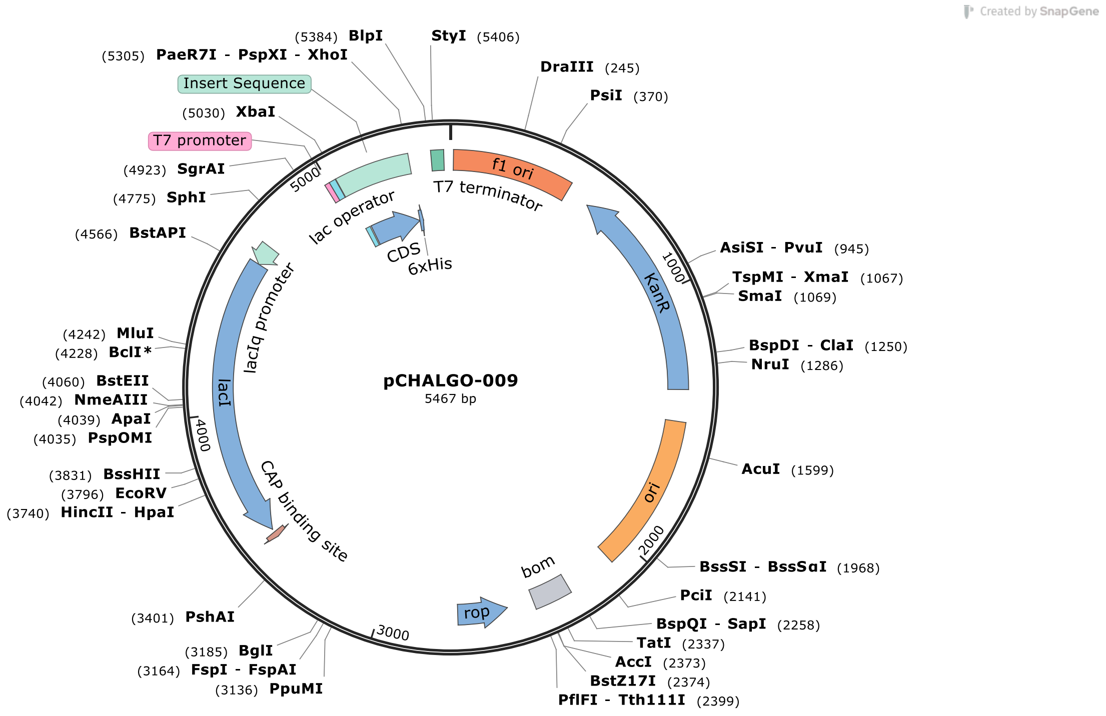

Created by SnapGene
(5384) BlpI
StyI (5406)
(5305) PaeR7I - PspXI - XhoI
DraIII (245)
Insert Sequence
PsiI (370)
(5030) XbaI
T7 promoter
(4923) SgrAI
(4775) SphI
(4566) BstAPI
AsiSI - PvuI (945)
TspMI - XmaI (1067)
SmaI (1069)
BspDI - ClaI (1250)
NruI (1286)
(4242) MluI
(4228) BclI*
(4060) BstEII
(4042) NmeAIII
(4039) ApaI
(4035) PspOMI
(3831) BssHII
(3796) EcoRV
(3740) HincII - HpaI
AcuI (1599)
BssSI - BssSαI (1968)
PciI (2141)
BspQI - SapI (2258)
TatI (2337)
AccI (2373)
BstZ17I (2374)
PflFI - Tth111I (2399)
(3401) PshAI
(3185) BglI
(3164) FspI - FspAI
(3136) PpuMI
f1 ori
T7 terminator
lac operator
CDS
6xHis
KanR
lacIq promoter
lacI
CAP binding site
ori
bom
rop
5000
1000
4000
2000
3000
pCHALGO-009
5467 bp

### pCHALGO-012

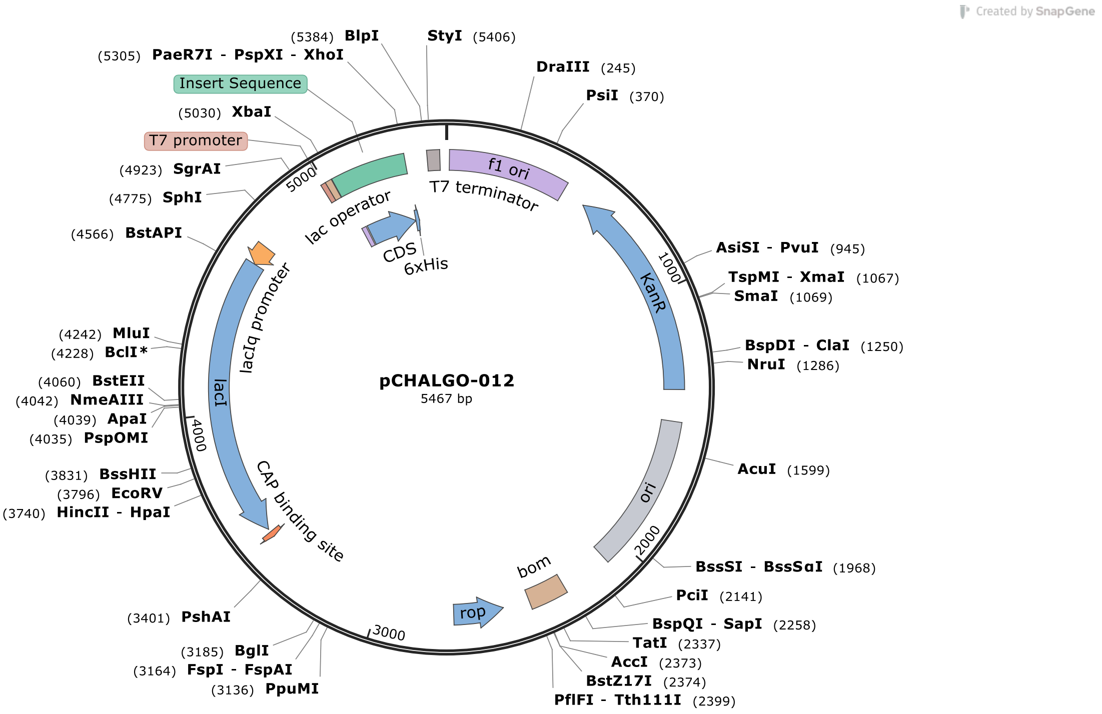
Created by SnapGene
(5384) BlpI
StyI (5406)
(5305) PaeR7I - PspXI - XhoI
DraIII (245)
Insert Sequence
PsiI (370)
(5030) XbaI
T7 promoter
(4923) SgrAI
(4775) SphI
(4566) BstAPI
AsiSI - PvuI (945)
TspMI - XmaI (1067)
SmaI (1069)
(4242) MluI
(4228) BclI*
BspDI - ClaI (1250)
NruI (1286)
(4060) BstEII
(4042) NmeAIII
(4039) ApaI
(4035) PspOMI
AcuI (1599)
(3831) BssHII
(3796) EcoRV
(3740) HincII - HpaI
BssSI - BssSαI (1968)
PciI (2141)
(3401) PshAI
BspQI - SapI (2258)
TatI (2337)
(3185) BglI
AccI (2373)
(3164) FspI - FspAI
BstZ17I (2374)
(3136) PpuMI
PflFI - Tth111I (2399)
f1 ori
T7 terminator
lac operator
CDS
6xHis
lacIq promoter
lacI
CAP binding site
KanR
ori
bom
rop
pCHALGO-012
5467 bp
1000
2000
3000
4000
5000

pCHALGO-024

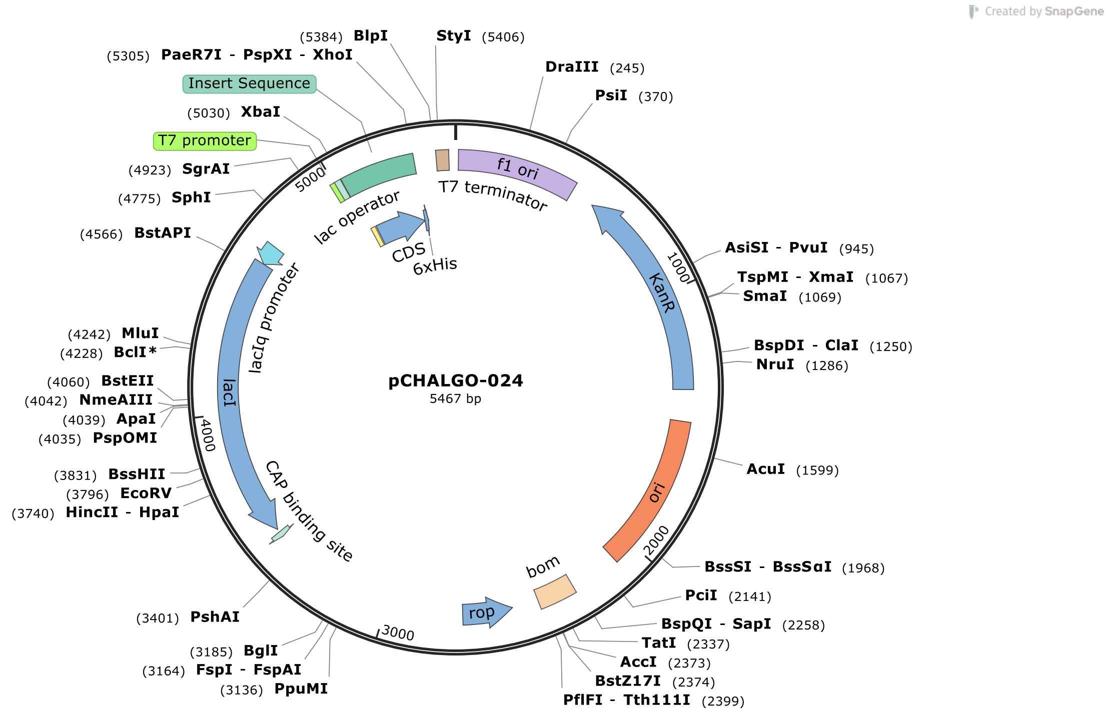
Created by SnapGene
pCHALGO-024
5467 bp
(5384) BlpI
StyI (5406)
(5305) PaeR7I - PspXI - XhoI
DraIII (245)
PsiI (370)
Insert Sequence
(5030) XbaI
T7 promoter
(4923) SgrAI
(4775) SphI
(4566) BstAPI
AsiSI - PvuI (945)
TspMI - XmaI (1067)
SmaI (1069)
BspDI - ClaI (1250)
NruI (1286)
(4242) MluI
(4228) BclI*
(4060) BstEII
(4042) NmeAIII
(4039) ApaI
(4035) PspOMI
AcuI (1599)
(3831) BssHII
(3796) EcoRV
(3740) HincII - HpaI
BssSI - BssSαI (1968)
PciI (2141)
(3401) PshAI
BspQI - SapI (2258)
TatI (2337)
AccI (2373)
(3185) BglI
BstZ17I (2374)
(3164) FspI - FspAI
PflFI - Tth111I (2399)
(3136) PpuMI
f1 ori
T7 terminator
lac operator
CDS
6xHis
lacIq promoter
KanR
lacI
ori
CAP binding site
bom
rop
1000
2000
3000
4000
5000

pCHALGO-044

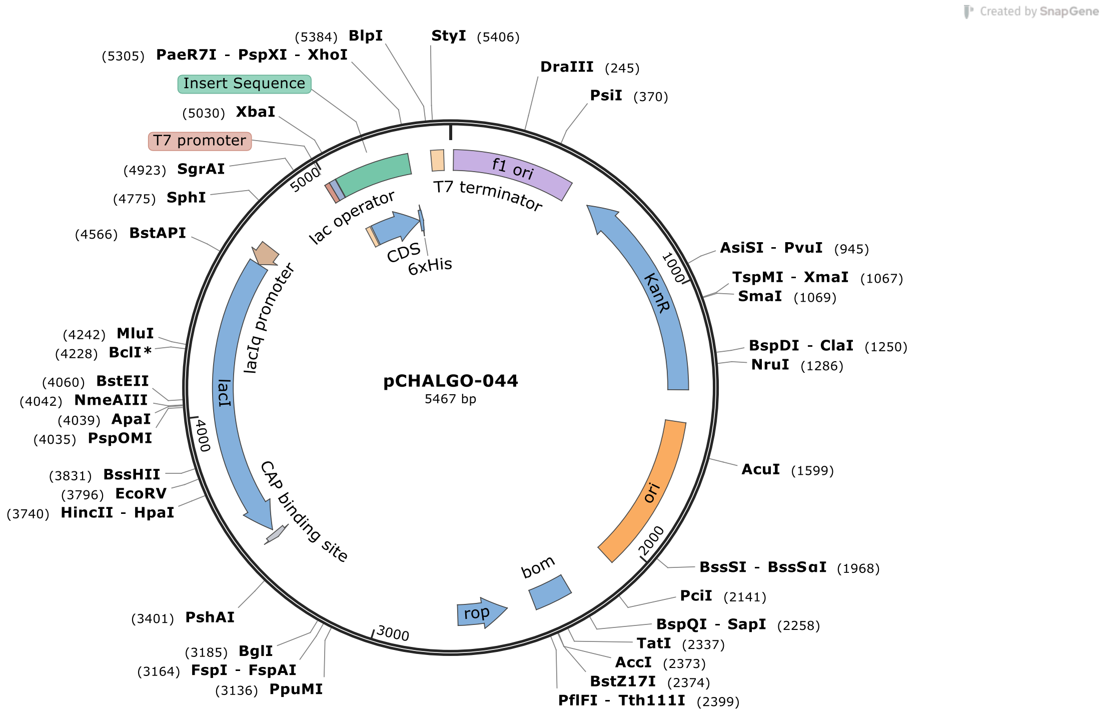
Created by SnapGene
(5384) BlpI
StyI (5406)
(5305) PaeR7I - PspXI - XhoI
DraIII (245)
PsiI (370)
Insert Sequence
(5030) XbaI
T7 promoter
(4923) SgrAI
(4775) SphI
(4566) BstAPI
f1 ori
T7 terminator
lac operator
CDS
6xHis
lacIq promoter
AsiSI - PvuI (945)
TspMI - XmaI (1067)
SmaI (1069)
KanR
BspDI - ClaI (1250)
NruI (1286)
(4242) MluI
(4228) BclI*
(4060) BstEII
(4042) NmeAIII
(4039) ApaI
(4035) PspOMI
lacI
pCHALGO-044
5467 bp
AcuI (1599)
ori
(3831) BssHII
(3796) EcoRV
(3740) HincII - HpaI
CAP binding site
BssSI - BssSαI (1968)
PciI (2141)
bom
rop
(3401) PshAI
BspQI - SapI (2258)
TatI (2337)
AccI (2373)
BstZ17I (2374)
PflFI - Tth111I (2399)
(3185) BglI
(3164) FspI - FspAI
(3136) PpuMI
1000
2000
3000
4000
5000

pCHALGO-097

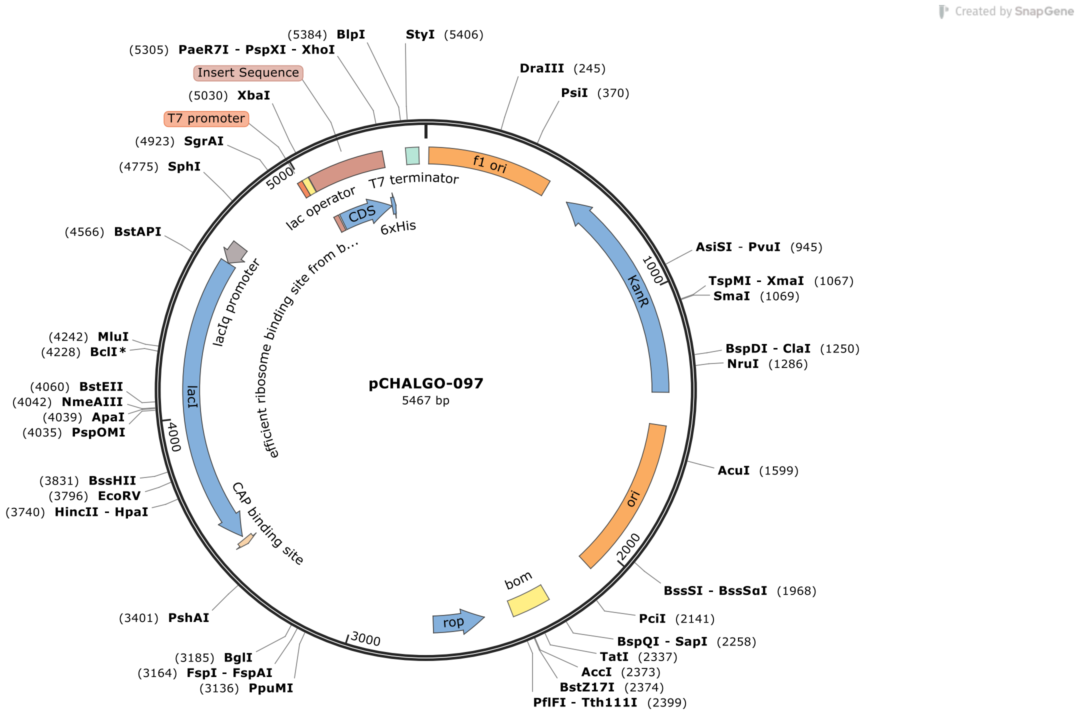
Created by SnapGene
(5384) BlpI
StyI (5406)
(5305) PaeR7I - PspXI - XhoI
Insert Sequence
DraIII (245)
PsiI (370)
(5030) XbaI
T7 promoter
(4923) SgrAI
(4775) SphI
(4566) BstAPI
AsiSI - PvuI (945)
TspMI - XmaI (1067)
SmaI (1069)
(4242) MluI
(4228) BclI*
BspDI - ClaI (1250)
NruI (1286)
(4060) BstEII
(4042) NmeAIII
(4039) ApaI
(4035) PspOMI
(3831) BssHII
(3796) EcoRV
(3740) HincII - HpaI
AcuI (1599)
BssSI - BssSαI (1968)
(3401) PshAI
PciI (2141)
BspQI - SapI (2258)
TatI (2337)
AccI (2373)
BstZ17I (2374)
PflFI - Tth111I (2399)
(3185) BglI
(3164) FspI - FspAI
(3136) PpuMI
T7 terminator
f1 ori
lac operator
CDS
6xHis
efficient ribosome binding site from b...
lacIq promoter
lacI
KanR
ori
bom
rop
CAP binding site
1000
2000
3000
4000
5000
pCHALGO-097
5467 bp
